# Myotonic Muscle Activity Drives a Persistent Pain State Responsive to Na_V_1.8-directed Analgesics

**DOI:** 10.1101/2024.06.19.599732

**Authors:** Tyler S. Nelson, Aida Calderon-Rivera, Heather N. Allen, Alaina L. Waters, Emanuel Loeza-Alcocer, Jorge B. Pineda-Farias, Narges Pachenari, Santiago Loya-Lopez, Kimberly Gomez, Erick J. Rodriguez-Palma, Paz Duran, Michael S. Gold, Rajesh Khanna

**Author notes:** **Corresponding author:** Name: Rajesh Khanna, Ph.D., Mailing Address: 1149 Newell Drive, Room L4-162, Gainesville, FL 32610, USA.

## Abstract

Pain is a common and disabling feature of myotonic disorders, yet its biological basis remains poorly understood and no targeted analgesic therapies currently exist. Here, we demonstrate that skeletal muscle hyperexcitability is sufficient to initiate a persistent pain state independent of inflammation, nerve injury, or overt tissue damage. Using complementary pharmacological and genetic models of myotonia resulting from loss of the voltage-gated skeletal muscle chloride channel ClC-1 function, we show that transient and chronic myotonia produce robust mechanical, thermal, and cold hypersensitivity, as well as spontaneous pain-like behavior. Notably, pain-like behaviors induced by transient myotonia persist long after overt motor symptoms have resolved, suggesting that a transient episode of muscle hyperexcitability is sufficient to trigger prolonged alterations in nociceptive processing. Physiological recordings revealed altered excitability of dorsal root ganglion and superficial dorsal horn neurons and enhanced sensory-evoked activity in the parabrachial nucleus, indicating altered nociceptive processing across multiple levels of the pain neuraxis. Transient myotonia increased total sodium current density in sensory neurons, with a shift toward a greater tetrodotoxin-resistant current fraction. Pharmacological inhibition with the Na_V_1.8-directed analgesic Suzetrigine markedly attenuated pain-like behaviors in both models of myotonia. Together, these findings establish a link between myotonia and persistent alterations in nociceptive processing and identify Na_V_1.8-directed analgesia as a promising therapeutic strategy for myotonia-associated pain.

## Introduction

Myotonic disorders are inherited neuromuscular conditions characterized by delayed muscle relaxation following voluntary contraction (1, 2). This defining feature, known as myotonia, arises from increased skeletal muscle excitability and is most commonly caused by mutations in genes *CLCN1* and *SCN4A*, which encode the skeletal muscle voltage-gated chloride channel ClC-1 and the voltage-gated sodium channel Na_V_1.4, respectively (3, 4). Clinically, myotonia may manifest as overt muscle stiffness or as more subtle impairments in muscle relaxation detectable only through electromyography (EMG) (3, 5, 6). These symptoms frequently interfere with motor function, elevate fall risk, and substantially reduce quality of life (7–9).

ClC-1 plays a central role in stabilizing skeletal muscle excitability by facilitating chloride ion conductance across the sarcolemma (10). Loss-of-function mutations in *CLCN1* impair chloride conductance, resulting in sustained depolarization and delayed repolarization of muscle fibers (11). This pathomechanism underlies both autosomal dominant (Thomsen’s disease) and autosomal recessive (Becker’s disease) forms of myotonia congenita (12–15). A similar mechanism contributes to myotonia in myotonic dystrophy type 1 (DM1), where mis-splicing of *CLCN1* mRNA leads to reduced ClC-1 expression and subsequent muscle hyperexcitability (16–18).

To investigate myotonia under experimental conditions, researchers have long employed monocarboxylic aromatic acids, which reliably induce muscle stiffness and characteristic EMG abnormalities in mammalian and amphibian skeletal muscle (19–27). Among these compounds, anthracene-9-carboxylic acid (9-AC) is one of the most widely used and well-characterized agents. By selectively blocking CLC-1 channels (21, 28–30), 9-AC rapidly induces myotonic symptoms across multiple species, including rodents (31–34), rabbits (30), goats (21), dogs (35), pigs (36), and human skeletal muscle (37, 38). In rodents, systemic administration of 9-AC produces robust, transient muscle hyperexcitability and has become a standard pharmacological tool for evaluating candidate anti-myotonic therapies *in vivo* (31, 32, 39–41).

Although the electrophysiological basis of myotonia has been extensively studied, its contribution to the pathophysiology of pain remains poorly understood. Historically, myotonic disorders have been conceptualized as skeletal muscle channelopathies, yet pain has long been described as a prominent and distressing symptom. Early accounts from the late 19th century document severe, recurring muscle pain in affected individuals. In an 1886 article, Dr. White described a patient whose “cramps caused great pain, and came on several times a day, and lasted a long while (42).” Dana (1888) reported that patients “complain bitterly at times of aching pains in the legs (43),” while Deligny (1885) observed muscle contractions “always accompanied with such violent pain that the patient would roll on the earth in his agony (44, 45).” These descriptions illustrate that pain has historically been recognized as a consistent and debilitating feature of the myotonic phenotype. Contemporary studies reinforce these observations (46, 47). Patients with myotonia congenita (8, 48–53), myotonic dystrophies (DM1 and DM2) (54–61), and *SCN4A*-related myotonic disorders (46, 62–67) frequently report high-impact, treatment-resistant muscle pain that substantially limits daily functioning and is often the primary motivation for seeking medical care (7, 46, 47, 56, 57, 68–71). However, despite its prevalence and clinical relevance, the mechanistic basis of pain in myotonic disorders remains poorly understood, and no targeted analgesic treatments currently exist.

Given this gap in knowledge and the historical and clinical evidence of pain in myotonic disorders, we hypothesized that skeletal muscle hyperexcitability itself may be sufficient to initiate persistent pain states. To test this hypothesis, we employed complementary pharmacological and genetic models of myotonia resulting from loss of ClC-1 function. We show that a single episode of 9-AC-induced myotonia produces prolonged mechanical, thermal, and cold hypersensitivity that persists well beyond resolution of motor symptoms and is accompanied by increased excitability of peripheral sensory neurons and amplification of central nociceptive circuits. Furthermore, mice with genetically induced myotonia congenita exhibit persistent evoked and spontaneous pain-like behaviors that are similarly associated with sensory neuron hyperexcitability. Finally, we demonstrate that pharmacological inhibition with the Na_V_1.8-directed analgesic Suzetrigine markedly attenuates pain-like behavior in both pharmacological and genetic models of myotonia. Together, these findings establish a mechanistic link between myotonia and pain and demonstrate that skeletal muscle hyperexcitability is not merely a driver of motor dysfunction but also a potent initiator of persistent alterations in nociceptive processing.

## Results

### Anthracene-9-carboxylic acid (9-AC) produces transient myotonia and persistent pain-like behavior in mice

To model an acute myotonic episode, we administered a single intraperitoneal injection of the ClC-1 antagonist, anthracene-9-carboxylic acid (9-AC; 30 mg/kg(31)). As illustrated in **Fig. 1A**, we used the righting reflex assay to quantify muscle hyperexcitability following drug administration, a validated measure of myotonia based on delayed recovery from forced supine positioning (31, 32). Consistent with previous reports in both mice (39) and rats (31, 34), 9-AC rapidly induced symptoms of muscle stiffness, impaired gait, and delayed movement initiation. Importantly, no sedation, respiratory suppression, or altered consciousness were observed. Quantification of the time to righting reflex (TRR) revealed a significant delay in 9-AC-treated animals compared to vehicle-injected controls, confirming successful induction of transient myotonia (**Fig. 1B-C**).

**Figure 1.**
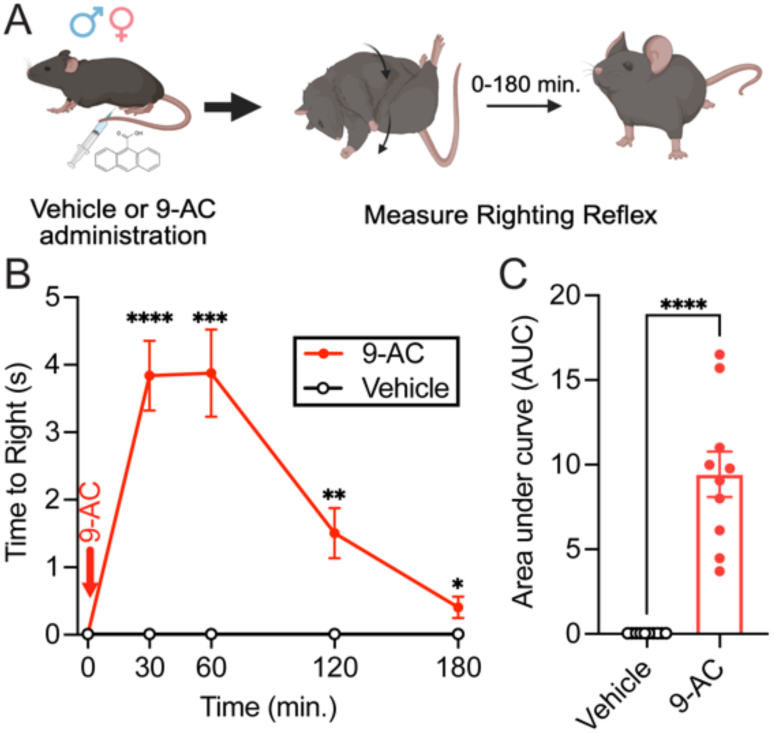
Anthracene-9-carboxylic acid (9-AC) induces transient myotonic behavior in mice. **(A)** Schematic illustrating the experimental timeline. Mice received a single intraperitoneal injection of 9-AC (30 mg/kg), followed by measurement of the righting reflex to assess myotonia-like muscle stiffness *in vivo*. **(B)** Time course of righting reflex latency analyzed by two-way repeated-measures ANOVA followed by Holm-Šidák’s post hoc test. **(C)** Quantification of area under the curve (AUC) analyzed by Student’s t test with Welch’s correction. Data are presented as mean ± SEM (n = 10 mice/group). Individual data points represent values from individual animals. *p < 0.05, **p < 0.01, ***p < 0.001, ****p < 0.0001. In some instances, error bars are too small to be visually discernible.

To determine whether this transient myotonia was sufficient to produce pain-like behavior, we performed longitudinal nociceptive testing over a 72-hour period following injection **(Fig. 2A)**. Mice treated with 9-AC developed robust hypersensitivity across multiple sensory modalities. Compared to vehicle controls, 9-AC-treated mice exhibited reduced mechanical withdrawal thresholds **(Fig. 2B-C)**, increased responsiveness to light brushing **(Fig. 2D-E)**, and exaggerated flinching in response to acetone application **(Fig. 2F-G)**, indicating the development of static mechanical allodynia, dynamic mechanical allodynia, and cold allodynia, respectively. In addition, 9-AC induced thermal hyperalgesia, as measured by reduced withdrawal latency on the 52 °C hot plate test **(Fig. 2H-I)**. These hypersensitivity phenotypes were evident as early as 6 hours post-injection and persisted for up to 48 hours, even though motor symptoms fully resolved by approximately 3 hours post-injection (**Fig. 1B**). Together, these findings demonstrate that a single, transient episode of myotonia is sufficient to initiate a prolonged pain-like behavioral state that persists beyond resolution of overt myotonia.

**Figure 2.**
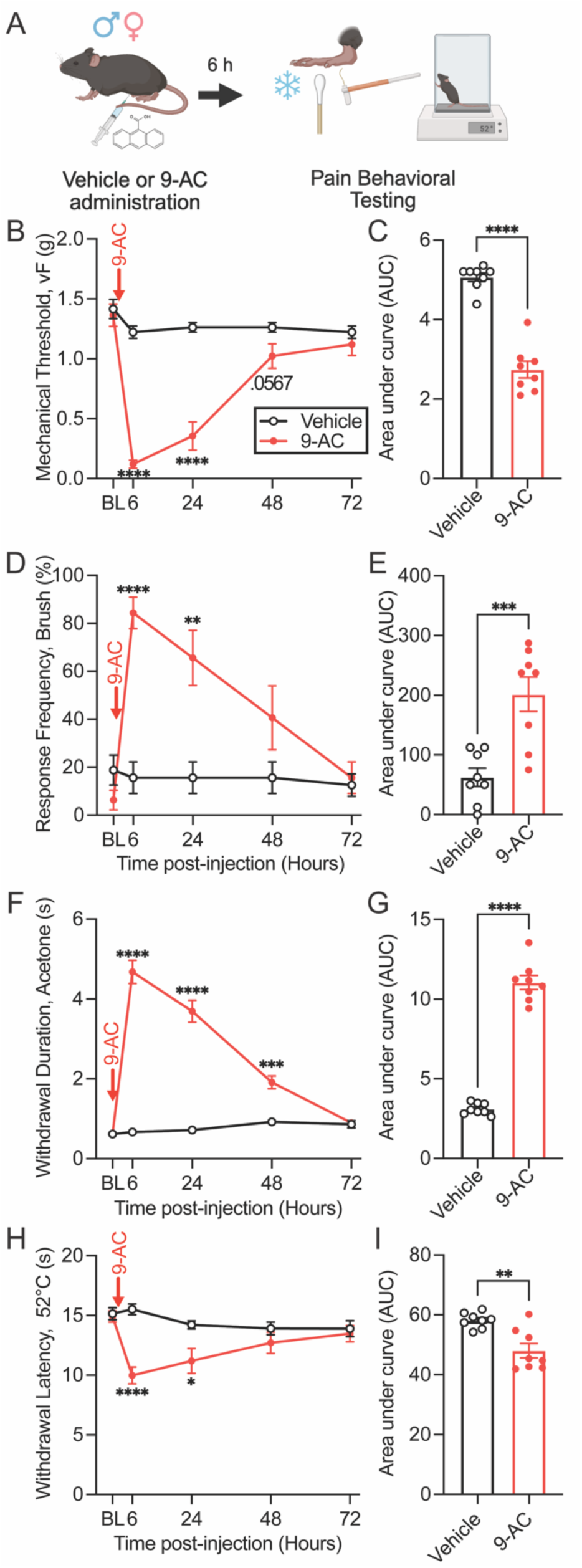
Pharmacological induction of myotonia with 9-AC induces persistent pain-like behavior in mice. **(A)** Schematic of the experimental timeline showing intraperitoneal administration of 9-AC (30 mg/kg) followed by behavioral pain testing over a 72-hour period. **(B-C)** Static mechanical allodynia assessed using von Frey filaments. **(D-E)** Dynamic mechanical allodynia assessed using brush stimulation. **(F-G)** Cold allodynia assessed via acetone droplet application. **(H-I)** Thermal hyperalgesia measured using a 52 °C hot plate assay. **(B, D, F, H)** Time course data were analyzed using two-way ANOVA followed by Dunnett’s post hoc test. **(C, E, G, I)** Area under the curve (AUC) values were analyzed using unpaired two-tailed Student’s t test. Data are presented as mean ± SEM (n = 8 mice/group). Individual data points represent values from individual animals. *p < 0.05, **p < 0.01, ***p < 0.001, ****p < 0.0001. In some instances, error bars are too small to be visually discernible.

### 9-AC increases excitability of small-diameter dorsal root ganglion (DRG) neurons

We next examined the neural mechanisms underlying 9-AC-induced pain-like behavior. Given that pain is driven by peripheral and/or central sensitization of nociceptive pathways (72, 73), we hypothesized that transient myotonia promotes peripheral sensitization through increased excitability of primary afferent or sensory neurons, whose cell bodies are housed in DRGs. To assess this, we performed whole-cell patch-clamp recordings from neurons dissociated from lumbar DRGs, 24 hours after 9-AC injection **(Fig. 3A)**. DRGs were harvested 6 hours after injection and maintained in culture until the time of recording. Representative current-clamp traces revealed increased action potential (AP) firing in small-diameter neurons from 9-AC-treated mice compared to vehicle controls **(Fig. 3B)**. Quantification showed no difference in resting membrane potential **(Fig. 3C)** or rheobase **(Fig. 3D)**, but 9-AC increased the number of evoked APs **(Fig. 3E)**, indicating enhanced excitability.

**Figure 3.**
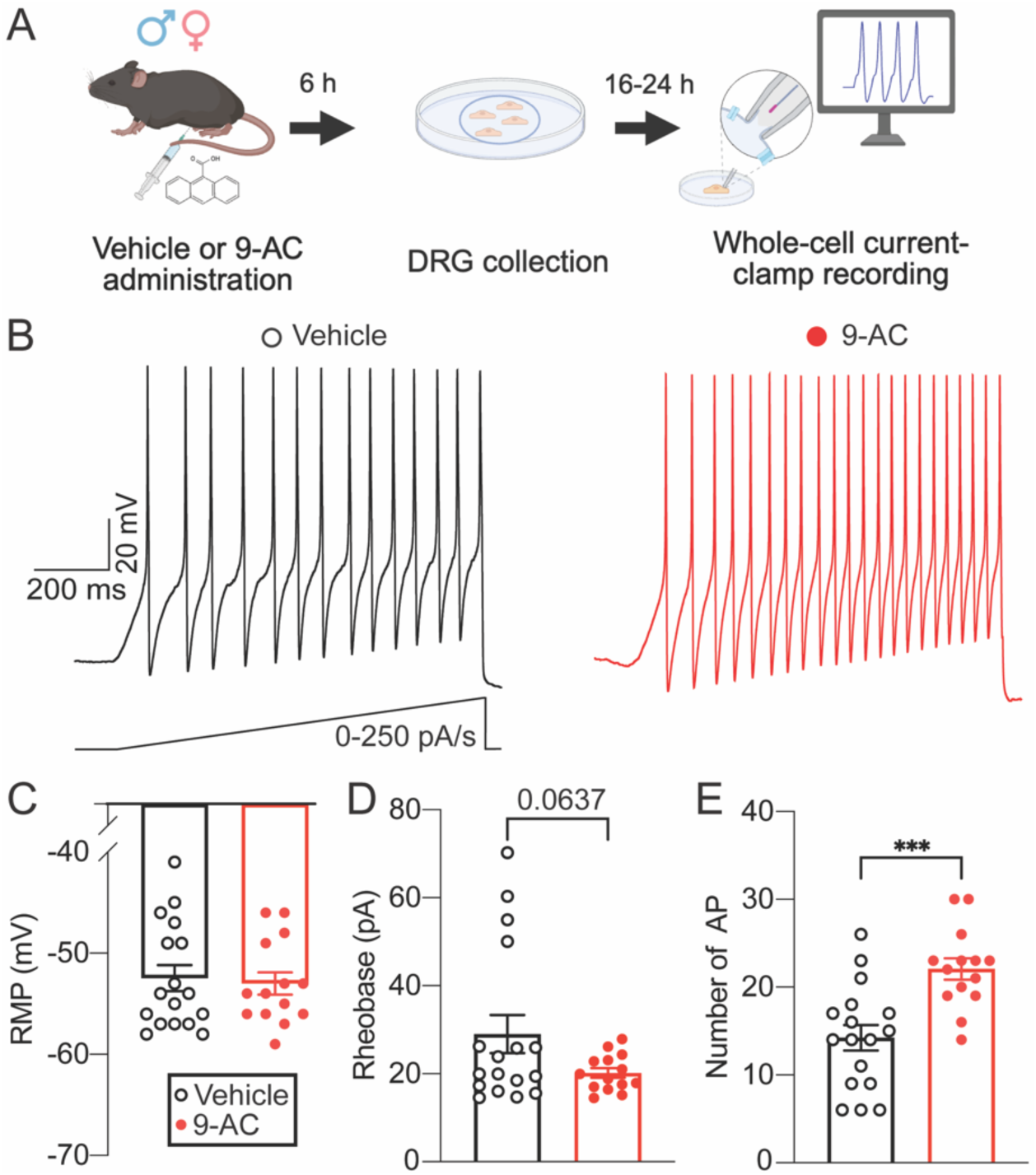
9-AC increases the excitability of dorsal root ganglion neurons. **(A)** Schematic illustrating the experimental timeline. Mice received a single intraperitoneal injection of 9-AC (30 mg/kg), followed by lumbar DRG dissociation 6 hours later. Whole-cell current-clamp recordings were performed in small- and medium-diameter DRG neurons 24 hours post-injection. **(B)** Representative traces showing action potential (AP) firing in sensory neurons from vehicle-treated (left) and 9-AC-treated (right) animals in response to suprathreshold current injection. **(C-E)** Quantification of (**C**) resting membrane potential (RMP), (**D**) rheobase, and (**E**) number of evoked APs. Data were analyzed using the Student’s t test with Welch’s correction. Data are presented as mean ± SEM (n = 15-17 neurons). Individual data points represent values from single neurons. ***p < 0.001.

To determine whether this increased sensory neuron excitability persisted beyond 24 hours, we performed a second set of recordings 48 hours after 9-AC injection (**Supplemental Fig. 1A**). At this later time point, representative traces (**Supplemental Fig. 1B**) and quantification of resting membrane potential (**Supplemental Fig. 1C**), rheobase (**Supplemental Fig. 1D**), and evoked APs (**Supplemental Fig. 1E**) showed no significant differences between groups. These electrophysiological findings align with behavioral results showing resolution of mechanical and thermal hypersensitivity by 48 hours post-injection, with the exception of persistent cold sensitivity (**Fig. 2**). Together, these results suggest that 9-AC induces a transient but reversible increase in the excitability of nociceptive DRG neurons.

Although small- and medium-diameter DRG neurons are the primary mediators of nociception, large-diameter afferents can also contribute to pain signaling under pathological conditions (74–77). We therefore assessed whether 9-AC affects primary afferents with a large cell body diameter. As shown in **Supplemental Fig. 2A**, DRGs were dissociated 6 hours post-injection, and large-diameter neurons were recorded 24 hours later. Representative current-clamp traces showed similar rheobase thresholds between groups (**Supplemental Fig. 2B**), and quantification revealed no differences in resting membrane potential or rheobase (**Supplemental Fig. 2C-D**), suggesting that large-diameter afferents are not sensitized in this model.

### 9-AC reduces A-fiber component of the compound action potential

To further evaluate changes in peripheral input, we examined compound action potentials (CAPs) from sciatic nerves *ex vivo*, 24 hours after 9-AC injection. CAPs provide a measure of axonal recruitment and excitability in response to electrical stimulation **(Fig. 4A)**. Analysis of C-fiber responses revealed no change in conduction velocity or area under the curve (AUC, **Supplemental Fig. 3)**, indicating intact unmyelinated fiber function. In contrast, A-fiber responses were significantly impaired following 9-AC treatment. Representative traces showed reduced A-wave amplitudes across a range of stimulation intensities (**Fig. 4B**) and while conduction velocity (**Fig. 4C**), and activity-dependent suppression of A-fibers at 30 Hz (2.1 ± 0.7 vs 0.6 ± 2.1%) and 100 Hz (14.3 ± 6.9 vs 3.2 ± 3.9%) (vehicle vs 9-AC treated, respectively) remained unchanged, several recruitment metrics were altered. Specifically, 9-AC shifted the A-wave recruitment curve to the right (**Fig. 4D**), reduced the AUC Max **(Fig. 4E**), increased the current required to reach half-maximal activation (**Fig. 4F**), and produced a trend toward a shallower slope in the stimulus-response function (**Fig. 4G**). These results indicate that 9-AC reduces the number of functional A-fibers and/or their excitability without producing overt conduction block.

**Figure 4.**
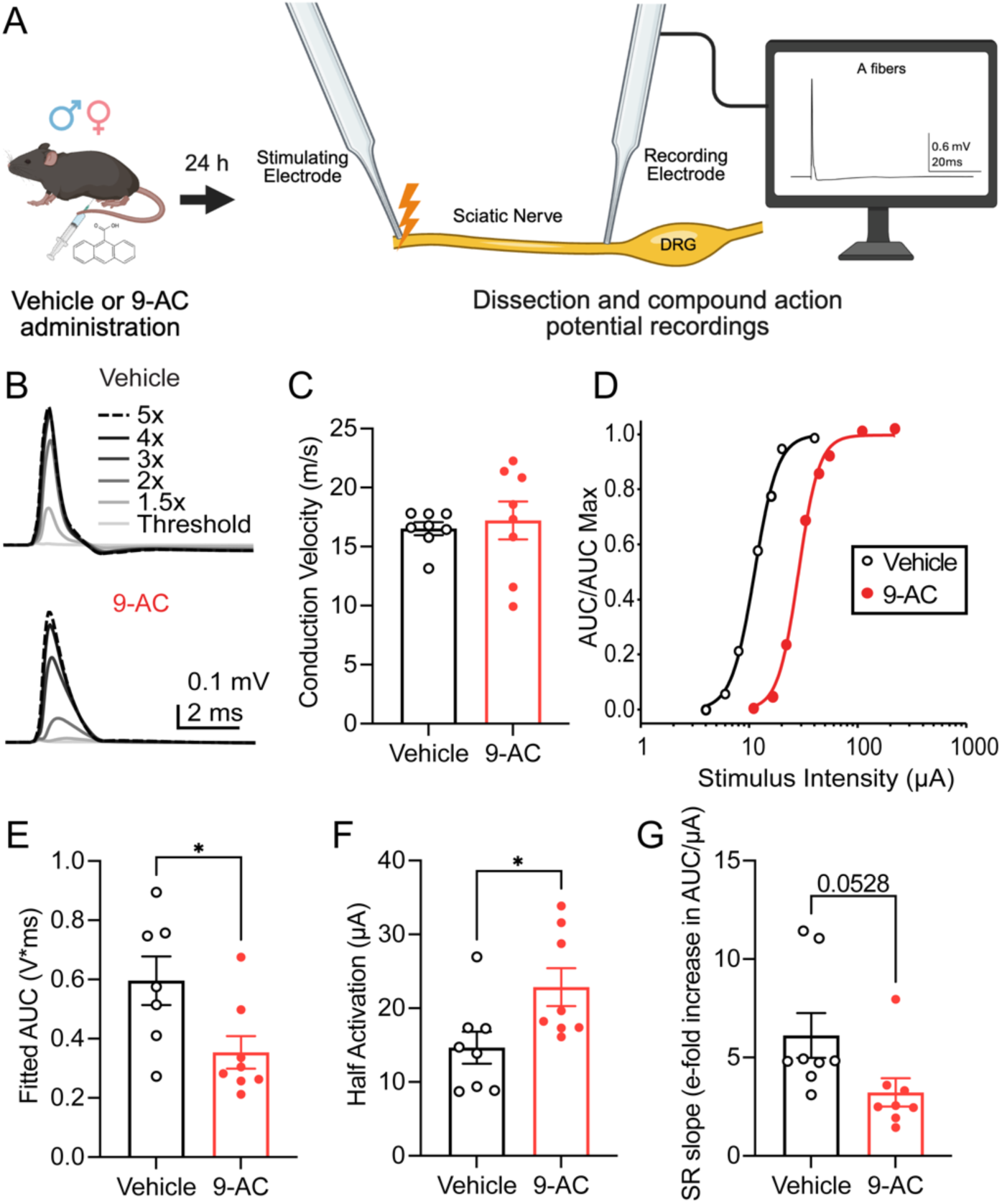
9-AC reduces A-fiber compound action potential responses. **(A)** Schematic illustrating the experimental timeline. Mice received a single intraperitoneal injection of 9-AC (30 mg/kg), followed by sciatic nerve dissection 24 hours later. Compound action potential recordings were performed *ex vivo* to assess A-fiber function. **(B)** Representative traces showing responses to threshold and 1.5-5x threshold stimulation intensities in sciatic nerves from vehicle-treated (top) and 9-AC-treated (bottom) animals. **(C-G)** Quantification of (**C**) conduction velocity, (**D**) AUC/AUC max, (**E**) fitted AUC, (**F**) half activation, and (**G**) SR slope. Data in panels **C** and **E-G** were analyzed using the Student’s t test with Welch’s correction. Data are presented as mean ± SEM (n = 7-8 animals/group). Individual data points represent values from single animals (average of 2 nerves/animal). *p < 0.05.

### 9-AC increases the excitability of superficial dorsal horn neurons

Peripheral sensory neurons are the first relay in the nociceptive circuit, but central circuits ultimately determine pain perception. The spinal dorsal horn serves as the first central relay for peripheral nociceptive information and is therefore well positioned to contribute to pain hypersensitivity following transient myotonia. To determine whether 9-AC alters the intrinsic excitability of dorsal horn neurons, we performed whole-cell patch-clamp recordings from neurons located within laminae I-II of the lumbar spinal cord 24 hours after drug administration **(Supplemental Fig. 4A)**. Superficial dorsal horn neurons displayed several distinct firing patterns, including single, phasic, tonic, and delay firing phenotypes **(Supplemental Fig. 4B)**. Analysis of firing frequency revealed a robust effect of current injection but no effect of treatment or current-by-treatment interaction **(Supplemental Fig. 4C)**. Likewise, the distribution of firing phenotypes did not differ between vehicle- and 9-AC-treated mice **(Supplemental Fig. 4D)**. Resting membrane potential, action potential threshold, threshold distance (Threshold distance = *V*_threshold_−*V*_rest_), rheobase, and spike latency were also similar between groups **(Supplemental Fig. 4E-I)**, suggesting that transient myotonia does not broadly alter the intrinsic properties or cellular composition of the superficial dorsal horn neuronal population.

Given the marked heterogeneity of dorsal horn neurons, we next stratified analyses by firing phenotype **(Supplemental Fig. 5)**. In delay firing neurons, 9-AC increased evoked action potential firing **(Supplemental Fig. 5B)**, hyperpolarized the action potential threshold **(Supplemental Fig. 5D)**, reduced threshold distance **(Supplemental Fig. 5E)**, and shortened spike latency **(Supplemental Fig. 5G)**. Tonic and phasic firing neurons, by contrast, showed no significant changes in firing frequency or intrinsic membrane properties **(Supplemental Fig. 5H-U)**. Thus, transient myotonia selectively enhances the excitability of delay firing neurons while leaving the broader electrophysiological profile of the superficial dorsal horn largely intact.

### 9-AC enhances sensory-evoked activity in the parabrachial nucleus

To further identify central contributors to 9-AC-induced hypersensitivity, we next investigated the parabrachial nucleus (PBN), a key brainstem structure that integrates spinal nociceptive input and projects to higher-order brain regions involved in affective and motivational aspects of pain(78–80). We used *in vivo* fiber photometry to monitor calcium activity in glutamatergic PBN neurons in awake, freely moving mice. Mice received stereotaxic injections of an adeno-associated virus encoding GCaMP6s, a genetically encoded calcium sensor that fluoresces in response to an elevation in intracellular calcium, and serves as a proxy for neuronal activation. The virus was targeted to the PBN to drive selective expression in local glutamatergic neurons. Additionally, an optical fiber cannula was implanted above the injection site to allow repeated recordings from awake, freely behaving animals. After baseline recordings, we re-assessed sensory-evoked PBN activity 24 hours after 9-AC or vehicle administration. After a one-month washout period, animals received the alternate treatment to enable within-subject comparisons (**Fig. 5A**).

**Figure 5.**
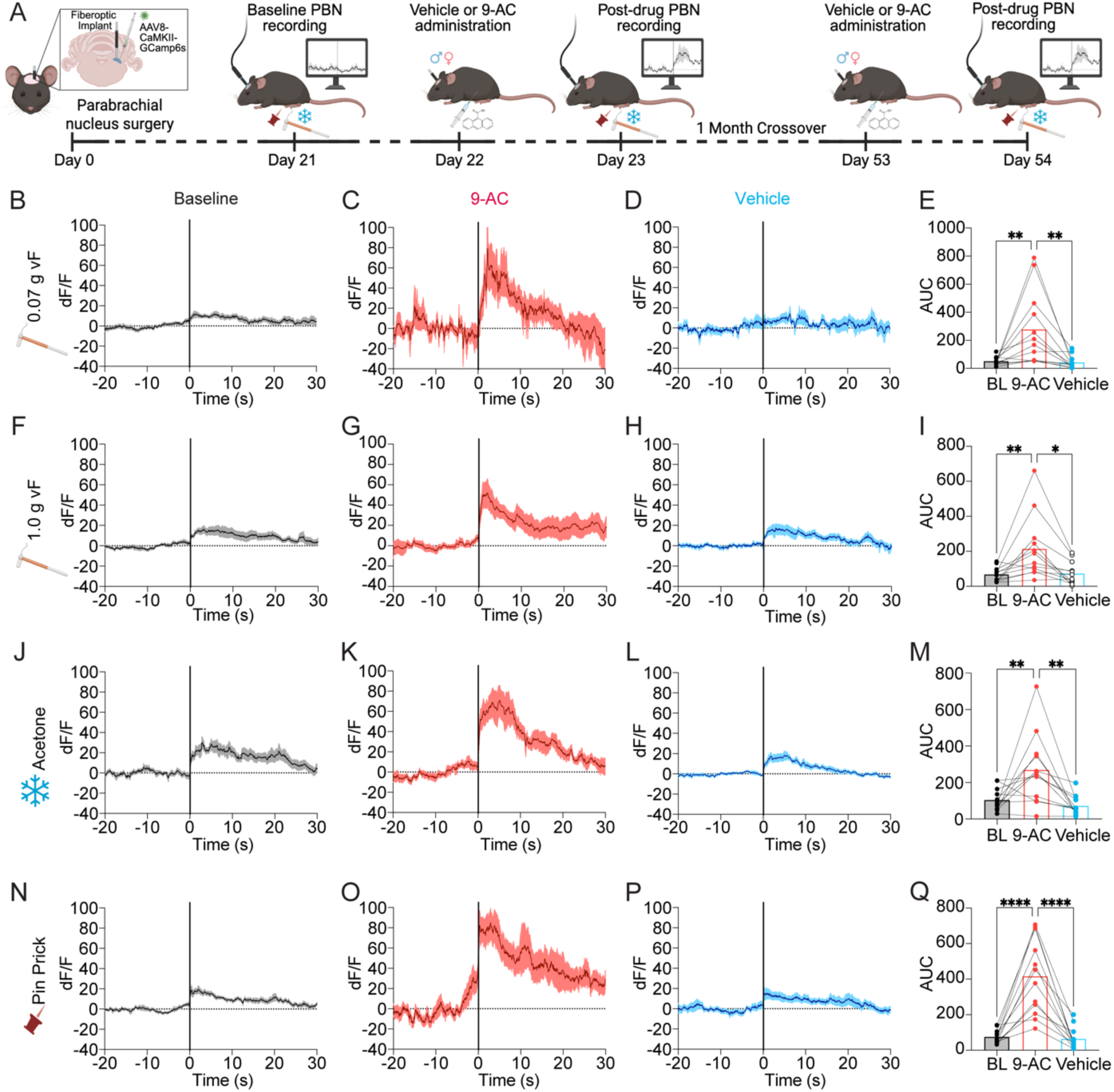
9-AC increases parabrachial nucleus activity in response to somatosensory stimuli. **(A)** Schematic illustrating the experimental timeline. Mice received intra-parabrachial injections of AAV-GCaMP6s and were implanted with a fiberoptic cannula. Three weeks later, baseline *in vivo* fiber photometry recordings were performed. Mice then received a single intraperitoneal injection of 9-AC (30 mg/kg) or vehicle, followed 24 hours later by a second fiber photometry recording. One month later, a crossover was performed and the experiment was repeated. **(B-D)** Representative traces of normalized calcium-dependent fluorescence in response to a 0.07 g von Frey (vF) filament, and **(E)** quantification of the area under the curve (AUC). **(F-H)** Representative traces in response to a 1.0 g vF filament, and **(I)** corresponding AUC analysis. **(J-L)** Representative traces in response to acetone application, and **(M)** corresponding AUC analysis. **(N-P)** Representative traces in response to blunted pinprick stimulation, and **(Q)** corresponding AUC analysis. Data were analyzed using one-way ANOVA followed by Dunnett’s post hoc test. Data are presented as mean ± SEM (n = 12 mice/group). Individual data points represent values from individual animals. *p < 0.05, **p < 0.01, ****p < 0.0001.

We observed robust enhancement of sensory-evoked activity in the PBN following 9-AC treatment. Light mechanical stimulation with a 0.07 g von Frey filament elicited exaggerated calcium transients (**Fig. 5B-D**), with a significant increase in AUC compared to vehicle (**Fig. 5E**). Similar increases were observed with a 1.0 g filament (**Fig. 5F-H**), acetone application (**Fig. 5J-L**), and pinprick stimulation (**Fig. 5N-P**), each showing corresponding AUC increases (**Fig. 5I, 5M, 5Q**). These findings indicate that a single episode of myotonia is sufficient to sensitize central pain-processing circuits, supporting a role for central amplification in sustaining nociceptive hypersensitivity.

### 9-AC alters tetrodotoxin-sensitive and tetrodotoxin-resistant sodium current fractions in sensory neurons

Given the increased excitability of small-diameter DRG neurons following transient myotonia, we next sought to identify the ionic mechanisms underlying this phenotype. Whole-cell voltage-clamp recordings revealed larger inward sodium currents in sensory neurons isolated from 9-AC-treated mice compared to vehicle controls **(Fig. 6B)**. Quantification of the current density-voltage relationship demonstrated increased sodium current density following 9-AC treatment **(Fig. 6C)**, which was associated with a significant increase in peak sodium current density **(Fig. 6D)**. In contrast, voltage-dependent activation and steady-state inactivation properties were preserved, as activation and inactivation curves overlapped between groups and no differences in half-maximal activation or inactivation voltages were detected **(Fig. 6E-H)**. These findings indicate that transient myotonia increases the magnitude of sodium currents in sensory neurons without appreciably altering channel gating properties.

**Figure 6.**
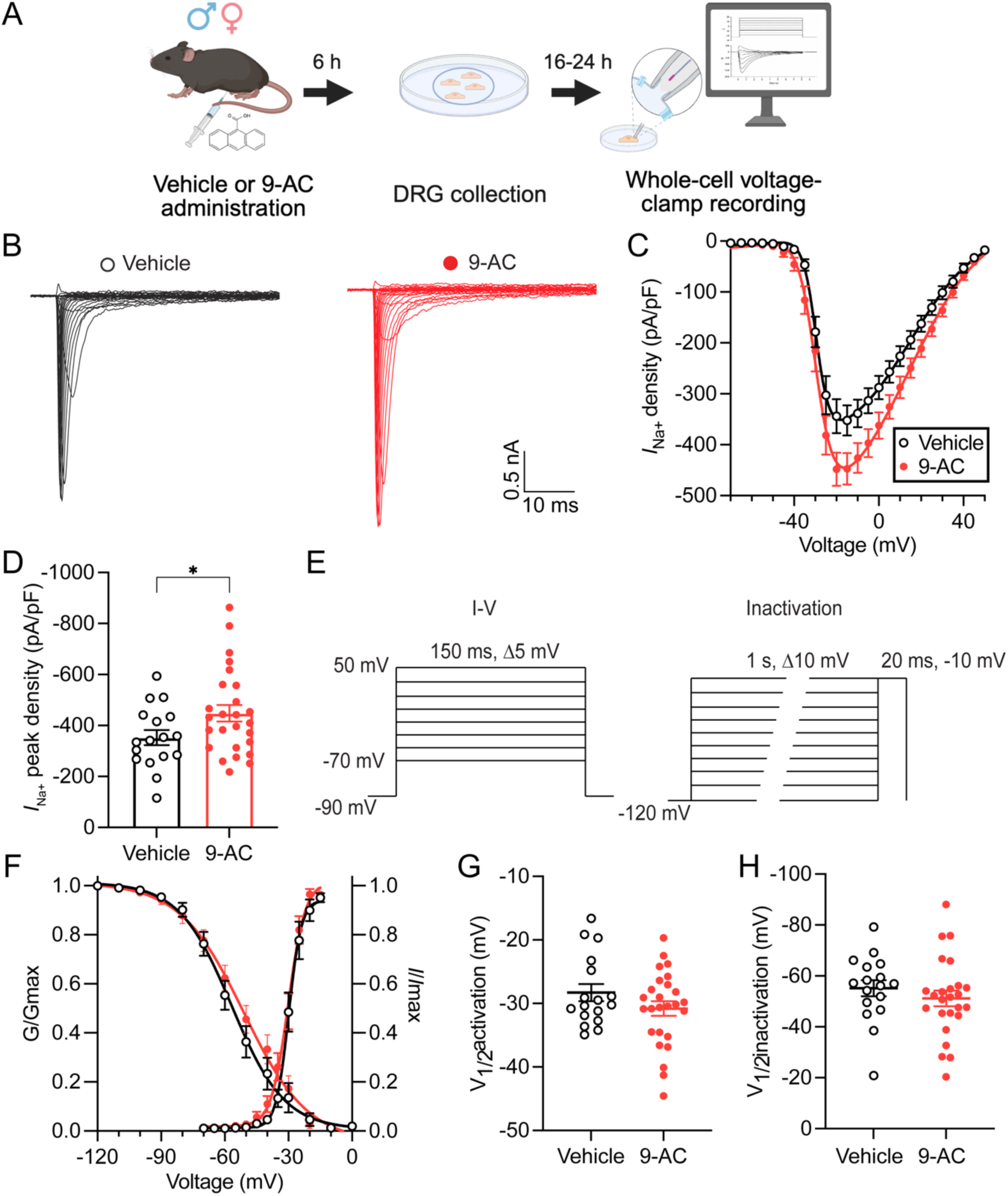
9-AC increases voltage-gated sodium current density in dorsal root ganglion neurons without altering channel gating properties. **(A)** Schematic illustrating the experimental timeline. Mice received a single intraperitoneal injection of 9-AC (30 mg/kg), followed by lumbar DRG dissociation 6 hours later. Whole-cell voltage-clamp recordings were performed in small- and medium-diameter DRG neurons 16–24 hours after dissociation. **(B)** Representative voltage-gated sodium current (*I*_Na_) traces recorded from sensory neurons isolated from vehicle-treated (left) and 9-AC-treated (right) animals. **(C)** Current density-voltage (I-V) relationships demonstrating increased sodium current density following 9-AC treatment. **(D)** Quantification of peak *I*_Na_ density. **(E)** Voltage protocols used to assess sodium channel activation and steady-state inactivation. **(F)** Activation (G/Gmax) and steady-state inactivation (*I*/*I*max) curves for sodium currents in vehicle- and 9-AC-treated neurons. (**G-H**) Quantification of the half-maximal voltage (V_1/2_) of **(G)** activation and **(H)** steady-state inactivation. Data were analyzed using Student’s t test with Welch’s correction. Data are presented as mean ± SEM (**C** and **F**) or as individual data points with mean ± SEM (**D**, **G,** and **H**) (n = 17-26 neurons). Individual data points represent values from single neurons. *p < 0.05.

To determine whether specific sodium channel subtypes contributed to this increase in sodium current, we next separated tetrodotoxin-sensitive (TTX-S) and tetrodotoxin-resistant (TTX-R) current fractions by digital subtraction analysis as previously described (**Fig. 7A**) (81). Although neither current fraction reached statistical significance individually, 9-AC treatment produced a reciprocal redistribution of sodium current composition, characterized by a trend toward reduced TTX-S current fraction and a corresponding trend toward increased TTX-R current fraction **(Fig. 7B-C)**. Given that TTX-R currents in small-diameter nociceptors are mediated predominantly by Na_V_1.8 and Na_V_1.9 channels (82), this reciprocal shift in sodium current composition is consistent with greater functional contributions of TTX-R conductances to the enhanced excitability of sensory neurons following transient myotonia.

**Figure 7.**
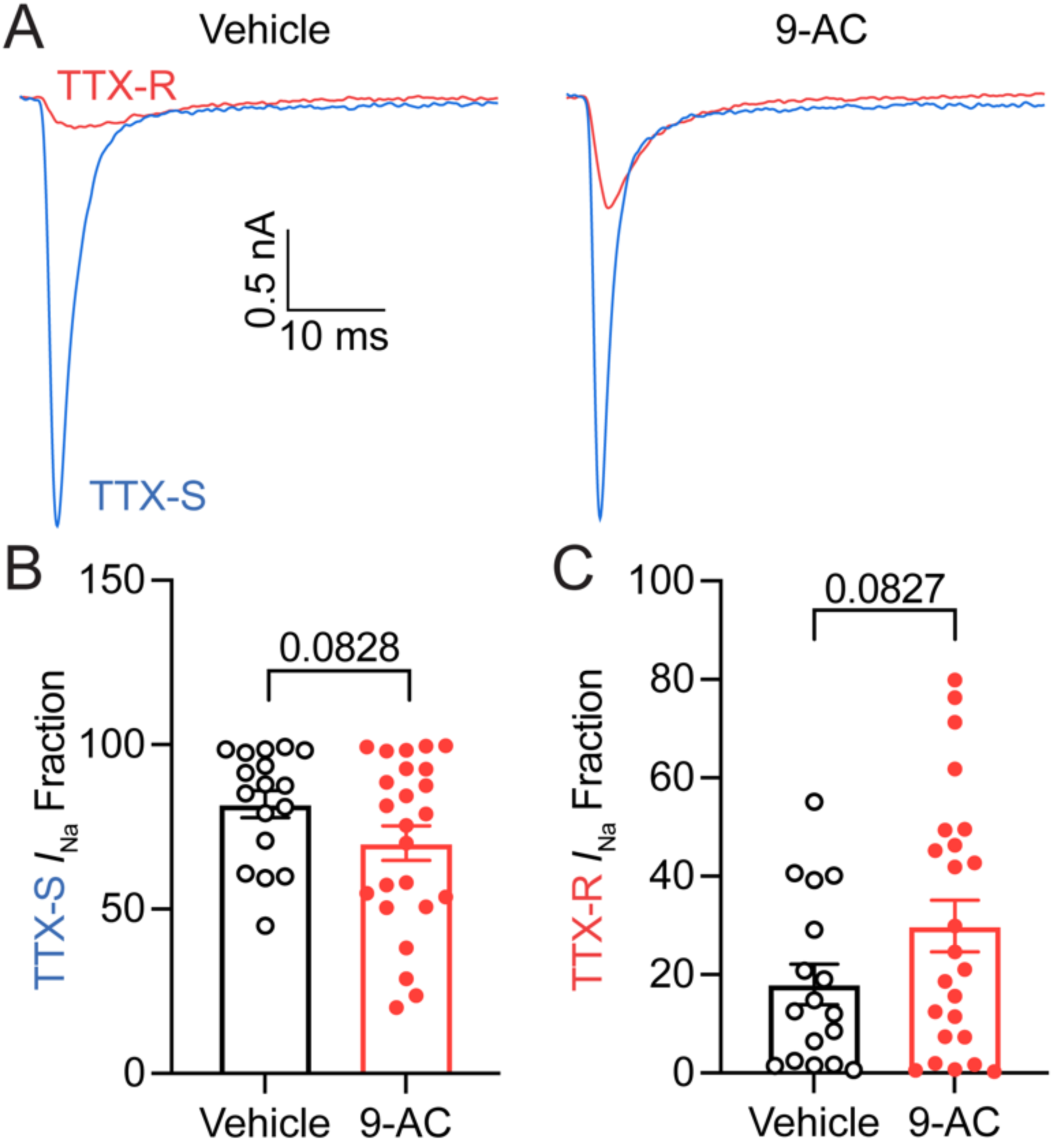
TTX-sensitive and TTX-resistant sodium current fractions following 9-AC administration. **(A)** Representative digitally subtracted tetrodotoxin-sensitive (TTX-S; blue) and tetrodotoxin-resistant (TTX-R; red) sodium currents from sensory neurons isolated from vehicle-treated (left) and 9-AC-treated (right) animals. TTX-S and TTX-R currents were determined offline by digital subtraction as previously described (81). **(B-C)** Quantification of **(B)** TTX-S and **(C)** TTX-R sodium current fractions in vehicle- and 9-AC-treated neurons. Data were analyzed using Student’s t test with Welch’s correction. Data are presented as mean ± SEM (n = 16-24 neurons). Individual data points represent values from single neurons. Trending but non-significant p values are indicated on the graph.

### Suzetrigine reduces sensory neuron excitability and alleviates 9-AC-induced pain-like behavior

To directly test the contribution of Na_V_1.8 to sensory neuron hyperexcitability, we acutely applied the selective Na_V_1.8 inhibitor Suzetrigine (5 μM) during whole-cell current-clamp recordings **(Fig. 8A)**. Representative traces demonstrated that Suzetrigine markedly reduced action potential firing in neurons isolated from both vehicle- and 9-AC-treated animals **(Fig. 8B)**. Quantification revealed no effects on resting membrane potential or rheobase **(Fig. 8C-D)**, indicating that Na_V_1.8 inhibition did not broadly alter intrinsic membrane properties. In contrast, Suzetrigine significantly reduced the number of evoked action potentials, abolishing the hyperexcitability observed following 9-AC treatment **(Fig. 8E)**. Analysis of action potential waveform properties similarly revealed minimal effects on amplitude, half-width, threshold, rise time, decay time, medium afterhyperpolarization, spike timing, or cell size **(Fig. 8F-N)**. Together, these findings demonstrate that acute Suzetrigine application reduces repetitive firing in sensory neurons from both control and 9-AC-treated mice.

**Figure 8.**
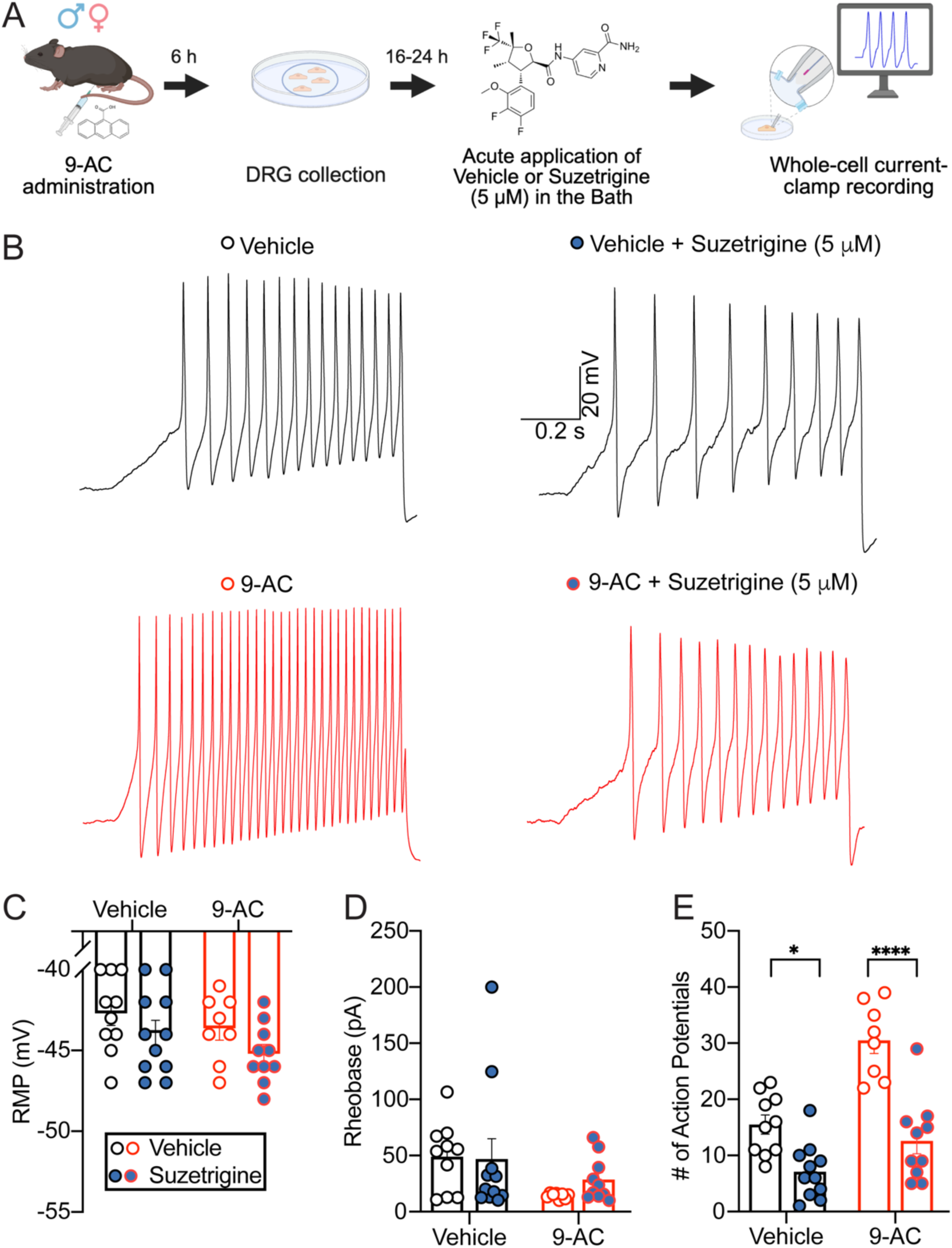
Acute pharmacological inhibition of Na_V_1.8 reduces sensory neuron hyperexcitability following 9-AC administration. **(A)** Schematic illustrating the experimental timeline. Mice received a single intraperitoneal injection of 9-AC (30 mg/kg), followed by lumbar DRG dissociation 6 hours later. Whole-cell current-clamp recordings were performed in small- and medium-diameter DRG neurons 16–24 hours after dissociation. During recordings, neurons were acutely exposed to either vehicle or the selective Na_V_1.8 inhibitor Suzetrigine (5 μM). **(B)** Representative traces showing action potential (AP) firing in sensory neurons from vehicle-treated (top) and 9-AC-treated (bottom) animals in the absence (left) or presence (right) of Suzetrigine. **(C-E)** Quantification of **(C)** resting membrane potential (RMP), **(D)** rheobase, and **(E)** number of evoked APs. Acute application of Suzetrigine significantly reduced AP firing in sensory neurons from both vehicle- and 9-AC-treated animals without altering RMP or rheobase. Data were analyzed using Student’s t test with Welch’s correction. Data are presented as mean ± SEM (n = 8-11 mice/group). Individual data points represent values from single neurons. *p < 0.05, ****p < 0.0001.

We next asked whether pharmacological inhibition of Na_V_1.8 could alleviate pain-like behavior *in vivo*. Mice received vehicle or Suzetrigine (10 mg/kg, i.p. (83)) 24 hours after 9-AC administration, a time point at which robust hypersensitivity is present despite complete resolution of motor symptoms **(Fig. 9A)**. Acute administration of Suzetrigine significantly attenuated static mechanical allodynia **(Fig. 9B-C)**, dynamic mechanical allodynia **(Fig. 9D-E)**, cold allodynia **(Fig. 9F-G)**, and thermal hyperalgesia **(Fig. 9H-I)**. Importantly, Suzetrigine produced little effect in vehicle-treated animals **(Fig. 9B-I)**, suggesting that Na_V_1.8 inhibition preferentially normalizes pathological sensory processing rather than suppressing physiological nociception. Collectively, these findings demonstrate robust efficacy of Na_V_1.8-directed pharmacological inhibition against transient myotonia-induced pain-like behavior.

**Figure 9.**
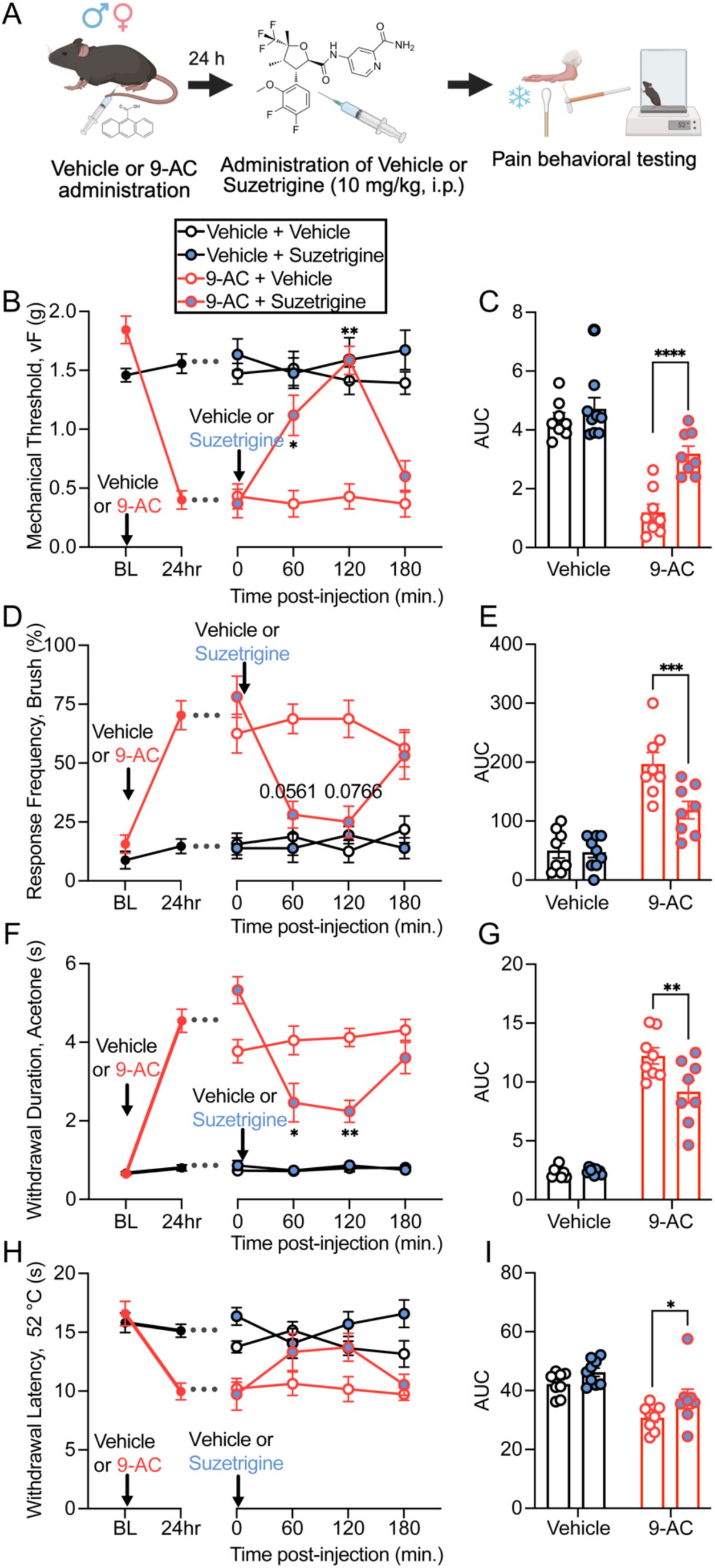
Acute pharmacological inhibition of Na_V_1.8 alleviates 9-AC-induced pain-like behaviors. **(A)** Schematic of the experimental timeline showing intraperitoneal administration of 9-AC (30 mg/kg) followed 24 hours later by intraperitoneal administration of vehicle or the selective Na_V_1.8 inhibitor Suzetrigine (10 mg/kg). Behavioral pain testing was performed for 3 hours following drug administration. **(B-C)** Static mechanical allodynia assessed using von Frey filaments. **(D-E)** Dynamic mechanical allodynia assessed using brush stimulation. **(F-G)** Cold allodynia assessed via acetone droplet application. (H-I) Thermal hyperalgesia measured using a 52 °C hot plate assay. **(B, D, F, H)** Time course data were analyzed using two-way ANOVA followed by Tukey’s post hoc test. **(C, E, G, I)** Area under the curve (AUC) values were analyzed using two-way ANOVA followed by Tukey’s post hoc test. Data are presented as mean ± SEM (n = 8-9 mice/group). Individual data points represent values from individual animals. *p < 0.05, **p < 0.01, ****p < 0.0001. In some instances, error bars are too small to be visually discernible.

### Mice with myotonia congenita exhibit persistent pain-like behavior responsive to Na_V_1.8-directed analgesia

Although 9-AC produces a transient episode of myotonia, patients with myotonia congenita experience recurrent myotonia throughout life due to inherited loss-of-function mutations in *CLCN1* (84). We therefore hypothesized that mice with genetically induced myotonia would exhibit persistent pain-like behavior and sensory neuron hyperexcitability. To test this hypothesis, we used the well-characterized adr (arrested development of righting response) mouse model of myotonia congenita, which carries a mutation in *Clcn1* resulting in myotonia due to impaired skeletal muscle ClC-1 function (**Fig. 10A** and **Supplemental Fig. 6**) (15, 85, 86). Consistent with the established phenotype of this model, homozygous *Clcn1*^adr/adr^ mice exhibited reduced body weight, impaired grip strength, and prolonged righting times compared with wildtype and heterozygous littermates, whereas heterozygous *Clcn1*^adr/+^ mice were indistinguishable from wildtype controls **(Supplemental Fig. 6)**.

**Figure 10.**
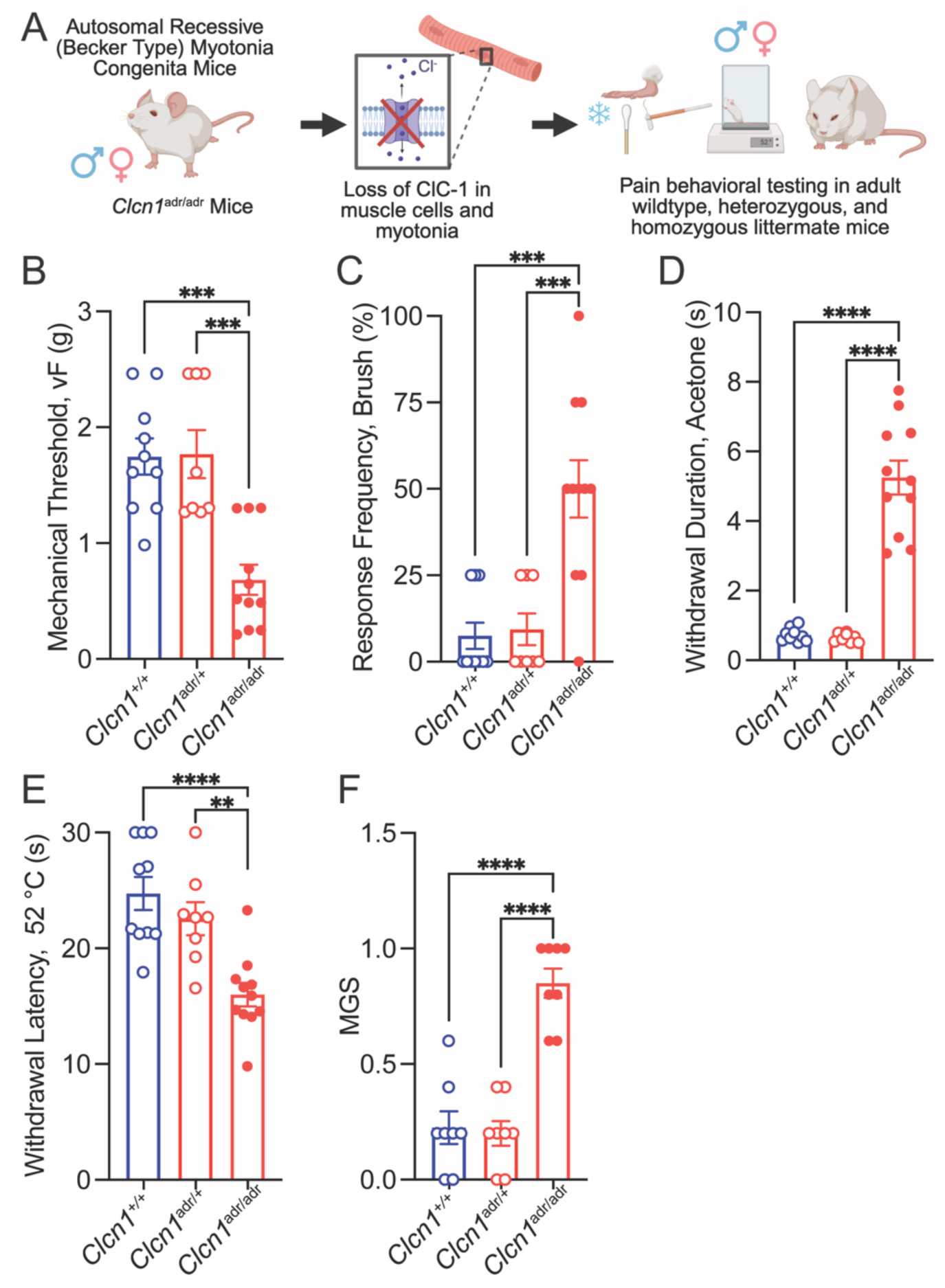
Mice with myotonia congenita exhibit pain-like behaviors. **(A)** Schematic illustrating the experimental design. Adult wildtype (*Clcn1*^+/+^), heterozygous (*Clcn1*^adr/+^), and homozygous (*Clcn1*^adr/adr^) littermate mice were subjected to behavioral testing to determine whether loss of CLC-1 function and myotonia are associated with altered nociceptive processing. **(B)** Static mechanical sensitivity assessed using von Frey filaments. **(C)** Dynamic mechanical sensitivity assessed using brush stimulation. **(D)** Cold sensitivity assessed via acetone droplet application. **(E)** Thermal sensitivity measured using a 52 °C hot plate assay. **(F)** Mouse Grimace Scale (MGS) scores as an index of spontaneous pain. Data were analyzed using one-way ANOVA followed by Tukey’s multiple comparisons test. Data are presented as mean ± SEM (n = 8-11 mice/group). Individual data points represent values from individual animals. **p < 0.01, ***p < 0.001, ****p < 0.0001.

Behavioral testing revealed a genotype-dependent pain-like phenotype. Compared with wildtype and heterozygous littermates, homozygous *Clcn1*^adr/adr^ mice exhibited reduced withdrawal thresholds to von Frey stimulation **(Fig. 10B)**, increased responsiveness to dynamic brushing **(Fig. 10C)**, exaggerated responses to acetone application **(Fig. 10D)**, and reduced withdrawal latency on the 52 °C hot plate assay **(Fig. 10E)**. In addition, *Clcn1*^adr/adr^ mice displayed elevated mouse grimace scores (MGS), indicating the presence of spontaneous pain-like behavior **(Fig. 10F)**. In contrast, heterozygous *Clcn1*^adr/+^ mice did not differ from wildtype controls across these behavioral measures **(Fig. 10B-F)**. Together, these findings demonstrate that chronic genetic myotonia is associated with persistent evoked and spontaneous pain-like behaviors.

To determine whether chronic myotonia alters peripheral sensory neuron physiology and whether these neurons remain responsive to Na_V_1.8 inhibition, we performed whole-cell current-clamp recordings from small- and medium-diameter lumbar DRG neurons isolated from adult wildtype and *Clcn1*^adr/adr^ mice (**Fig. 11A**). Baseline repetitive firing did not differ significantly between genotypes. Acute Suzetrigine application reduced repetitive action potential firing in neurons from both wildtype and *Clcn1*^adr/adr^ mice (**Fig. 11B-E**). Additional analysis revealed genotype-associated differences in selected membrane and action potential properties, indicating that chronic genetic myotonia is accompanied by electrophysiological remodeling distinct from the enhanced repetitive firing observed following transient 9-AC-induced myotonia **(Fig. 11F-N)**.

**Figure 11.**
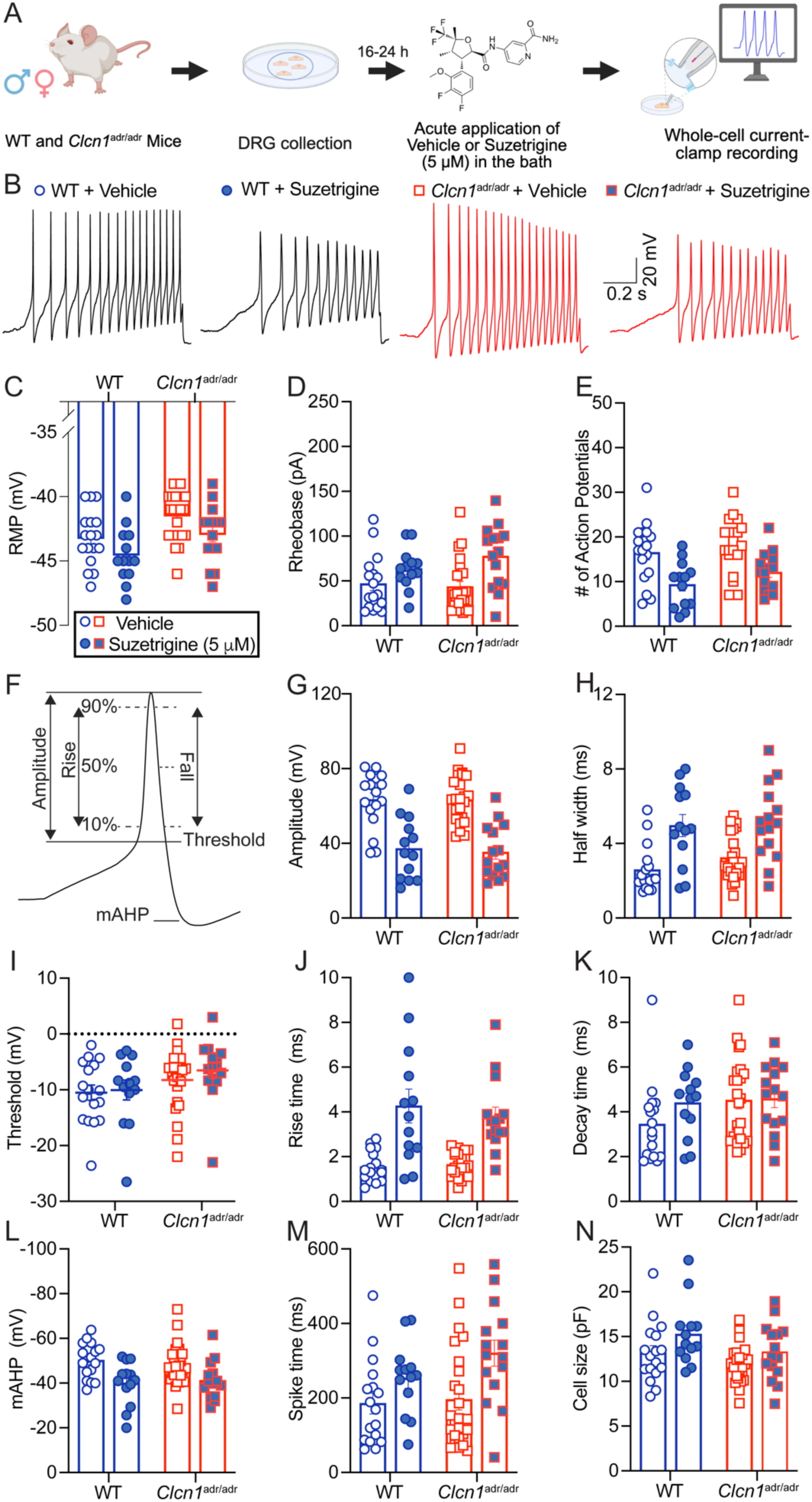
Acute Suzetrigine alters action potential firing in sensory neurons from wildtype and *Clcn1*^adr/adr^ mice. **(A)** Schematic illustrating the experimental timeline. Lumbar dorsal root ganglia (DRG) were collected from adult wildtype (*Clcn1*^+/+^) and homozygous myotonic (*Clcn1*^adr/adr^) mice. Following dissociation, whole-cell current-clamp recordings were performed in small- and medium-diameter DRG neurons 16–24 hours later. During recordings, neurons were acutely exposed to either vehicle or the selective Na_V_1.8 inhibitor Suzetrigine (5 μM). **(B)** Representative traces showing action potential (AP) firing in sensory neurons from wildtype (left) and *Clcn1*^adr/adr^ (right) mice in the absence or presence of Suzetrigine. **(C-E)** Quantification of **(C)** resting membrane potential (RMP), **(D)** rheobase, and **(E)** number of evoked APs. **(F)** Schematic illustrating the electrophysiological parameters measured from individual action potentials. **(G-N)** Quantification of **(G)** AP amplitude, **(H)** half-width, **(I)** threshold, **(J)** rise time, **(K)** decay time, **(L)** medium afterhyperpolarization (mAHP), **(M)** spike time, and **(N)** cell size. Data were analyzed using two-way ANOVA. Data are presented as mean ± SEM (n = 13-24 neurons). Individual data points represent values from single neurons. See Supplemental Statistics Table for full results.

We next asked whether Na_V_1.8 inhibition could alleviate pain-like behavior *in vivo*. Acute administration of Suzetrigine significantly attenuated static mechanical allodynia **(Fig. 12B)**, dynamic mechanical allodynia **(Fig. 12C)**, cold allodynia **(Fig. 12D)**, thermal hyperalgesia **(Fig. 12E)**, and elevated mouse grimace scores **(Fig. 12F)** in *Clcn1*^adr/adr^ mice. In contrast, Suzetrigine produced minimal effects in wildtype animals **(Fig. 12 B-F)**. Collectively, these findings demonstrate that chronic genetic myotonia produces persistent pain-like behavior that is strongly responsive to Na_V_1.8-directed analgesia, despite an electrophysiological phenotype distinct from that observed following transient myotonia.

**Figure 12.**
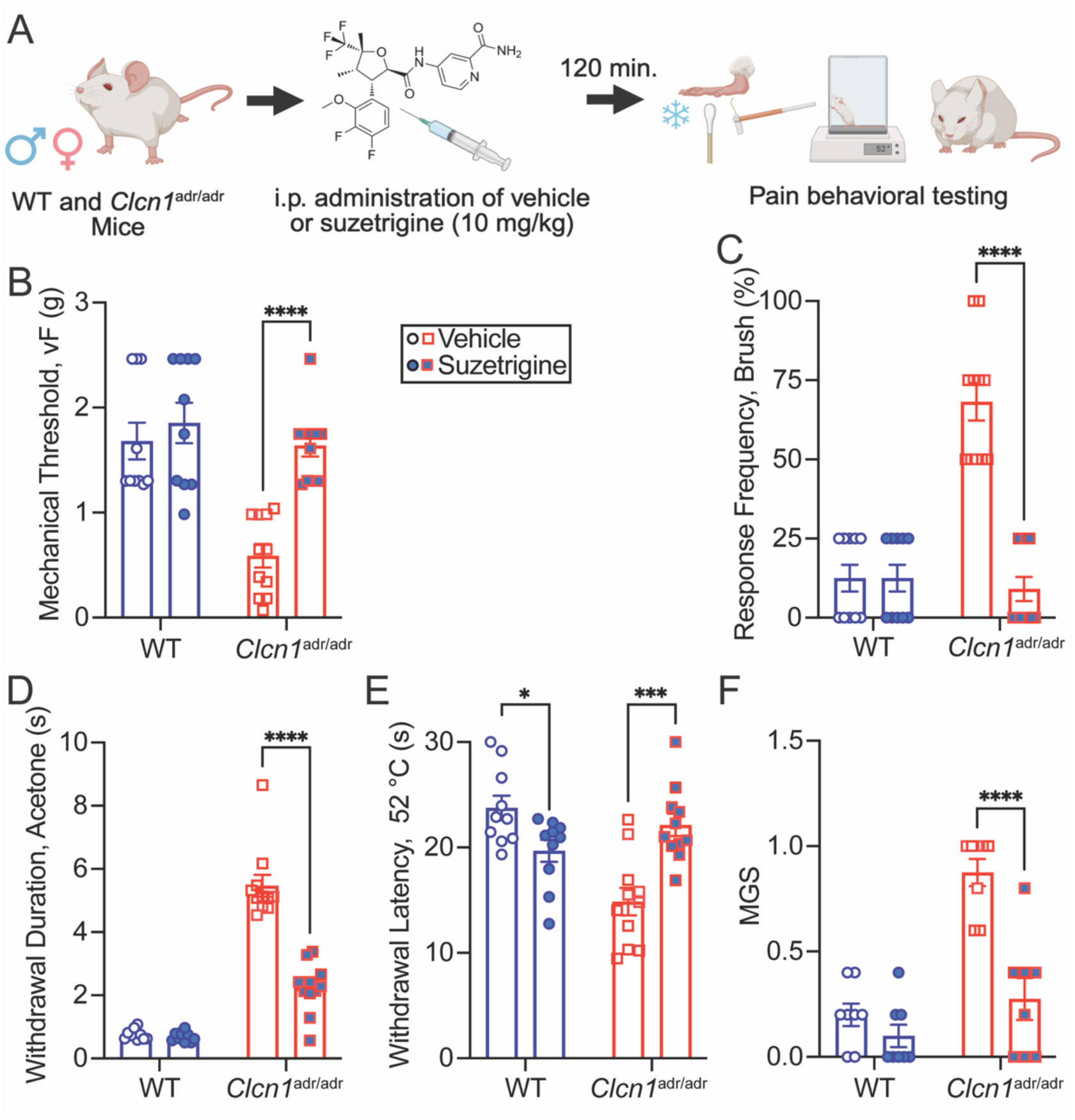
Pharmacological inhibition of Na_V_1.8 alleviates pain-like behaviors in mice with myotonia congenita. **(A)** Schematic of the experimental timeline showing intraperitoneal administration of vehicle or the selective Na_V_1.8 inhibitor Suzetrigine (10 mg/kg) followed 120 minutes later by behavioral testing in adult wildtype (*Clcn1*^+/+^) and homozygous myotonic (*Clcn1*^adr/adr^) mice. **(B)** Static mechanical sensitivity assessed using von Frey filaments. **(C)** Dynamic mechanical sensitivity assessed using brush stimulation. **(D)** Cold sensitivity assessed via acetone droplet application. **(E)** Thermal sensitivity measured using a 52 °C hot plate assay. **(F)** Mouse Grimace Scale (MGS) scores as an index of spontaneous pain. Data were analyzed using two-way ANOVA followed by Holm-Šidák’s post hoc test. Data are presented as mean ± SEM (n = 8–10 mice/group). Individual data points represent values from individual animals. *p < 0.05, ***p < 0.001, ****p < 0.0001.

## Discussion

This study provides the first experimental evidence that myotonia, independent of overt injury, inflammation, or systemic pathology, is sufficient to evoke persistent pain-like behavior. Both pharmacological and genetic disruption of ClC-1 function produced robust and prolonged hypersensitivity to mechanical, thermal, and cold stimuli, and mice with genetic myotonia congenita additionally exhibited spontaneous pain-like behavior. Notably, a single transient episode of pharmacologically induced myotonia generated pain-like behaviors that persisted beyond the resolution of overt motor symptoms. These behavioral changes were accompanied by altered sensory neuron excitability and peripheral nerve recruitment, selective changes in superficial dorsal horn neurons, and amplified activity in central nociceptive circuits. Finally, pharmacological inhibition of Na_V_1.8 markedly attenuated pain-like behavior in both pharmacological and genetic models of myotonia, supporting Na_V_1.8-directed analgesia as a promising therapeutic strategy for myotonia-associated pain. Together, these findings challenge the view of myotonia as a purely motor disorder and establish a link between skeletal muscle hyperexcitability and persistent alterations in nociceptive processing.

### Mechanistic Insights into Peripheral and Central Sensitization

ClC-1 governs membrane repolarization in myofibers, and its disruption leads to delayed relaxation, repetitive firing, and pathological excitability. Importantly, *Clcn1* is not expressed in primary sensory neurons under either physiological or pathological conditions (87), making it unlikely that the sensory changes observed here result from direct disruption of ClC-1 within DRG neurons. Instead, pathological muscle activity may secondarily alter sensory input from affected muscle (**Figure 13**). Repetitive contraction and muscle membrane hyperexcitability could increase activation of nociceptive muscle afferents, while the reduced A-fiber recruitment observed following transient myotonia raises the possibility that low-threshold sensory signaling is also altered. Together, increased nociceptive drive and reduced low-threshold afferent input could favor the development of peripheral and central sensitization (88, 89).

**Figure 13.**
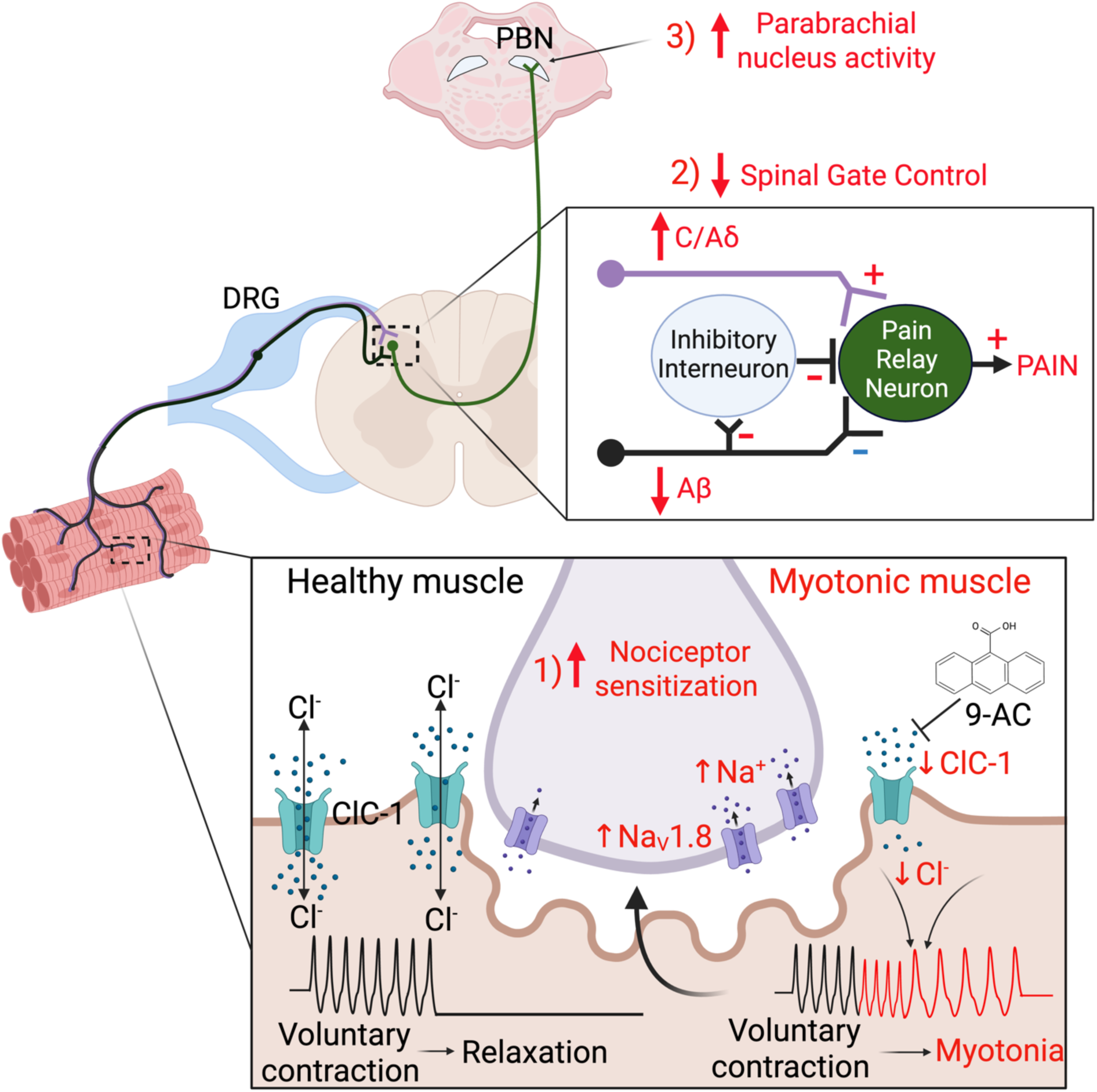
Schematic illustration of the proposed mechanisms by which myotonia leads to pain. The ClC-1 chloride channel is essential for stabilizing skeletal muscle membrane excitability and promoting efficient repolarization following voluntary contraction. In myotonia, loss or dysfunction of ClC-1 reduces chloride conductance, delays membrane repolarization, and produces repetitive muscle action potential firing and sustained involuntary contractions. We propose that pathological muscle activity initiates a cascade of neuroplastic changes throughout the pain neuraxis. (1) Repetitive myotonic discharges promote sensitization of muscle-innervating primary afferents, potentially through increased membrane expression and activity of the nociceptor-enriched sodium channel Na_V_1.8. (2) Altered peripheral input, characterized by enhanced nociceptive signaling and reduced recruitment of large-diameter myelinated afferents, disrupts spinal gate control and facilitates transmission through spinal pain relay circuits. (3) These changes culminate in enhanced sensory-evoked activity within the parabrachial nucleus, a major brainstem hub for nociceptive processing and affective pain signaling. Together, these findings support a model in which myotonia is sufficient to drive persistent pain through coordinated peripheral and central sensitization mechanisms.

Several observations support this model. First, pain-like behaviors persisted beyond the resolution of overt myotonic motor symptoms. This temporal dissociation suggests that a transient period of pathological muscle activity can produce persistent changes within nociceptive pathways rather than pain-like behavior simply tracking overt motor dysfunction. Second, increased action potential firing was restricted to small- and medium-diameter sensory neurons, whereas large-diameter sensory neurons remained largely unaffected. This finding is consistent with preferential sensitization of putative nociceptive afferents and argues against generalized dysfunction of all sensory neurons. Finally, voltage-clamp recordings revealed a numerical increase in tetrodotoxin-resistant (TTX-R) sodium current following transient myotonia, although this difference did not reach statistical significance. TTX-R currents in nociceptors are carried predominantly by Na_V_1.8 and Na_V_1.9, with Na_V_1.8 playing an important role in action potential generation and repetitive firing (82, 90, 91). Pharmacological inhibition of Na_V_1.8 with Suzetrigine, a recently United States Food and Drug Administration (FDA)-approved Na_V_1.8 inhibitor (90, 92, 93), reduced firing in sensory neurons and markedly attenuated pain-like behavior in both transient and genetic myotonia. However, Suzetrigine also reduced firing in control neurons, and the electrophysiological data therefore do not establish increased Na_V_1.8 activity as the primary mechanism responsible for post-myotonia hyperexcitability. Rather, considered together with the behavioral efficacy of Suzetrigine, these findings support Na_V_1.8-directed analgesia as a promising strategy for modulating pain associated with myotonia. The high prevalence of Na_V_1.8 expression among muscle-innervating nociceptors provides a plausible anatomical substrate for the pronounced behavioral efficacy of Na_V_1.8 inhibition (94, 95), although direct identification of the afferent populations sensitized by myotonia will require future study.

One of the more unexpected findings of the present study was the reduction in A-fiber recruitment following transient myotonia. Although conduction velocity and activity-dependent suppression were preserved, maximal recruitment was reduced, and higher stimulation intensities were required to achieve half-maximal activation. At present, the mechanism underlying this phenomenon remains unclear. Nevertheless, the finding is intriguing because it aligns with both established pain theory and clinical observations in myotonic disorders. One possibility is that reduced A-fiber recruitment diminishes inhibitory gating of spinal nociceptive transmission. According to gate control theory, Aβ fibers exert inhibitory control over spinal nociceptive transmission by activating local spinal inhibitory interneurons (96). Disruption of this input could remove an important brake, allowing nociceptive signals to dominate and promoting vulnerability to central sensitization (97). Such a mechanism may be particularly relevant in the context of myotonia, where persistent nociceptor activity and diminished low-threshold sensory input could act synergistically to amplify pain processing (**Figure 13**). This possibility is consistent with reports of altered sensory nerve function, reduced sensory nerve action potentials, proprioceptive deficits, reduced intraepidermal nerve fiber density, and impaired vibration sensation in patients with ClC-1-associated disorders (54, 61, 98–107). Although their relationship to pain remains poorly understood, these observations raise the possibility that impaired low-threshold afferent signaling contributes to pain in myotonic disorders.

Evidence consistent with altered central nociceptive processing was observed at both spinal and supraspinal levels. Broad analysis of superficial dorsal horn neurons revealed relatively modest changes following transient myotonia. However, stratification by firing phenotype uncovered increased excitability specifically within delay firing neurons. This finding is notable because delay firing neurons within lamina II are predominantly excitatory glutamatergic interneurons that have been implicated in the amplification and transmission of nociceptive information during pathological pain states (108). Increased excitability of this population could enhance ascending nociceptive output from the spinal cord and may therefore contribute to the robust enhancement of sensory-evoked activity observed in the parabrachial nucleus (79). The PBN is increasingly recognized as a critical hub for integrating sensory and affective dimensions of pain and exhibits heightened activity across numerous chronic pain conditions (79, 109). Accordingly, the exaggerated PBN responses to innocuous mechanical and thermal stimuli observed here suggest that transient myotonia engages central amplification mechanisms extending beyond the spinal cord and into higher-order pain-processing networks.

### Implications for Nociplastic Pain

These observations fit within the emerging conceptual framework of nociplastic pain, a category proposed by the International Association for the Study of Pain to describe pain arising from altered nociceptive processing in the absence of identifiable tissue or nerve injury (110). Nociplastic pain commonly presents as diffuse pain accompanied by fatigue, sleep disturbances, and mood disorders. The symptom profile of myotonic disorders and the changes in nociceptive signaling described in the present study share features with nociplastic pain, including persistent hypersensitivity and altered peripheral and central nociceptive processing in the absence of overt tissue or nervous system injury (111–114). Indeed, patients with myotonia congenita and other myotonic disorders frequently report diffuse, treatment-resistant pain, and a substantial proportion are misdiagnosed with fibromyalgia or meet its diagnostic criteria (46, 48, 115, 116). Particularly striking was the observation that a transient episode of myotonia produced prolonged hypersensitivity that persisted after overt motor symptoms had resolved. Together, these findings raise the possibility that recurrent skeletal muscle hyperexcitability can initiate persistent alterations in nociceptive processing with features consistent with nociplastic pain.

### Translational Potential of Targeting Na_V_1.8 for Myotonia-induced Pain

An additional and clinically salient finding of the present study is that pharmacological inhibition of Na_V_1.8 markedly attenuated pain-like behaviors in both pharmacological and genetic models of myotonia. The sensory electrophysiological phenotypes underlying these pain states, however, differ between models. In the transient 9-AC model, enhanced repetitive firing accompanied by a numerical, but nonsignificant, increase in TTX-resistant sodium current is consistent with altered nociceptor sodium channel function; however, these data do not establish increased Na_V_1.8 activity as the mechanism responsible for hyperexcitability. In contrast, sensory neurons from *Clcn1*^adr/adr^ mice did not exhibit increased repetitive firing despite robust pain-like behavior. Rather, these neurons displayed a depolarized resting membrane potential and alterations in action potential waveform properties, including increased half-width and altered decay kinetics. These findings suggest broader electrophysiological remodeling that is unlikely to be explained solely by Na_V_1.8. Changes in persistent sodium or repolarizing potassium conductances represent plausible contributors, but defining these mechanisms will require direct voltage-clamp analysis in the genetic model. Nevertheless, the robust behavioral efficacy of Suzetrigine supports Na_V_1.8-directed analgesia as a promising therapeutic strategy despite differences in the underlying electrophysiological phenotypes.

From a translational perspective, these observations are particularly intriguing given the recent FDA approval of the selective Na_V_1.8 inhibitor Suzetrigine (Journavx®, VX-548) for the treatment of acute pain (93). At present, there are no targeted therapies for pain associated with myotonic disorders, and evidence supporting pharmacological management of myotonic pain remains remarkably limited. To our knowledge, only four drugs, mexiletine, lamotrigine, ranolazine, and carbamazepine, have demonstrated efficacy in reducing pain in patients with myotonic disorders. Notably, all four compounds inhibit voltage-gated sodium channels. Mexiletine, the current first-line treatment for myotonia, reduces pain severity in patients with myotonia congenita and non-dystrophic myotonia (117, 118). Interestingly, mexiletine inhibits tetrodotoxin-resistant sodium currents in rat DRG neurons in both a state- and use-dependent manner (119). Similarly, the anticonvulsant lamotrigine exhibits anti-myotonic efficacy comparable to mexiletine and produces modest improvements in pain (118). In heterologous expression systems, lamotrigine and several of its derivatives inhibit both TTX-R and Na_V_1.8 (120). In contrast, open-label studies of ranolazine have demonstrated robust reductions in pain severity in patients with myotonia congenita (121) and paramyotonia congenita (122). Although originally developed as an antianginal agent and subsequently repurposed as an anti-myotonic therapy, ranolazine inhibits multiple sodium channel subtypes, including Na_V_1.8 (123). Likewise, approximately half of patients with non-dystrophic myotonia report improvements in pain during treatment with carbamazepine (99), another sodium channel inhibitor with activity at Na_V_1.8 (124). Although these clinical observations are indirect, together with our findings they raise the possibility that inhibition of nociceptor sodium channels contributes to the analgesic effects of existing sodium channel-blocking therapies in myotonic disorders. Importantly, several of these agents also inhibit skeletal muscle Na_V_1.4 and may therefore reduce both the initiating myotonia and downstream nociceptive signaling. Our results identify selective Na_V_1.8 inhibition as a promising analgesic strategy that warrants further investigation for myotonia-associated pain, while broader sodium channel blockers may offer complementary therapeutic advantages by acting at both muscle and sensory-neuron targets.

### Future Directions

Several questions emerge from these findings. First, studies in patients using quantitative sensory testing, microneurography, and electrophysiological measures could determine whether the peripheral sensory abnormalities identified here translate to human myotonic disorders. Second, direct analysis of muscle-innervating afferents will be necessary to establish which sensory populations are sensitized and how pathological muscle activity produces persistent neuronal plasticity. The reduction in A-fiber recruitment following transient myotonia also warrants investigation in genetic models to determine whether altered low-threshold sensory input is a conserved feature of chronic myotonia. Finally, future studies should define the spinal and supraspinal circuits that maintain pain after myotonic episodes and determine whether recurrent muscle hyperexcitability produces durable changes in central pain-processing networks.

### Limitations and Conclusions

Several limitations should be considered when interpreting the present findings. First, we intentionally simplified the myotonic disorder experience by focusing on models resulting from ClC-1 dysfunction. By contrast, patients with myotonic disorders frequently exhibit additional neurological, muscular, and systemic abnormalities that may further shape their pain experience. This reductionist approach nevertheless allowed us to isolate the contribution of myotonic muscle dysfunction to pain-like behavior, a relationship that had not previously been demonstrated experimentally. Second, because muscle stiffness and delayed movement can influence withdrawal responses, reflexive assays cannot completely dissociate altered nociception from motor impairment. Third, the duration of 9-AC activity *in vivo* was inferred from recovery of overt motor function rather than direct measurement of muscle ClC-1 conductance or electromyographic myotonia at later time points. Although righting reflex impairment resolved within approximately 3 hours, this assay may not detect milder residual myotonic activity in individual muscle fibers. Therefore, while the persistence of pain-like behavior beyond the period of overt motor impairment suggests a dissociation between gross myotonia and nociceptive hypersensitivity, we cannot exclude the possibility that subclinical muscle hyperexcitability persists beyond behavioral recovery. Future studies using longitudinal EMG recordings or direct assessment of muscle excitability will be important to define the precise temporal relationship between myotonic activity, sensory neuron sensitization, and pain-like behavior. Finally, we focused exclusively on ClC-1 dysfunction and did not examine painful myotonic disorders caused by mutations in *SCN4A* or other genes. Whether similar mechanisms contribute to pain across genetically distinct myotonic syndromes remains an important question for future investigation.

An important caveat of our electrophysiological studies is that acute Suzetrigine application reduced repetitive firing in sensory neurons from both control and myotonic mice. In contrast, the behavioral effects of Suzetrigine were highly selective, producing robust analgesia in myotonic animals while exerting minimal effects in vehicle-treated or wildtype mice. One possible explanation is concentration-dependent off-target actions that become apparent during acute *in vitro* recordings but are less evident behaviorally *in vivo*, where Suzetrigine produces robust analgesia in myotonic animals with minimal effects in controls. Suzetrigine was developed as a highly selective inhibitor of human Na_V_1.8, where it inhibits the channel with subnanomolar potency (IC_50_ ≈ 0.68 nM) (92). However, native mouse Na_V_1.8 currents are approximately 500-fold less sensitive (IC_50_ ≈ 0.21-0.35 μM), necessitating micromolar concentrations to achieve near-complete channel inhibition in mouse electrophysiological experiments (83, 92). At these concentrations, Suzetrigine is no longer functionally selective in mouse neurons. Recent work demonstrated that the concentrations required to fully inhibit mouse Na_V_1.8 also inhibit voltage-gated potassium channels, producing smaller and narrower action potentials and altering repetitive firing independent of Na_V_1.8 blockade (125). Thus, the acute electrophysiological effects of Suzetrigine should be interpreted cautiously, whereas its robust behavioral efficacy in myotonic animals supports further investigation of Na_V_1.8-directed analgesia for myotonia-associated pain.

In conclusion, this work challenges the long-standing assumption that myotonia is a purely motor disorder and establishes a physiological link between skeletal muscle hyperexcitability and persistent alterations in nociceptive processing. Our findings demonstrate that pain-like behavior can persist beyond overt myotonic motor dysfunction and is accompanied by changes in peripheral sensory neurons, spinal neurons, and supraspinal nociceptive activity. Pharmacological inhibition of Na_V_1.8 robustly alleviated pain-like behavior in both pharmacological and genetic models, supporting Na_V_1.8-directed analgesia as a promising therapeutic strategy while revealing that the sensory electrophysiological consequences of myotonia differ between acute and chronic disease models. More broadly, these findings establish a framework in which pathological skeletal muscle activity can initiate persistent remodeling of nociceptive pathways and provide a foundation for mechanism-based treatment of pain in myotonic disorders.

## Methods

Adult male and female mice were used throughout. Animals in pharmacological studies were randomized to treatment groups and investigators were blinded to treatment during data collection and analysis; genotype masking procedures for genetic studies are detailed in the **Supplementary Material**. Pain-related and motor phenotypes were assessed using established behavioral assays measuring mechanical, dynamic mechanical, cold, and thermal sensitivity, grip strength, and righting reflex. Peripheral and spinal neuronal function was examined using whole-cell patch-clamp electrophysiology of dissociated dorsal root ganglion neurons and superficial dorsal horn neurons, while *ex vivo* sciatic nerve compound action potential recordings assessed peripheral nerve function. *In vivo* fiber photometry was used to measure sensory-evoked activity of glutamatergic parabrachial nucleus neurons before and after induction of myotonia. Detailed animal procedures, drug preparation and administration, electrophysiological protocols, behavioral assays, fiber photometry, blinding and randomization procedures, and statistical analyses are provided in the **Supplementary Material**.

### Study Approval

All procedures were consistent with the guidelines for the treatment of animals of the International Association for the Study of Pain and were approved by the University of Florida’s and the University of Pittsburgh’s Institutional Animal Care and Use Committee.

### Data Availability

All data supporting the findings of this study are available from the corresponding author upon reasonable request. The data plotted in all figures can be found in the **Supporting data values** file. No custom code was generated.

## Author Contributions

TSN conceived the project, designed and performed experiments, and wrote the manuscript. ACR led the DRG electrophysiological recordings in dorsal root ganglion neurons. HNA designed and conducted the fiber photometry studies. ELA, JBPF, and NP performed compound action potential recordings and analysis. AW, KG, and SLL contributed to DRG electrophysiology experiments. EJRP assisted with creation of the summary schematic. PD contributed to data acquisition. MSG supervised the compound action potential recording studies and provided critical input on data interpretation. RK directed the overall project, secured funding, and provided scientific oversight. All authors contributed to manuscript editing and approved the final version.

## Funding Support

-National Institutes of Health, K00NS124190, TSN

-National Institutes of Health, F32NS128392, HNA

-National Institutes of Health, K99NS134965, KG

-National Institutes of Health, RF1NS131165, RK

-National Institutes of Health, R61NS126026, RK

-National Institutes of Health, R01NS120663, RK

-American Neuromuscular Foundation, Development Grant, TSN

-PhRMA Foundation, Postdoctoral Fellowship #1335819, EJRP

This work is the result of NIH funding and is subject to the NIH Public Access Policy. Through acceptance of this federal funding, the NIH has been given a right to make the work publicly available in PubMed Central.

## Acknowledgements

We thank Dr. Mark Rich for providing the *Clcn1*^adr/adr^ mice and Dr. Dan Wang for assistance performing the spinal cord dorsal horn slice electrophysiological recordings.

