## Supplementary Material for "Myotonic Muscle Activity Drives a Persistent Pain State Responsive to Na_V_1.8-directed Analgesics"

**Supplementary Methods:**

Animals

Adult C57Bl/6J (Jackson Laboratory, # 000664), and SWR/J-*Clcn1*$^{adr\text{-}mto}$/J (Jackson Laboratory, # 000939) male and female mice were group housed, provided access to food and water *ad libitum*, and maintained on a 12:12 hour light:dark cycle in temperature and humidity-controlled rooms.

Sex as a Biological Variable

Equal numbers of male and female mice were used in all experiments. Although we were not powered to detect significant sex differences, no major/obvious trends in sex differences were observed and means from both sexes were pooled.

Drug Administration

Anthracene-9-carboxylic acid, 99% purity (Thermo Scientific, Cat# 104880100), was diluted in a solution containing 50% dimethyl sulfoxide and 50% sterile 0.9% sodium chloride, while the vehicle comprised the same constituents. The drug was freshly prepared each day just before administration due to its tendency to precipitate out of solution rapidly. Suzetrigine (MedChemExpress, Cat# HY-148800) was diluted in a solution containing 10% dimethyl sulfoxide, 10% Tween 80, and 80% sterile 0.9% sodium chloride, while the vehicle comprised the same constituents. Mice were briefly scruffed and received a single intraperitoneal (i.p.) injection using a 30-gauge needle.

Electrophysiological Recordings in Peripheral Sensory Neurons

Whole-cell patch-clamp recordings were performed in dissociated dorsal root ganglion (DRG) neurons, following protocols previously established by our group (1, 2). Lumbar DRGs were harvested from adult male and female mice (9 weeks to 3 months old). For 9-AC recordings, DRG isolation occurred either 6 hours (for recordings at 16-24 hours post-9-AC injection) or 24 hours (for recordings at ~48 hours post-injection) after administration of 9-AC (30 mg/kg, i.p.). DRG tissues were enzymatically dissociated at 37°C for 50 minutes in Dulbecco's Modified Eagle Medium (DMEM) containing collagenase type I (1.66 mg/mL, Cat# LS004194; Worthington) and neutral protease (1.04 mg/mL, Cat# LS02104; Worthington). Following centrifugation (800 rpm, 5 minutes), dissociated neurons were plated onto 12-mm poly-D-lysine-coated coverslips (0.1 mg/mL, Cat# P6407; Millipore Sigma) and maintained in culture medium supplemented with 10% fetal bovine serum (HyClone) and 1% penicillin/streptomycin (Cat# 15140; Life Technologies).

Current Clamp: Electrophysiological recordings were conducted 15-24 hours after plating using the whole-cell configuration of the patch-clamp technique in current-clamp mode. Recordings were performed at room temperature (20-22°C) using an EPC 10 amplifier (HEKA Elektronik, Germany) and acquired with PatchMaster. Data were analyzed using FitMaster and Easy Electrophysiology software. Borosilicate glass pipettes (resistance ~2-3.5 MΩ) were filled with internal solution (see **Supplemental Table 1**) and giga-ohm seals were formed with DRG neurons. After achieving the whole-cell configuration under voltage clamp, recordings were switched to current-clamp mode to assess neuronal excitability. Neurons were categorized as small (<15 pF and diameter under 23 μm) or large (>40 pF and diameter ~37 μm) based on measured whole-cell capacitance and approximate size. In small-diameter neurons (capacitance <15 pF), action potentials (APs) were elicited with a depolarizing ramp protocol (0-150 or 0-250 pA over 1 second), and rheobase was calculated as the minimal current required to evoke an AP. In contrast, large-diameter neurons (average capacitance ~45 pF) were not amenable to ramp protocols due to intrinsic membrane properties. Specifically, their larger capacitance and differential channel expression reduce the rate of membrane depolarization during ramps, leading to depolarization-induced inactivation and failure to generate APs. Therefore, in large neurons, rheobase was determined using step current injections starting at 0 pA and increasing in 100 pA increments (1-second duration per step). Input resistance was estimated in current-clamp mode by measuring the deflection in the membrane potential ($\Delta V$) in response to a small hyperpolarizing current pulse ($\Delta I$), by using Ohm's law.

$$R_{in} = \frac{\Delta V}{\Delta I}$$

Voltage Clamp: Electrophysiological recordings were conducted ~15 hours after plating using the whole-cell configuration of the patch-clamp technique in voltage-clamp mode. Briefly, total sodium currents were evoked by 150-ms voltage steps from −70 to +50 mV in 10-mV increments from a holding potential of −100 mV.

Solutions for electrophysiological recordings are detailed in **Supplemental Table 1**.

Electrophysiological Recordings in Spinal Dorsal Horn Neurons

Lumbar spinal cord slices were prepared from adult male and female mice. Following deep isoflurane anesthesia and transcardial perfusion with ice-cold oxygenated artificial cerebrospinal fluid (ACSF), the lumbar spinal cord was rapidly removed and sectioned into transverse slices. Individual slices were transferred to a submerged recording chamber mounted on an upright microscope (ECLIPSE FN1, Nikon, Japan) and continuously superfused with oxygenated ACSF at a rate of 2 mL/min at room temperature (20–22°C). Neurons located within lamina I and lamina II of the superficial dorsal horn were visualized using differential interference contrast optics. Patch electrodes (5–8 MΩ) were fabricated from borosilicate glass and filled with an intracellular potassium-based solution containing (in mM): 130 K-gluconate, 1 $MgCl_2$, 1 $CaCl_2$, 1 KCl, 10 HEPES, 11 EGTA, 2 Mg-ATP, and 0.3 Na-GTP (pH 7.3; 290 mOsm). Signals were low-pass filtered at 3 kHz and digitized at 10 kHz using an EPC-10 amplifier controlled by PatchMaster software (HEKA Elektronik, Lambrecht, Germany). Resting membrane potential (RMP) was recorded shortly after establishing the whole-cell configuration. Cells with membrane potentials more depolarized than −40 mV were excluded from analysis. The liquid junction potential (approximately −13 mV) was not corrected. Series resistance was continuously monitored throughout recordings and maintained below 30 MΩ. Passive membrane properties were assessed under voltage-clamp conditions by holding neurons at −70 mV and applying a brief hyperpolarizing step to −80 mV. Voltage-gated membrane currents were evaluated using a series of voltage steps ranging from −90 mV to +40 mV in 10 mV increments (500 ms duration per step). Neuronal excitability was assessed in current-clamp mode at the resting membrane potential. Action potentials were evoked using 500 ms current injections ranging from −20 pA to +200 pA in 20 pA increments. Rheobase was determined using 100 ms current injections delivered in 2 pA increments and was defined as the minimal current required to evoke an action potential. Neurons were classified according to their firing patterns as single, phasic, tonic, or delay firing based on responses to depolarizing current injection. Action potential kinetics were analyzed using Easy Electrophysiology software (version 2.8.0; https://github.com/RoyVII/EasyElectrophysiology).

Compound Action Potentials

For compound action potential (CAP) recording, sciatic nerves, from the trifurcation to the iliac crest, were removed bilaterally after mice had been deeply anesthetized with isoflurane and perfused transcardially with ice cold 0.1 M phosphate buffered saline. Nerves were placed in oxygenated Krebs solution (136 NaCl, 5.6 KCl, 14.3 $NaHCO_3$, 1.2 $NaH_2PO_4$, 2.2 $CaCl_2$, 1.2 $MgCl_2$, and 11 glucose bubbled continuously with carbogen ($O_2$ 95%/$CO_2$ 5%) to achieve a pH ranging from 7.2 to 7.4.) and stored on ice until recording. Glass suction electrodes were used for both stimulation (at the distal end) and recording (at the proximal end), where an amplifier (A-M Systems, Inc Sequim WA) was used to deliver square pulses of 0.3 (A-wave) and 3 ms (C-wave) to the nerve. The CAP recording electrode was connected to a differential preamplifier (0.1-10 kHz; model DAM-80, WPI), and the recording was sampled at 20 kHz via a Molecular Devices Digidata (1440a) analog-to-digital converter and acquired and analyzed using pCLAMP version 10 for MS Windows (Molecular Devices, San Jose CA). After mounting the nerve in the recording chamber, it was given 30 minutes to settle prior to further study. Recordings were performed at room temperature.

For each nerve, the A-wave component of the CAP, representing fibers conducting at >6 m/s, was studied first. The threshold was defined as the current intensity required to evoke a deflection in the voltage trace >3x the baseline noise in the voltage trace (taken 10 seconds prior to stimulation). A recruitment curve was then generated by stimulating the nerve at 1.5, 2, 3, 4, 5, 10, and 20x threshold. A 10 second inter-stimulus interval was used. The nerve was then stimulated 20 times at 10x threshold intensity at 30 Hz and 100 Hz to assess activity-dependent changes in the waveform. The C-wave was then studied with the same criteria used to determine the threshold for the activation of fibers conducting at <2 m/s. A recruitment curve was then generated by stimulating the nerve at 1.5, 2, and 3x threshold.

Conduction velocity was estimated based on the length of the nerve divided by latency of the peak of the rectified A- and C-waves relative to the initiation of the stimulation artifact. The integral (Area under the curve, AUC) of the rectified waveform evoked in response to each stimulus was used to generate recruitment curves. The AUC of the waveform evoked with the $20^{th}$ pulse divided by that of the first pulse was used to estimate activity-dependent changes in the CAP A-wave. Recruitment curves for the A-wave of the CAP were fitted with a logistic equation, f(x) = (AUCmax*x^slope)/(x^slope+AUC50^slope) that enabled estimation of the maximal AUC (AUCmax), current intensity associated with the activation of a CAP 50% of maximal (AUC50), and the slope of the curve at the 50% point.

### Fiber Photometry Recordings

As previously described by our group (1, 3–5), adult male and female wild-type mice received unilateral stereotaxic injections of 300 nL AAV9-CaMKIIa-GCaMP6s-WPRE-SV40 (Addgene, Watertown, MA) into the right parabrachial nucleus (PBN) to selectively express the calcium indicator GCaMP6s in glutamatergic neurons. Stereotaxic coordinates targeting the PBN were anteroposterior (AP) −5.15 mm, mediolateral (ML) ±1.45 mm, and dorsoventral (DV) −3.45 mm from bregma. Viral infusions were performed using a Nanoject II Auto-Nanoliter Injector (Drummond) at a rate of 2 nL/sec, with a post-infusion dwell time of 5 minutes to minimize reflux. Immediately following viral delivery, a fiber optic cannula (RWD; 1.25 mm ferrule diameter, 200 µm core diameter, 0.37 numerical aperture) was chronically implanted above the injection site and secured with dental cement (Cat# 10-000-786, Stoelting, Wood Dale, IL). Mice were allowed to recover for 21 days to allow for robust viral expression. Prior to each recording session, animals were acclimated in acrylic chambers with wire mesh flooring for a minimum of one hour, with the fiber optic patch cord connected and the experimenter in the room. Real-time calcium signals were acquired using the FP3002 fiber photometry system (Neurophotometrics). The plantar surface of the left hindpaw was sequentially stimulated with four sensory stimuli: a 0.07 g von Frey filament (1-second application), a 1.0 g von Frey filament (1-second application), a 10 µL acetone drop, and a blunt pinprick. Each stimulus was presented three times at 2-minute intervals, and the three trials were averaged to represent each animal’s response. Calcium signals were analyzed using custom MATLAB scripts. The 470 nm GCaMP6s signal was normalized to the 405 nm isosbestic control channel to correct for motion artifacts and photobleaching. The change in fluorescence (ΔF/F) was calculated by subtracting the baseline signal (10 seconds prior to stimulus onset) from the peak response following stimulation. Area under the curve (AUC) was computed for the 10-second window following each stimulus. After baseline recordings, mice received a single intraperitoneal injection of vehicle or 9-AC (30 mg/kg). Recordings were repeated 24 hours later. A crossover design was employed whereby, following a one-month washout period, mice received the alternate treatment and underwent a second round of recordings. Upon completion of the study, animals were deeply anesthetized and transcardially perfused with ice-cold 1x PBS followed by 10% neutral buffered formalin (Cat# SF98-4, Fisher Scientific, Waltham, MA). Brains were extracted, post-fixed, cryoprotected, and coronally sectioned at 30 µm thickness using a cryostat. Sections were stored at 4°C until further processing. To verify GCaMP6s expression and fiber placement, immunohistochemistry for GFP was performed. Sections were washed three times in PBS, blocked in 5% normal goat serum (NGS) in PBS with 0.1% Triton X-100 for one hour, and incubated overnight at room temperature with rabbit anti-GFP (1:1000; Cat# AB3080, Millipore Sigma, St. Louis, MO) in blocking buffer. After washing, sections were incubated for 1.5 hours in goat anti-rabbit AlexaFluor 488 secondary antibody (Cat# A11008, Invitrogen, Waltham, MA), washed again, mounted on SuperFrost Plus microscope slides (Cat# 22-037-246, Fisher Scientific), and coverslipped using Vectashield Plus antifade mounting medium with DAPI (H-2000-10, Vector Laboratories). Images were acquired using a Leica DMi8 inverted widefield fluorescence microscope at 20x magnification. No animals were excluded based on post hoc verification of viral transfection or fiber placement.

### Behavioral Testing

*Static Mechanical Sensitivity (von Frey):* Testing was performed as previously described (1, 6–8). Mice were habituated to plexiglass chambers (80 × 80 × 110 mm) on a raised wire mesh platform for 60 minutes immediately prior to behavioral testing. Testing was performed using a calibrated set of logarithmically increasing von Frey monofilaments (Braintree Scientific Cat# 58011) that range in gram force from 0.007 to 6.0 g. Beginning with a 0.4 g filament, these were applied perpendicular to the lateral hindpaw surface with sufficient force to cause a slight bending of the filament. A positive response was denoted as a rapid withdrawal of the paw within 4 seconds of application and was followed by application of the next lower filament. A negative response was followed by application of the next higher filament. An up-down method (9) was used to calculate the 50% withdrawal threshold for each mouse.

*Dynamic Mechanical Sensitivity (Brush):* Testing was performed as previously described (8). Immediately following von Frey testing, dynamic mechanical sensitivity testing was performed on mice in the same plexiglass chambers on a raised wire mesh platform. A cotton swab was “puffed-out” to about three times its original size. The left hindpaw was then brushed using this cotton swab in the heel-to-toe direction. A positive response was noted as a rapid withdrawal of the paw in response to stimulation. The test was repeated four times, and the frequency of responses was reported.

*Cool Withdrawal Duration (Acetone):* Testing was performed as previously described (1, 6, 7). Immediately following dynamic brush testing, sensitivity to non-noxious cold testing was performed on mice in the same plexiglass chambers on a raised wire mesh platform. Using a syringe connected to PE-90 tubing, flared at the tip to a diameter of 3 1/2 mm, we applied a drop of acetone (VWR Cat# BDH1101-1LP) to the plantar surface of the hind left paw. Surface tension maintained the volume of the drop to ~10 $\mu$L. The duration of time the animal lifted or shook its paw was recorded for 30 seconds. Three observations were averaged.

*Time to Righting Reflex (TRR)*: Mice were gently placed on their backs on a flat surface to initiate the test. The duration it took for the mouse to return to an upright position, termed the righting reflex, was recorded. Each trial was repeated three times at each time point, and the average of these observations was calculated. This reflex serves as an indicator of the mouse's motor coordination and recovery capability. In healthy mice, the time to right for all animals was less than 0.5 seconds making variability smaller than could be plotted.

*Grip Strength Test*: Forelimb grip strength was assessed using the Grip Strength Test (MazeEngineers). Mice were allowed to grasp the metal grid of the apparatus with their forepaws and were then gently and steadily pulled backward by the base of the tail until they released the grid. The peak force exerted immediately prior to release was automatically recorded by the apparatus. Each mouse underwent three consecutive trials at each time point, and the average of these measurements was used for analysis.

*Thermal Withdrawal Threshold (Hot Plate):* We evaluated acute nociceptive responses with a hot plate as previously described (7, 10, 11). Briefly, prior to testing, mice were acclimated to the testing room for at least 30 minutes to minimize stress-induced behaviors. Following acclimation, mice were gently placed on a 52.5 °C Ugo Basile hot/cold plate (Stoelting Cat:55075). The trial ended when the mouse exhibited a nociceptive response (such as paw licking or jumping) or after a cutoff time of 30 seconds if no response was observed. Testing was repeated for 1-3 trials per mouse per temperature with intervals of at least 10 minutes between each trial. The latency to respond for each trial was averaged to produce the mean latency for statistical analysis.

### Blinding and Randomization

Animals in pharmacological studies were randomly assigned to treatment groups, and experimenters were blinded to treatment allocation during behavioral testing, fiber photometry recordings, and electrophysiological experiments. For studies using the *Clcn1*$^{adr/adr}$ model, formal genotype information was withheld from experimenters during behavioral testing, electrophysiological recordings, and data analysis. However, complete blinding to genotype was not possible because homozygous *Clcn1*$^{adr/adr}$ mice exhibit readily apparent phenotypic characteristics, including altered body size, coat appearance, and myotonic motor behavior. Accordingly, these experiments should be considered genotype-masked where feasible rather than fully blinded. Data analysis was performed without access to treatment or genotype assignments whenever feasible, with group identities revealed after completion of the relevant analyses.

### Statistics

All data are expressed as mean ± standard error of the mean (SEM), with individual data points representing either single animals (behavioral and fiber photometry experiments) or individual neurons (electrophysiological experiments). Statistical analyses were performed using GraphPad Prism (v10.2.3).

Comparisons between two groups were performed using unpaired two-tailed t tests with Welch's correction for unequal variances or Mann-Whitney U tests for nonparametric data, as appropriate. Comparisons involving more than two groups were analyzed using one-way ANOVA followed by either Tukey's or Dunnett's multiple comparisons tests. Factorial experiments were analyzed using two-way ANOVA, two-way repeated-measures ANOVA, three-way repeated-measures ANOVA, or mixed-effects models, as appropriate for the experimental design and presence of repeated measurements. When omnibus analyses revealed a significant interaction effect, prespecified post hoc multiple comparisons were performed using Tukey's, Holm-Šidák's, Dunnett's, or uncorrected Fisher's least significant difference (LSD) tests. Area under the curve (AUC) values were calculated using the trapezoidal rule. Statistical significance was defined as $p < 0.05$. No formal power analyses were performed. Sample sizes were determined based on prior experience with similar behavioral and electrophysiological assays. Group sizes, exact n values, and statistical tests used for each experiment are provided in the figure legends and **Supplemental Table 2**.

Graphics

Figures were generated using GraphPad Prism (v10.2.3), Adobe Illustrator 2022, and Biorender.com.

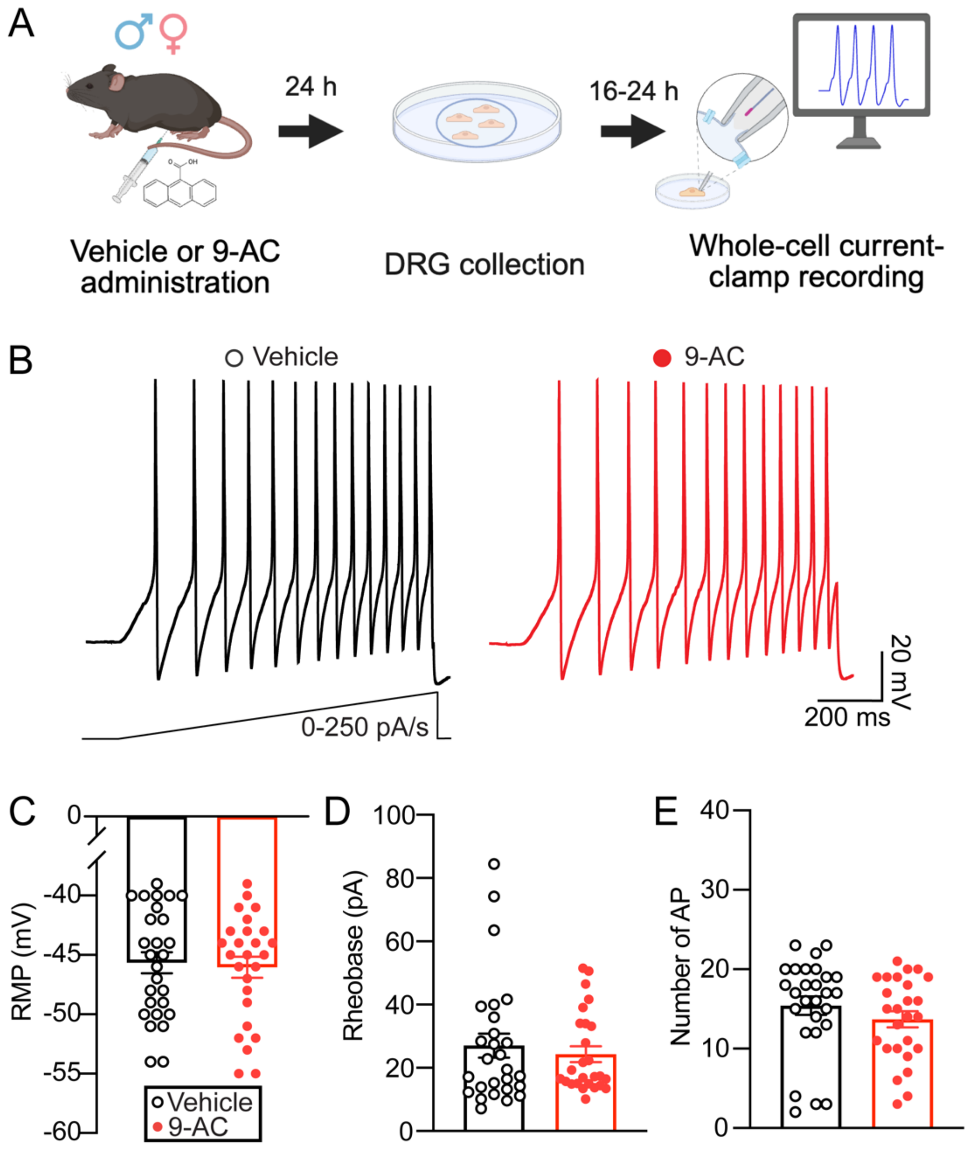


**Supplemental Figure 1. 9-AC does not increase DRG neuron excitability 48 hours post-injection.**

**(A)** Schematic illustrating the experimental timeline. Mice received a single intraperitoneal injection of 9-AC (30 mg/kg), followed by lumbar DRG dissociation 24 hours later. Whole-cell current-clamp recordings were performed in small- and medium-diameter DRG neurons 48 hours post-injection. **(B)** Representative traces showing action potential (AP) firing in sensory neurons from vehicle-treated (left) and 9-AC-treated (right) animals in response to suprathreshold current injection. **(C-E)** Quantification of (**C**) resting membrane potential, (**D**) rheobase, and (**E**) number of evoked APs. (n=26-27 neurons). Data were analyzed using the Mann-Whitney U test. Data are presented as mean ± SEM. Individual data points represent values from single neurons.

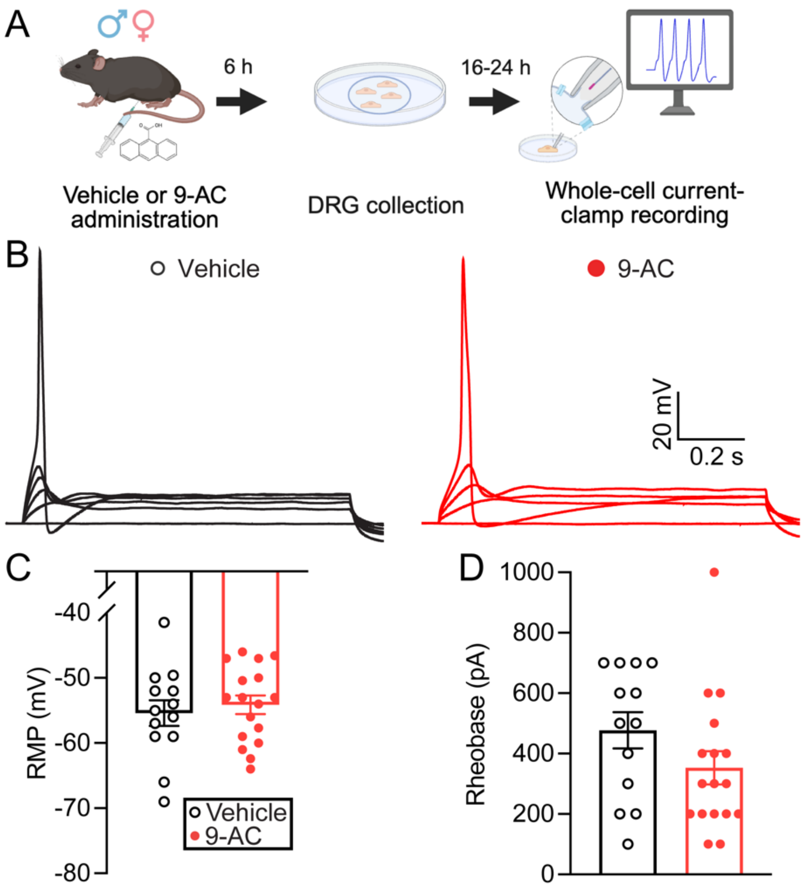


**Supplemental Figure 2. Rheobase is not altered in large-diameter DRG neurons after 9-AC administration. (A)** Schematic illustrating the experimental timeline. Mice received a single intraperitoneal injection of 9-AC (30 mg/kg), followed by lumbar DRG dissociation 6 hours later. Whole-cell current-clamp recordings were performed in large-diameter DRG neurons 24 hours post-injection. **(B)** Representative traces of current-clamp recordings showing rheobase in large-diameter DRG neurons from vehicle-treated and 9-AC-treated mice. **(C-D)** Quantification of **(C)** resting membrane potential and **(D)** rheobase, measured in 100 pA increments. Data were analyzed using the Mann-Whitney U test. Data are presented as mean ± SEM (n = 13-17 neurons). Individual data points represent values from single neurons.

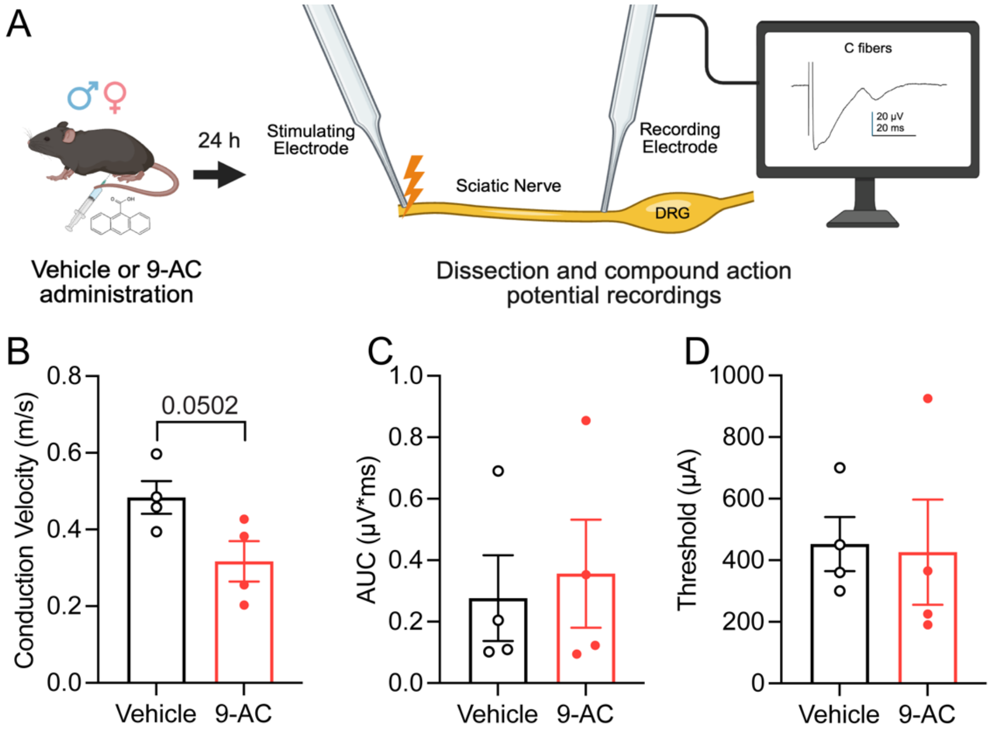


**Supplemental Figure 3. 9-AC does not alter C-fiber compound action potential responses. (A)** Schematic illustrating the experimental timeline. Mice received a single intraperitoneal injection of 9-AC (30 mg/kg), followed by sciatic nerve dissection 24 hours later. Compound action potential recordings were performed to assess C-fiber function. Quantification of (**B**) conduction velocity, (**C**) AUC, and (**D**) threshold. Data in panels **C-D** were analyzed using the Welch's T test. Data are presented as mean ± SEM (n = 4 animals/group). Individual data points represent values from single animals (average of 2 nerves/animal).

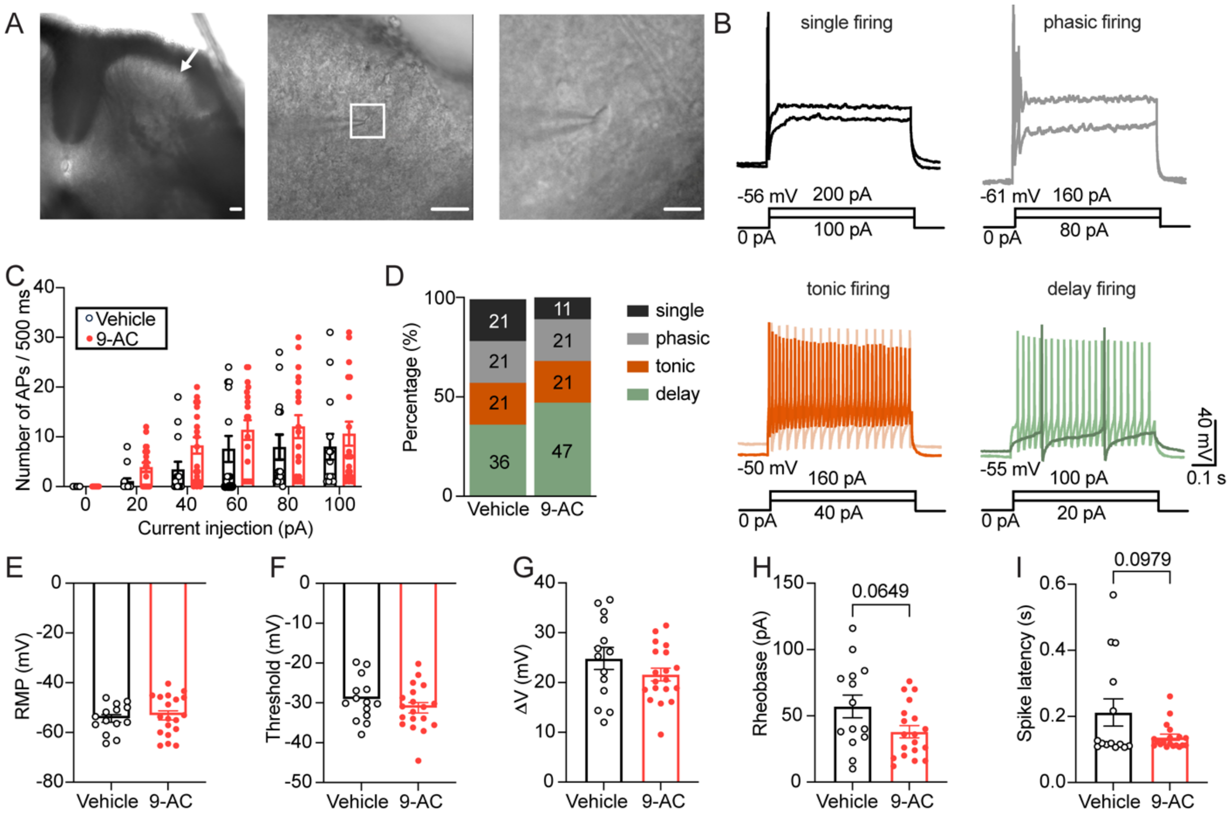


**Supplemental Figure 4. 9-AC does not broadly alter the excitability, intrinsic membrane properties, or firing-pattern distribution of superficial dorsal horn neurons. (A)** Representative coronal spinal cord sections showing the superficial dorsal horn (laminae I-II, white arrow) and a recorded neuron at progressively higher magnifications. Scale bars: 100 μm (left), 50 μm (middle), and 10 μm (right). **(B)** Representative traces illustrating the four firing patterns observed in superficial dorsal horn neurons: single, phasic, tonic, and delay firing. Corresponding current injection protocols are shown below each trace. **(C)** Frequency-current **(F-I)** relationship demonstrating the number of evoked action potentials (APs) in response to increasing depolarizing current injections in neurons from vehicle- and 9-AC-treated mice. **(D)** Distribution of neuronal firing patterns in vehicle- and 9-AC-treated groups. **(E-I)** Quantification of **(E)** resting membrane potential (RMP), **(F)** action potential threshold, **(G)** threshold distance (ΔV; VThreshold − Vrest), **(H)** rheobase, and **(I)** spike latency. **(C)** Data were analyzed using two-way repeated-measures ANOVA. **(D)** Firing pattern distributions were compared using Fisher's exact test. **(E-I)** Electrophysiological parameters were analyzed using unpaired two-tailed Student's t tests with Welch's correction. Data are presented as mean ± SEM (n = 14-19 neurons). Individual data points represent values from single neurons. Exact p values are indicated on the graphs. In some instances, error bars are too small to be visually discernible.

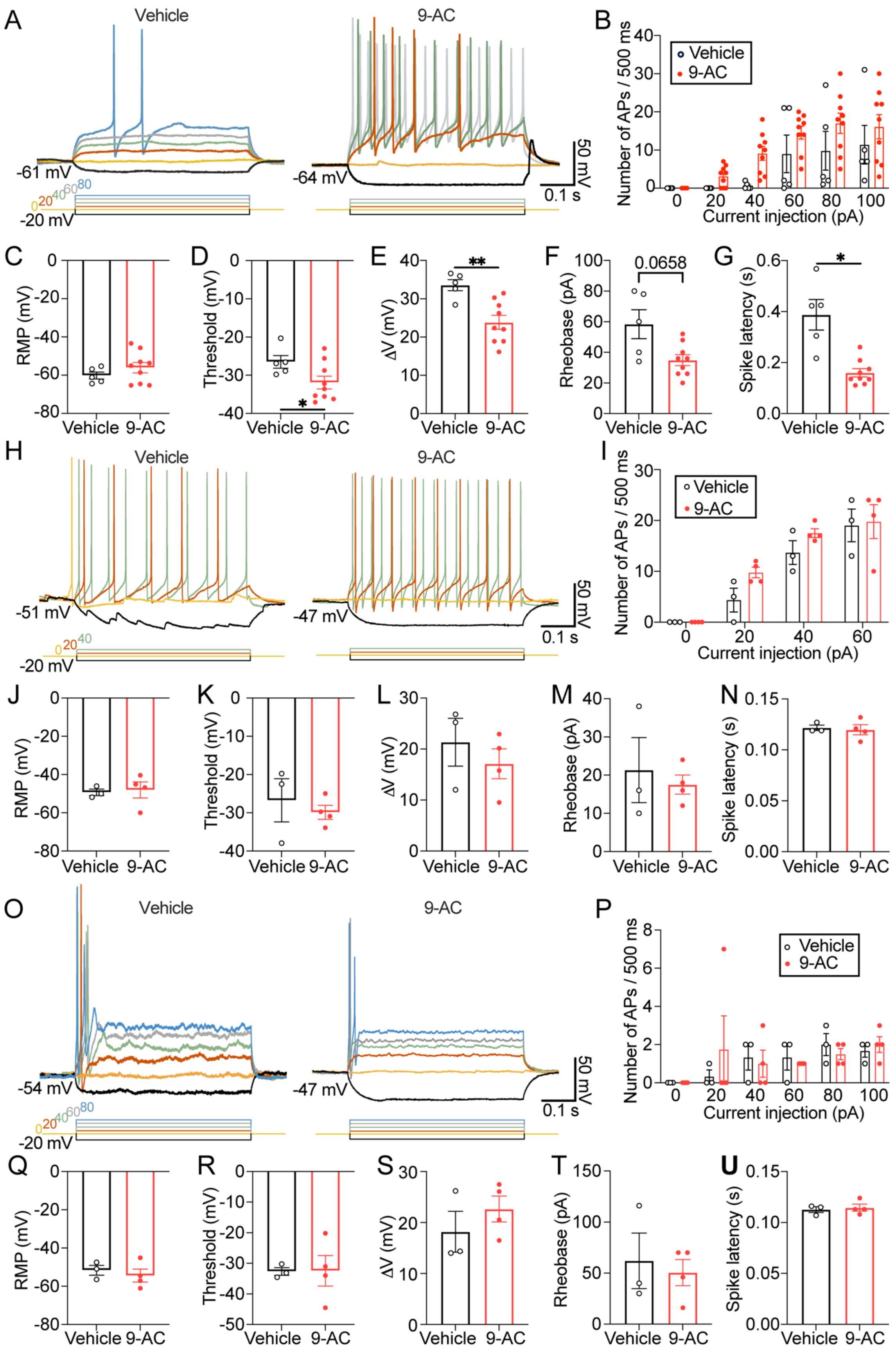


**Supplemental Figure 5. Decomposition of superficial dorsal horn neurons by firing pattern reveals enhanced firing and altered action potential properties in delay firing neurons following 9-AC treatment. (A-G)** Electrophysiological characterization of delay firing neurons from vehicle- and 9-AC-treated mice. **(A)** Representative traces showing action potential (AP) firing in response to increasing depolarizing current injections. **(B)** Frequency-current (F-I) relationship demonstrating the number of evoked APs in response to increasing current injections. Quantification of **(C)** resting membrane potential (RMP), **(D)** action potential threshold, **(E)** threshold distance (ΔV; VThreshold − Vrest), **(F)** rheobase, and **(G)** spike latency. **(H-N)** Electrophysiological characterization of tonic firing neurons, including **(H)** representative traces, **(I)** F-I relationship, and quantification of **(J)** RMP, **(K)** action potential threshold, **(L)** threshold distance (ΔV), **(M)** rheobase, and **(N)** spike latency. **(O-U)** Electrophysiological characterization of phasic firing neurons, including **(O)** representative traces, **(P)** F-I relationship, and quantification of **(Q)** RMP, **(R)** action potential threshold, **(S)** threshold distance (ΔV), **(T)** rheobase, and **(U)** spike latency. Frequency-current relationships (**B**, **I**, and **P**) were analyzed using two-way repeated-measures ANOVA. Electrophysiological parameters (**C-G**, **J-N**, and **Q-U**) were analyzed using unpaired two-tailed Student's t tests with Welch's correction. Data are presented as mean ± SEM (n = 3-9 neurons). Individual data points represent values from single neurons. Exact p values are indicated on the graphs. *$p < 0.05$, **$p < 0.01$.

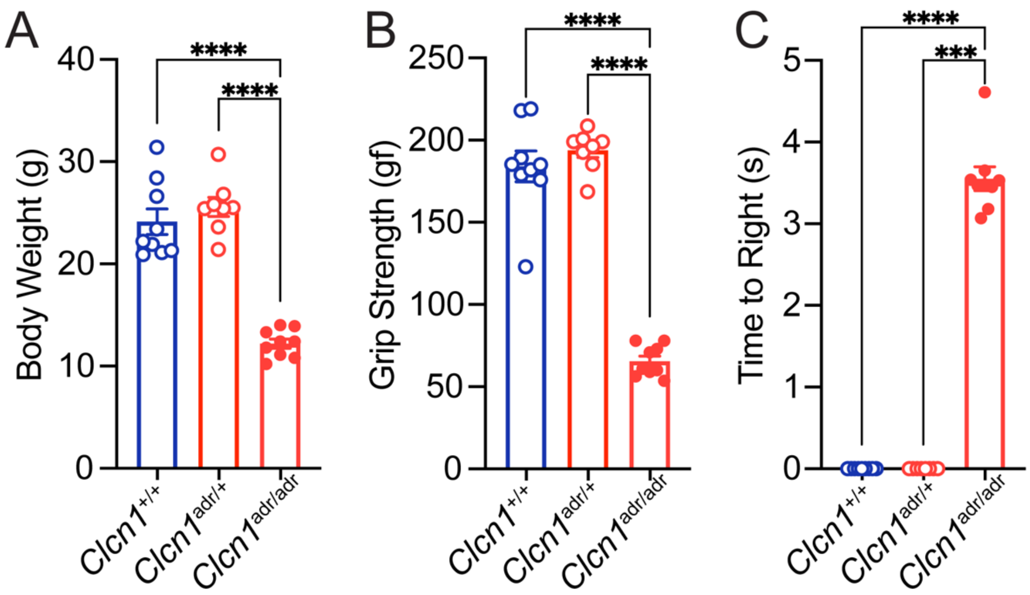


**Supplemental Figure 6. Homozygous *Clcn1*$^{adr/adr}$ mice exhibit reduced body weight, impaired muscle strength, and delayed righting reflex. (A)** Body weight of adult wildtype (*Clcn1*$^{+/+}$), heterozygous (*Clcn1*$^{adr/+}$), and homozygous myotonic (*Clcn1*$^{adr/adr}$) mice. **(B)** Forelimb grip strength measured using a digital grip strength meter. **(C)** Time to righting reflex (TRR) following placement in the supine position, a validated measure of myotonia based on delayed recovery from forced supine positioning. Homozygous *Clcn1*$^{adr/adr}$ mice exhibited reduced body weight and grip strength and displayed a marked delay in righting compared with both wildtype and heterozygous littermates, whereas heterozygous mice were indistinguishable from wildtype controls. Data in **(A)** and **(B)** were analyzed using one-way ANOVA followed by Tukey's multiple comparisons test. Data in **(C)** were analyzed using the Kruskal-Wallis test followed by Dunn's multiple comparisons test. Data are presented as mean ± SEM (n = 8-9 mice/group). Individual data points represent values from individual animals. ***$p < 0.001$, ****$p < 0.0001$.

| | |
|---|---|
| Current clamp: External solution for small and large DRG neurons (in mM) | 130 NaCl, 3 KCl, 2.5 $CaCl_2$, 0.6 $MgCl_2$, 10 HEPES, 10 Glucose (pH adjusted to 7.4) |
| Current Clamp: Internal solution for small DRG neurons (in mM) | 120 K-gluconate, 10 NaCl, 4 Mg-ATP, 10 HEPES, 5 EGTA, 2 $MgCl_2$ (pH adjusted to 7.2) |
| Current Clamp: Internal solution for large DRG neurons (in mM) | 110 K-methanesulfonate, 30 KCl, 5 NaCl, 1 $CaCl_2$, 2 $MgCl_2$, 2 Mg-ATP, 1 Li-GTP, 10 HEPES, 11 EGTA (pH adjusted to 7.2) |
| Voltage Clamp External solution for small DRG neurons (in mM) | 50 NaCl, 100 TEA-Cl, 1.8 $CaCl_2$, 0.1 $CdCl_2$, 1 $MgCl_2$, 10 D-glucose, and 10 HEPES (pH 7.3; 310 mOsm/L) |
| Voltage Clamp: Internal solution for small DRG neurons (in mM) | 140 CsF, 1.1 Cs-EGTA, 10 NaCl, and 15 HEPES (pH 7.3; 300 mOsm/L) |

**Supplemental Table 1:** Solutions for Electrophysiological Recordings

| Figure | Comparison | Statistical Test | P Value |
|---|---|---|---|
| **1B** | Time x Drug Interaction | Two-way RM ANOVA F (4, 72) = 30.13 | P<0.0001 |
| | Time | F (2.354, 42.37) = 30.13 | P<0.0001 |
| | Drug | F (1, 18) = 46.55 | P<0.0001 |
| | Time 30 min: Vehicle vs. 9-AC | Tukey's multiple comparisons test | **** P<0.0001 |
| | Time 60 min: Vehicle vs. 9-AC | Tukey's multiple comparisons test | *** P=0.0002 |
| | Time 120 min: Vehicle vs. 9-AC | Tukey's multiple comparisons test | ** P=0.0030 |
| | Time 180 min: Vehicle vs. 9-AC | Tukey's multiple comparisons test | * P=0.0337 |
| **1C** | Vehicle vs. 9-AC | Unpaired two-tailed t test with Welch's correction | ** P=0.0017 |
| **2B** | Time x Drug Interaction | Two-way RM ANOVA F (4, 56) = 20.88 | P<0.0001 |
| | Time | F (2.383, 33.36) = 29.79 | P<0.0001 |
| | Drug | F (1, 14) = 91.48 | P<0.0001 |
| | BL: Vehicle vs. 9-AC | Dunnett's multiple comparisons test | P=0.4456 |
| | Time 6 hours: Vehicle vs. 9-AC | Dunnett's multiple comparisons test | **** P<0.0001 |
| | Time 24 hours: Vehicle vs. 9-AC | Dunnett's multiple comparisons test | **** P<0.0001 |
| | Time 48 hours: Vehicle vs. 9-AC | Dunnett's multiple comparisons test | NS P=0.0567 |
| | Time 72 hours: Vehicle vs. 9-AC | Dunnett's multiple comparisons test | P=0.3658 |
| **2C** | Vehicle vs. 9-AC | Unpaired two-tailed t test | **** P<0.0001 |
| **2D** | Time x Drug Interaction | Two-way RM ANOVA F (4, 56) = 13.01 | P<0.0001 |
| | Time | F (2.924, 40.94) = 12.66 | P<0.0001 |
| | Drug | F (1, 14) = 13.61 | P<0.0001 |
| | BL: Vehicle vs. 9-AC | Dunnett's multiple comparisons test | P=0.1200 |
| | Time 6 hours: Vehicle vs. 9-AC | Dunnett's multiple comparisons test | **** P<0.0001 |
| | Time 24 hours: Vehicle vs. 9-AC | Dunnett's multiple comparisons test | ** P=0.0030 |
| | Time 48 hours: Vehicle vs. 9-AC | Dunnett's multiple comparisons test | P=0.1224 |
| | Time 72 hours: Vehicle vs. 9-AC | Dunnett's multiple comparisons test | P=0.7059 |
| **2E** | Vehicle vs. 9-AC | Unpaired two-tailed t test | *** P=0.0008 |
| **2F** | Time x Drug Interaction | Two-way RM ANOVA F (4, 56) = 76.67 | P<0.0001 |
| | Time | F (2.284, 31.98) = 70.86 | P<0.0001 |
| | Drug | F (1, 14) = 303.5 | P<0.0001 |
| | BL: Vehicle vs. 9-AC | Dunnett's multiple comparisons test | P=0.7941 |
| | Time 6 hours: Vehicle vs. 9-AC | Dunnett's multiple comparisons test | **** P<0.0001 |
| | Time 24 hours: Vehicle vs. 9-AC | Dunnett's multiple comparisons test | **** P<0.0001 |
| | Time 48 hours: Vehicle vs. 9-AC | Dunnett's multiple comparisons test | *** P=0.0002 |
| | Time 72 hours: Vehicle vs. 9-AC | Dunnett's multiple comparisons test | P=0.7790 |
| **2G** | Vehicle vs. 9-AC | Unpaired two-tailed t test | **** P<0.0001 |
| **2H** | Time x Drug Interaction | Two-way RM ANOVA F (4, 56) = 6.850 | P=0.0001 |
| | Time | F (2.284, 31.98) = 70.86 | P=0.0061 |
| | Drug | F (1, 14) = 303.5 | P=0.0018 |
| | BL: Vehicle vs. 9-AC | Dunnett's multiple comparisons test | P=0.7334 |
| | Time 6 hours: Vehicle vs. 9-AC | Dunnett's multiple comparisons test | **** P<0.0001 |
| | Time 24 hours: Vehicle vs. 9-AC | Dunnett's multiple comparisons test | * P=0.0229 |
| | Time 48 hours: Vehicle vs. 9-AC | Dunnett's multiple comparisons test | P=0.2712 |
| | Time 72 hours: Vehicle vs. 9-AC | Dunnett's multiple comparisons test | P=0.6651 |
| **2I** | Vehicle vs. 9-AC | Unpaired two-tailed t test | ** P=0.0015 |
| **3C** | Vehicle vs. 9-AC | Unpaired two-tailed t test with Welch's correction | P=0.7563 |
| **3D** | Vehicle vs. 9-AC | Unpaired two-tailed t test with Welch's correction | NS P=0.0637 |

| | | | |
|---|---|---|---|
| **3E** | Vehicle vs. 9-AC | Unpaired two-tailed t test with Welch's correction | *** P=0.0003 |
| **4C** | Vehicle vs. 9-AC | Unpaired two-tailed t test with Welch's correction | P=0.6924 |
| **4E** | Vehicle vs. 9-AC | Unpaired two-tailed t test with Welch's correction | * P=0.0327 |
| **4F** | Vehicle vs. 9-AC | Unpaired two-tailed t test with Welch's correction | * P=0.0287 |
| **4G** | Vehicle vs. 9-AC | Unpaired two-tailed t test with Welch's correction | NS P=0.0528 |
| **5E** | BL vs. 9-AC vs. Vehicle | One-way ANOVA<br>$F (2, 33) = 8.948$ | P=0.0008 |
| | BL vs. 9-AC | Tukey's multiple comparisons test | ** P=0.0029 |
| | BL vs. Vehicle | Tukey's multiple comparisons test | P=0.9902 |
| | 9-AC vs. Vehicle | Tukey's multiple comparisons test | ** P=0.0020 |
| **5I** | BL vs. 9-AC vs. Vehicle | One-way ANOVA<br>$F (2, 33) = 6.372$ | P=0.0046 |
| | BL vs. 9-AC | Tukey's multiple comparisons test | ** P=0.0096 |
| | BL vs. Vehicle | Tukey's multiple comparisons test | P=0.9939 |
| | 9-AC vs. Vehicle | Tukey's multiple comparisons test | * P=0.0125 |
| **5M** | BL vs. 9-AC vs. Vehicle | One-way ANOVA<br>$F (2, 33) = 9.031$ | P=0.0007 |
| | BL vs. 9-AC | Tukey's multiple comparisons test | ** P=0.0062 |
| | BL vs. Vehicle | Tukey's multiple comparisons test | P=0.7935 |
| | 9-AC vs. Vehicle | Tukey's multiple comparisons test | ** P=0.0011 |
| **5Q** | BL vs. 9-AC vs. Vehicle | One-way ANOVA<br>$F (2, 33) = 9.031$ | P<0.0001 |
| | BL vs. 9-AC | Tukey's multiple comparisons test | **** P<0.0001 |
| | BL vs. Vehicle | Tukey's multiple comparisons test | P=0.9773 |
| | 9-AC vs. Vehicle | Tukey's multiple comparisons test | **** P<0.0001 |
| **6D** | Vehicle vs. 9-AC | Unpaired two-tailed t test with Welch's correction | * P=0.0353 |
| **6G** | Vehicle vs. 9-AC | Unpaired two-tailed t test with Welch's correction | P=0.1659 |
| **6H** | Vehicle vs. 9-AC | Unpaired two-tailed t test with Welch's correction | P=0.3770 |
| **7B** | Vehicle vs. 9-AC | Unpaired two-tailed t test with Welch's correction | NS P=0.0828 |
| **7C** | Vehicle vs. 9-AC | Unpaired two-tailed t test with Welch's correction | NS P=0.0827 |
| **8C** | Myotonia x Drug<br>Interaction | Two-way ANOVA<br>$F (1, 35) = 0.06384$ | P=0.8020 |
| | Myotonia (Vehicle or 9-AC) | $F (1, 35) = 2.341$ | P=0.1350 |
| | Drug (Vehicle or Suzetrigine) | $F (1, 35) = 3.696$ | P=0.0627 |
| **8D** | Myotonia x Drug<br>Interaction | Two-way ANOVA<br>$F (1, 35) = 0.4668$ | P=0.4990 |
| | Myotonia (Vehicle or 9-AC) | $F (1, 35) = 4.896$ | P=0.0335 |
| | Drug (Vehicle or Suzetrigine) | $F (1, 35) = 0.2586$ | P=0.6143 |
| **8E** | Myotonia x Drug<br>Interaction | Two-way ANOVA<br>$F (1, 35) = 5.886$ | P=0.0206 |
| | Myotonia (Vehicle or 9-AC) | $F (1, 35) = 27.49$ | P<0.0001 |
| | Drug (Vehicle or Suzetrigine) | $F (1, 35) = 45.23$ | P<0.0001 |
| | Vehicle:Vehicle vs. Vehicle:Suzetrigine | Tukey's multiple comparisons test | * P=0.0158 |
| | Vehicle:Vehicle vs.9-AC:Vehicle | Tukey's multiple comparisons test | P<0.0001 |
| | Vehicle:Vehicle vs.9-AC:Suzetrigine | Tukey's multiple comparisons test | P=0.7103 |
| | Vehicle: Suzetrigine vs.9-AC:Vehicle | Tukey's multiple comparisons test | P<0.0001 |
| | Vehicle: Suzetrigine vs.9-AC:Suzetrigine | Tukey's multiple comparisons test | P=0.1800 |
| | 9-AC:Vehicle vs.9-AC:Suzetrigine | Tukey's multiple comparisons test | **** P<0.0001 |
| **9B** | Time post-Drug | Three-way RM ANOVA<br>$F (2.595, 75.25) = 5.581$ | P=0.0026 |

| | | | |
|---|---|---|---|
| | Vehicle (9-AC) vs. 9-AC | F (1, 29) = 81.12 | P<0.0001 |
| | Vehicle (Suzetrigine) vs. Suzetrigine | F (1, 29) = 12.05 | P=0.0016 |
| | Time post-drug x Vehicle (9-AC) vs. 9-AC | F (3, 87) = 7.881 | P=0.0001 |
| | Time post-drug x Vehicle (Suzetrigine) vs. Suzetrigine | F (3, 87) = 5.799 | P=0.0012 |
| | Vehicle (9-AC) vs. 9-AC vs Suzetrigine | F (1, 29) = 3.84 | P=0.0597 |
| | Time post-drug x Vehicle (9-AC) vs. Suzetrigine | F (3, 87) = 7.925 | P<0.0001 |
| | 0:Vehicle Vehicle vs. 0:Vehicle Suzetrigine | Tukey’s multiple comparisons test | P=0.9989 |
| | 0:Vehicle Vehicle vs. 0:9-AC Vehicle | Tukey’s multiple comparisons test | P=0.0002 |
| | 0:Vehicle Vehicle vs. 0:9-AC Suzetrigine | Tukey’s multiple comparisons test | |
| | 0:Vehicle Vehicle vs. 0:Vehicle Suzetrigine | Tukey’s multiple comparisons test | P=0.9989 |
| | 0:Vehicle Vehicle vs. 60:Vehicle Vehicle | Tukey’s multiple comparisons test | P>0.9999 |
| | 0:Vehicle Vehicle vs. 60:Vehicle Suzetrigine | Tukey’s multiple comparisons test | P>0.9999 |
| | 0:Vehicle Vehicle vs. 60:9-AC Vehicle | Tukey’s multiple comparisons test | P=0.0002 |
| | 0:Vehicle Vehicle vs. 60:9-AC Suzetrigine | Tukey’s multiple comparisons test | P=0.8548 |
| | 0:Vehicle Vehicle vs. 120:Vehicle Vehicle | Tukey’s multiple comparisons test | P>0.9999 |
| | 0:Vehicle Vehicle vs. 120:Vehicle Suzetrigine | Tukey’s multiple comparisons test | P>0.9999 |
| | 0:Vehicle Vehicle vs. 120:9-AC Vehicle | Tukey’s multiple comparisons test | P=0.0002 |
| | 0:Vehicle Vehicle vs. 120:9-AC Suzetrigine | Tukey’s multiple comparisons test | P>0.9999 |
| | 0:Vehicle Vehicle vs. 180:Vehicle Vehicle | Tukey’s multiple comparisons test | P>0.9999 |
| | 0:Vehicle Vehicle vs. 180:Vehicle Suzetrigine | Tukey’s multiple comparisons test | P=0.9980 |
| | 0:Vehicle Vehicle vs. 180:9-AC Vehicle | Tukey’s multiple comparisons test | P=0.0002 |
| | 0:Vehicle Vehicle vs. 180:9-AC Suzetrigine | Tukey’s multiple comparisons test | P=0.0069 |
| | 0:Vehicle Suzetrigine vs. 0:9-AC Vehicle | Tukey’s multiple comparisons test | P=0.0003 |
| | 0:Vehicle Suzetrigine vs. 0:9-AC Suzetrigine | Tukey’s multiple comparisons test | P=0.0003 |
| | 0:Vehicle Suzetrigine vs. 60:Vehicle Vehicle | Tukey’s multiple comparisons test | P>0.9999 |
| | 0:Vehicle Suzetrigine vs. 60:Vehicle Suzetrigine | Tukey’s multiple comparisons test | P=0.7206 |
| | 0:Vehicle Suzetrigine vs. 60:9-AC Vehicle | Tukey’s multiple comparisons test | P=0.0002 |
| | 0:Vehicle Suzetrigine vs. 60:9-AC Suzetrigine | Tukey’s multiple comparisons test | P=0.5836 |
| | 0:Vehicle Suzetrigine vs. 120:Vehicle Vehicle | Tukey’s multiple comparisons test | P=0.9924 |
| | 0:Vehicle Suzetrigine vs. 120:Vehicle Suzetrigine | Tukey’s multiple comparisons test | P>0.9999 |
| | 0:Vehicle Suzetrigine vs. 120:9-AC Vehicle | Tukey’s multiple comparisons test | P=0.0003 |
| | 0:Vehicle Suzetrigine vs. 120:9-AC Suzetrigine | Tukey’s multiple comparisons test | P>0.9999 |
| | 0:Vehicle Suzetrigine vs. 180:Vehicle Vehicle | Tukey’s multiple comparisons test | P=0.9691 |
| | 0:Vehicle Suzetrigine vs. 180:Vehicle Suzetrigine | Tukey’s multiple comparisons test | P>0.9999 |
| | 0:Vehicle Suzetrigine vs. 180:9-AC Vehicle | Tukey’s multiple comparisons test | P=0.0002 |
| | 0:Vehicle Suzetrigine vs. 180:9-AC Suzetrigine | Tukey’s multiple comparisons test | P=0.0038 |
| | 0:9-AC Vehicle vs. 0:9-AC Suzetrigine | Tukey’s multiple comparisons test | P>0.9999 |
| | 0:9-AC Vehicle vs. 60:Vehicle Vehicle | Tukey’s multiple comparisons test | P=0.0025 |
| | 0:9-AC Vehicle vs. 60:Vehicle Suzetrigine | Tukey’s multiple comparisons test | P=0.0010 |
| | 0:9-AC Vehicle vs. 60:9-AC Vehicle | Tukey’s multiple comparisons test | P=0.9998 |
| | 0:9-AC Vehicle vs. 60:9-AC Suzetrigine | Tukey’s multiple comparisons test | P=0.1629 |
| | 0:9-AC Vehicle vs. 120:Vehicle Vehicle | Tukey’s multiple comparisons test | P=0.0014 |
| | 0:9-AC Vehicle vs. 120:Vehicle Suzetrigine | Tukey’s multiple comparisons test | P=0.0081 |
| | 0:9-AC Vehicle vs. 120:9-AC Vehicle | Tukey’s multiple comparisons test | P>0.9999 |
| | 0:9-AC Vehicle vs. 120:9-AC Suzetrigine | Tukey’s multiple comparisons test | P=0.0003 |
| | 0:9-AC Vehicle vs. 180:Vehicle Vehicle | Tukey’s multiple comparisons test | P=0.0006 |
| | 0:9-AC Vehicle vs. 180:Vehicle Suzetrigine | Tukey’s multiple comparisons test | P=0.0016 |
| | 0:9-AC Vehicle vs. 180:9-AC Vehicle | Tukey’s multiple comparisons test | P>0.9999 |
| | 0:9-AC Vehicle vs. 180:9-AC Suzetrigine | Tukey’s multiple comparisons test | P=0.9990 |
| | 0:9-AC Suzetrigine vs. 60:Vehicle Vehicle | Tukey’s multiple comparisons test | P=0.0020 |
| | 0:9-AC Suzetrigine vs. 60:Vehicle Suzetrigine | Tukey’s multiple comparisons test | P=0.0010 |
| | 0:9-AC Suzetrigine vs. 60:9-AC Vehicle | Tukey’s multiple comparisons test | P>0.9999 |
| | 0:9-AC Suzetrigine vs. 60:9-AC Suzetrigine | Tukey’s multiple comparisons test | * P=0.0416 |

| | | | |
|---|---|---|---|
| | 0:9-AC Suzetrigine vs. 120:Vehicle Vehicle | Tukey's multiple comparisons test | P=0.0014 |
| | 0:9-AC Suzetrigine vs. 120:Vehicle Suzetrigine | Tukey's multiple comparisons test | P=0.0059 |
| | 0:9-AC Suzetrigine vs. 120:9-AC Vehicle | Tukey's multiple comparisons test | P>0.9999 |
| | 0:9-AC Suzetrigine vs. 120:9-AC Suzetrigine | Tukey's multiple comparisons test | ** P=0.0016 |
| | 0:9-AC Suzetrigine vs. 180:Vehicle Vehicle | Tukey's multiple comparisons test | P=0.0009 |
| | 0:9-AC Suzetrigine vs. 180:Vehicle Suzetrigine | Tukey's multiple comparisons test | P=0.0012 |
| | 0:9-AC Suzetrigine vs. 180:9-AC Vehicle | Tukey's multiple comparisons test | P>0.9999 |
| | 0:9-AC Suzetrigine vs. 180:9-AC Suzetrigine | Tukey's multiple comparisons test | P=0.9584 |
| | 60:Vehicle Vehicle vs. 60:Vehicle Suzetrigine | Tukey's multiple comparisons test | P>0.9999 |
| | 60:Vehicle Vehicle vs. 60:9-AC Vehicle | Tukey's multiple comparisons test | P=0.0016 |
| | 60:Vehicle Vehicle vs. 60:9-AC Suzetrigine | Tukey's multiple comparisons test | P=0.8890 |
| | 60:Vehicle Vehicle vs. 120:Vehicle Vehicle | Tukey's multiple comparisons test | P>0.9999 |
| | 60:Vehicle Vehicle vs. 120:Vehicle Suzetrigine | Tukey's multiple comparisons test | P>0.9999 |
| | 60:Vehicle Vehicle vs. 120:9-AC Vehicle | Tukey's multiple comparisons test | P=0.0025 |
| | 60:Vehicle Vehicle vs. 120:9-AC Suzetrigine | Tukey's multiple comparisons test | P>0.9999 |
| | 60:Vehicle Vehicle vs. 180:Vehicle Vehicle | Tukey's multiple comparisons test | P=0.9999 |
| | 60:Vehicle Vehicle vs. 180:Vehicle Suzetrigine | Tukey's multiple comparisons test | P>0.9999 |
| | 60:Vehicle Vehicle vs. 180:9-AC Vehicle | Tukey's multiple comparisons test | P=0.0017 |
| | 60:Vehicle Vehicle vs. 180:9-AC Suzetrigine | Tukey's multiple comparisons test | P=0.0192 |
| | 60:Vehicle Suzetrigine vs. 60:9-AC Vehicle | Tukey's multiple comparisons test | P=0.0007 |
| | 60:Vehicle Suzetrigine vs. 60:9-AC Suzetrigine | Tukey's multiple comparisons test | P=0.9219 |
| | 60:Vehicle Suzetrigine vs. 120:Vehicle Vehicle | Tukey's multiple comparisons test | P>0.9999 |
| | 60:Vehicle Suzetrigine vs. 120:Vehicle Suzetrigine | Tukey's multiple comparisons test | P>0.9999 |
| | 60:Vehicle Suzetrigine vs. 120:9-AC Vehicle | Tukey's multiple comparisons test | P=0.0011 |
| | 60:Vehicle Suzetrigine vs. 120:9-AC Suzetrigine | Tukey's multiple comparisons test | P>0.9999 |
| | 60:Vehicle Suzetrigine vs. 180:Vehicle Vehicle | Tukey's multiple comparisons test | P>0.9999 |
| | 60:Vehicle Suzetrigine vs. 180:Vehicle Suzetrigine | Tukey's multiple comparisons test | P=0.9686 |
| | 60:Vehicle Suzetrigine vs. 180:9-AC Vehicle | Tukey's multiple comparisons test | P=0.0007 |
| | 60:Vehicle Suzetrigine vs. 180:9-AC Suzetrigine | Tukey's multiple comparisons test | P=0.0147 |
| | 60:9-AC Vehicle vs. 60:9-AC Suzetrigine | Tukey's multiple comparisons test | P=0.1098 |
| | 60:9-AC Vehicle vs. 120:Vehicle Vehicle | Tukey's multiple comparisons test | P=0.0009 |
| | 60:9-AC Vehicle vs. 120:Vehicle Suzetrigine | Tukey's multiple comparisons test | P=0.0054 |
| | 60:9-AC Vehicle vs. 120:9-AC Vehicle | Tukey's multiple comparisons test | P>0.9999 |
| | 60:9-AC Vehicle vs. 120:9-AC Suzetrigine | Tukey's multiple comparisons test | P=0.0002 |
| | 60:9-AC Vehicle vs. 180:Vehicle Vehicle | Tukey's multiple comparisons test | P=0.0005 |
| | 60:9-AC Vehicle vs. 180:Vehicle Suzetrigine | Tukey's multiple comparisons test | P=0.0010 |
| | 60:9-AC Vehicle vs. 180:9-AC Vehicle | Tukey's multiple comparisons test | P>0.9999 |
| | 60:9-AC Vehicle vs. 180:9-AC Suzetrigine | Tukey's multiple comparisons test | P=0.9846 |
| | 60:9-AC Suzetrigine vs. 120:Vehicle Vehicle | Tukey's multiple comparisons test | P=0.9740 |
| | 60:9-AC Suzetrigine vs. 120:Vehicle Suzetrigine | Tukey's multiple comparisons test | P=0.8648 |
| | 60:9-AC Suzetrigine vs. 120:9-AC Vehicle | Tukey's multiple comparisons test | P=0.1630 |
| | 60:9-AC Suzetrigine vs. 120:9-AC Suzetrigine | Tukey's multiple comparisons test | P=0.7027 |
| | 60:9-AC Suzetrigine vs. 180:Vehicle Vehicle | Tukey's multiple comparisons test | P=0.9754 |
| | 60:9-AC Suzetrigine vs. 180:Vehicle Suzetrigine | Tukey's multiple comparisons test | P=0.6179 |
| | 60:9-AC Suzetrigine vs. 180:9-AC Vehicle | Tukey's multiple comparisons test | P=0.1118 |
| | 60:9-AC Suzetrigine vs. 180:9-AC Suzetrigine | Tukey's multiple comparisons test | P=0.7074 |
| | 120:Vehicle Vehicle vs. 120:Vehicle Suzetrigine | Tukey's multiple comparisons test | P>0.9999 |
| | 120:Vehicle Vehicle vs. 120:9-AC Vehicle | Tukey's multiple comparisons test | P=0.0014 |
| | 120:Vehicle Vehicle vs. 120:9-AC Suzetrigine | Tukey's multiple comparisons test | P=0.9987 |
| | 120:Vehicle Vehicle vs. 180:Vehicle Vehicle | Tukey's multiple comparisons test | P>0.9999 |

| | | | |
|---|---|---|---|
| | 120:Vehicle Vehicle vs. 180:Vehicle Suzetrigine | Tukey's multiple comparisons test | P=0.9903 |
| | 120:Vehicle Vehicle vs. 180:9-AC Vehicle | Tukey's multiple comparisons test | P=0.0010 |
| | 120:Vehicle Vehicle vs. 180:9-AC Suzetrigine | Tukey's multiple comparisons test | P=0.0214 |
| | 120:Vehicle Suzetrigine vs. 120:9-AC Vehicle | Tukey's multiple comparisons test | P=0.0081 |
| | 120:Vehicle Suzetrigine vs. 120:9-AC Suzetrigine | Tukey's multiple comparisons test | P>0.9999 |
| | 120:Vehicle Suzetrigine vs. 180:Vehicle Vehicle | Tukey's multiple comparisons test | P=0.9995 |
| | 120:Vehicle Suzetrigine vs. 180:Vehicle Suzetrigine | Tukey's multiple comparisons test | P>0.9999 |
| | 120:Vehicle Suzetrigine vs. 180:9-AC Vehicle | Tukey's multiple comparisons test | P=0.0054 |
| | 120:Vehicle Suzetrigine vs. 180:9-AC Suzetrigine | Tukey's multiple comparisons test | P=0.0365 |
| | 120:9-AC Vehicle vs. 120:9-AC Suzetrigine | Tukey's multiple comparisons test | P=0.0003 |
| | 120:9-AC Vehicle vs. 180:Vehicle Vehicle | Tukey's multiple comparisons test | P=0.0007 |
| | 120:9-AC Vehicle vs. 180:Vehicle Suzetrigine | Tukey's multiple comparisons test | P=0.0016 |
| | 120:9-AC Vehicle vs. 180:9-AC Vehicle | Tukey's multiple comparisons test | P=0.9999 |
| | 120:9-AC Vehicle vs. 180:9-AC Suzetrigine | Tukey's multiple comparisons test | P=0.9990 |
| | 120:9-AC Suzetrigine vs. 180:Vehicle Vehicle | Tukey's multiple comparisons test | P=0.9908 |
| | 120:9-AC Suzetrigine vs. 180:Vehicle Suzetrigine | Tukey's multiple comparisons test | P>0.9999 |
| | 120:9-AC Suzetrigine vs. 180:9-AC Vehicle | Tukey's multiple comparisons test | P=0.0002 |
| | 120:9-AC Suzetrigine vs. 180:9-AC Suzetrigine | Tukey's multiple comparisons test | P=0.0033 |
| | 180:Vehicle Vehicle vs. 180:Vehicle Suzetrigine | Tukey's multiple comparisons test | P=0.9698 |
| | 180:Vehicle Vehicle vs. 180:9-AC Vehicle | Tukey's multiple comparisons test | P=0.0005 |
| | 180:Vehicle Vehicle vs. 180:9-AC Suzetrigine | Tukey's multiple comparisons test | P=0.0166 |
| | 180:Vehicle Suzetrigine vs. 180:9-AC Vehicle | Tukey's multiple comparisons test | P=0.0011 |
| | 180:Vehicle Suzetrigine vs. 180:9-AC Suzetrigine | Tukey's multiple comparisons test | P=0.0094 |
| | 180:9-AC Vehicle vs. 180:9-AC Suzetrigine | Tukey's multiple comparisons test | P=0.9856 |
| **9C** | Myotonia x Drug Interaction | RM Mixed-effects model F (1, 14) = 7.876 | P=0.0140 |
| | Myotonia (Vehicle or 9-AC) | F (1, 14) = 63.72 | P<0.0001 |
| | Drug (Vehicle or Suzetrigine) | F (1, 15) = 15.22 | P=0.0014 |
| | Vehicle: Vehicle vs. Suzetrigine | Uncorrected Fisher's LSD | P=0.3969 |
| | 9-AC: Vehicle vs. Suzetrigine | Uncorrected Fisher's LSD | **** P<0.0001 |
| | Vehicle: Vehicle vs. 9-AC | Uncorrected Fisher's LSD | P<0.0001 |
| | Suzetrigine: Vehicle vs. 9-AC | Uncorrected Fisher's LSD | P=0.0023 |
| **9D** | Time post-Drug | Three-way RM ANOVA F (2.759, 80.01) = 3.723 | P=0.0171 |
| | Vehicle (9-AC) vs. 9-AC | F (1, 29) = 67.24 | P<0.0001 |
| | Vehicle (Suzetrigine) vs. Suzetrigine | F (1, 29) = 4.402 | P=0.0447 |
| | Time post-drug x Vehicle (9-AC) vs. 9-AC | F (3, 87) = 4.775 | P=0.0040 |
| | Time post-drug x Vehicle (Suzetrigine) vs. Suzetrigine | F (3, 87) = 6.580 | P=0.0005 |
| | Vehicle (9-AC) vs. 9-AC vs Suzetrigine | F (1, 29) = 2.873 | P=0.1008 |
| | Time post-drug x Vehicle (9-AC) vs. Suzetrigine | F (3, 87) = 9.576 | P<0.0001 |
| | 0:Vehicle Vehicle vs. 0:Vehicle Suzetrigine | Tukey's multiple comparisons test | P>0.9999 |
| | 0:Vehicle Vehicle vs. 0:9-AC Vehicle | Tukey's multiple comparisons test | P=0.0179 |
| | 0:Vehicle Vehicle vs. 0:9-AC Suzetrigine | Tukey's multiple comparisons test | P=0.0033 |
| | 0:Vehicle Vehicle vs. 60:Vehicle Vehicle | Tukey's multiple comparisons test | P>0.9999 |
| | 0:Vehicle Vehicle vs. 60:Vehicle Suzetrigine | Tukey's multiple comparisons test | P>0.9999 |
| | 0:Vehicle Vehicle vs. 60:9-AC Vehicle | Tukey's multiple comparisons test | P=0.0008 |
| | 0:Vehicle Vehicle vs. 60:9-AC Suzetrigine | Tukey's multiple comparisons test | P=0.9093 |
| | 0:Vehicle Vehicle vs. 120:Vehicle Vehicle | Tukey's multiple comparisons test | P>0.9999 |
| | 0:Vehicle Vehicle vs. 120:Vehicle Suzetrigine | Tukey's multiple comparisons test | P>0.9999 |

| | | | |
|---|---|---|---|
| | 0:Vehicle Vehicle vs. 120:9-AC Vehicle | Tukey’s multiple comparisons test | P=0.0050 |
| | 0:Vehicle Vehicle vs. 120:9-AC Suzetrigine | Tukey’s multiple comparisons test | P=0.9956 |
| | 0:Vehicle Vehicle vs. 180:Vehicle Vehicle | Tukey’s multiple comparisons test | P=0.9979 |
| | 0:Vehicle Vehicle vs. 180:Vehicle Suzetrigine | Tukey’s multiple comparisons test | P>0.9999 |
| | 0:Vehicle Vehicle vs. 180:9-AC Vehicle | Tukey’s multiple comparisons test | P=0.0365 |
| | 0:Vehicle Vehicle vs. 180:9-AC Suzetrigine | Tukey’s multiple comparisons test | P=0.1824 |
| | 0:Vehicle Suzetrigine vs. 0:9-AC Vehicle | Tukey’s multiple comparisons test | P=0.0133 |
| | 0:Vehicle Suzetrigine vs. 0:9-AC Suzetrigine | Tukey’s multiple comparisons test | P=0.0026 |
| | 0:Vehicle Suzetrigine vs. 60:Vehicle Vehicle | Tukey’s multiple comparisons test | P>0.9999 |
| | 0:Vehicle Suzetrigine vs. 60:Vehicle Suzetrigine | Tukey’s multiple comparisons test | P>0.9999 |
| | 0:Vehicle Suzetrigine vs. 60:9-AC Vehicle | Tukey’s multiple comparisons test | P=0.0005 |
| | 0:Vehicle Suzetrigine vs. 60:9-AC Suzetrigine | Tukey’s multiple comparisons test | P=0.7993 |
| | 0:Vehicle Suzetrigine vs. 120:Vehicle Vehicle | Tukey’s multiple comparisons test | P>0.9999 |
| | 0:Vehicle Suzetrigine vs. 120:Vehicle Suzetrigine | Tukey’s multiple comparisons test | P=0.9982 |
| | 0:Vehicle Suzetrigine vs. 120:9-AC Vehicle | Tukey’s multiple comparisons test | P=0.0037 |
| | 0:Vehicle Suzetrigine vs. 120:9-AC Suzetrigine | Tukey’s multiple comparisons test | P=0.9787 |
| | 0:Vehicle Suzetrigine vs. 180:Vehicle Vehicle | Tukey’s multiple comparisons test | P=0.9972 |
| | 0:Vehicle Suzetrigine vs. 180:Vehicle Suzetrigine | Tukey’s multiple comparisons test | P>0.9999 |
| | 0:Vehicle Suzetrigine vs. 180:9-AC Vehicle | Tukey’s multiple comparisons test | P=0.0263 |
| | 0:Vehicle Suzetrigine vs. 180:9-AC Suzetrigine | Tukey’s multiple comparisons test | P=0.1456 |
| | 0:9-AC Vehicle vs. 0:9-AC Suzetrigine | Tukey’s multiple comparisons test | P=0.9884 |
| | 0:9-AC Vehicle vs. 60:Vehicle Vehicle | Tukey’s multiple comparisons test | P=0.0734 |
| | 0:9-AC Vehicle vs. 60:Vehicle Suzetrigine | Tukey’s multiple comparisons test | P=0.0173 |
| | 0:9-AC Vehicle vs. 60:9-AC Vehicle | Tukey’s multiple comparisons test | P=0.9425 |
| | 0:9-AC Vehicle vs. 60:9-AC Suzetrigine | Tukey’s multiple comparisons test | P=0.1518 |
| | 0:9-AC Vehicle vs. 120:Vehicle Vehicle | Tukey’s multiple comparisons test | P=0.0113 |
| | 0:9-AC Vehicle vs. 120:Vehicle Suzetrigine | Tukey’s multiple comparisons test | P=0.0296 |
| | 0:9-AC Vehicle vs. 120:9-AC Vehicle | Tukey’s multiple comparisons test | P=0.9979 |
| | 0:9-AC Vehicle vs. 120:9-AC Suzetrigine | Tukey’s multiple comparisons test | P=0.1249 |
| | 0:9-AC Vehicle vs. 180:Vehicle Vehicle | Tukey’s multiple comparisons test | P=0.0582 |
| | 0:9-AC Vehicle vs. 180:Vehicle Suzetrigine | Tukey’s multiple comparisons test | P=0.0133 |
| | 0:9-AC Vehicle vs. 180:9-AC Vehicle | Tukey’s multiple comparisons test | P=0.9425 |
| | 0:9-AC Vehicle vs. 180:9-AC Suzetrigine | Tukey’s multiple comparisons test | P>0.9999 |
| | 0:9-AC Suzetrigine vs. 60:Vehicle Vehicle | Tukey’s multiple comparisons test | P=0.0100 |
| | 0:9-AC Suzetrigine vs. 60:Vehicle Suzetrigine | Tukey’s multiple comparisons test | P=0.0026 |
| | 0:9-AC Suzetrigine vs. 60:9-AC Vehicle | Tukey’s multiple comparisons test | P=0.9998 |
| | 0:9-AC Suzetrigine vs. 60:9-AC Suzetrigine | Tukey’s multiple comparisons test | NS P=0.0561 |
| | 0:9-AC Suzetrigine vs. 120:Vehicle Vehicle | Tukey’s multiple comparisons test | P=0.0022 |
| | 0:9-AC Suzetrigine vs. 120:Vehicle Suzetrigine | Tukey’s multiple comparisons test | P=0.0058 |
| | 0:9-AC Suzetrigine vs. 120:9-AC Vehicle | Tukey’s multiple comparisons test | P>0.9999 |
| | 0:9-AC Suzetrigine vs. 120:9-AC Suzetrigine | Tukey’s multiple comparisons test | NS 0.0766 |
| | 0:9-AC Suzetrigine vs. 180:Vehicle Vehicle | Tukey’s multiple comparisons test | P=0.0082 |
| | 0:9-AC Suzetrigine vs. 180:Vehicle Suzetrigine | Tukey’s multiple comparisons test | P=0.0026 |
| | 0:9-AC Suzetrigine vs. 180:9-AC Vehicle | Tukey’s multiple comparisons test | P=0.8564 |
| | 0:9-AC Suzetrigine vs. 180:9-AC Suzetrigine | Tukey’s multiple comparisons test | P=0.9031 |
| | 60:Vehicle Vehicle vs. 60:Vehicle Suzetrigine | Tukey’s multiple comparisons test | P>0.9999 |
| | 60:Vehicle Vehicle vs. 60:9-AC Vehicle | Tukey’s multiple comparisons test | P=0.0121 |
| | 60:Vehicle Vehicle vs. 60:9-AC Suzetrigine | Tukey’s multiple comparisons test | P=0.9993 |
| | 60:Vehicle Vehicle vs. 120:Vehicle Vehicle | Tukey’s multiple comparisons test | P=0.9979 |
| | 60:Vehicle Vehicle vs. 120:Vehicle Suzetrigine | Tukey’s multiple comparisons test | P>0.9999 |
| | 60:Vehicle Vehicle vs. 120:9-AC Vehicle | Tukey’s multiple comparisons test | P=0.0245 |
| | 60:Vehicle Vehicle vs. 120:9-AC Suzetrigine | Tukey’s multiple comparisons test | P>0.9999 |
| | 60:Vehicle Vehicle vs. 180:Vehicle Vehicle | Tukey’s multiple comparisons test | P>0.9999 |

| | | | |
|---|---|---|---|
| | 60:Vehicle Vehicle vs. 180:Vehicle Suzetrigine | Tukey's multiple comparisons test | P>0.9999 |
| | 60:Vehicle Vehicle vs. 180:9-AC Vehicle | Tukey's multiple comparisons test | P=0.1575 |
| | 60:Vehicle Vehicle vs. 180:9-AC Suzetrigine | Tukey's multiple comparisons test | P=0.4068 |
| | 60:Vehicle Suzetrigine vs. 60:9-AC Vehicle | Tukey's multiple comparisons test | P=0.0010 |
| | 60:Vehicle Suzetrigine vs. 60:9-AC Suzetrigine | Tukey's multiple comparisons test | P=0.9121 |
| | 60:Vehicle Suzetrigine vs. 120:Vehicle Vehicle | Tukey's multiple comparisons test | P>0.9999 |
| | 60:Vehicle Suzetrigine vs. 120:Vehicle Suzetrigine | Tukey's multiple comparisons test | P>0.9999 |
| | 60:Vehicle Suzetrigine vs. 120:9-AC Vehicle | Tukey's multiple comparisons test | P=0.0046 |
| | 60:Vehicle Suzetrigine vs. 120:9-AC Suzetrigine | Tukey's multiple comparisons test | P=0.9932 |
| | 60:Vehicle Suzetrigine vs. 180:Vehicle Vehicle | Tukey's multiple comparisons test | P=0.9994 |
| | 60:Vehicle Suzetrigine vs. 180:Vehicle Suzetrigine | Tukey's multiple comparisons test | P>0.9999 |
| | 60:Vehicle Suzetrigine vs. 180:9-AC Vehicle | Tukey's multiple comparisons test | P=0.0379 |
| | 60:Vehicle Suzetrigine vs. 180:9-AC Suzetrigine | Tukey's multiple comparisons test | P=0.1778 |
| | 60:9-AC Vehicle vs. 60:9-AC Suzetrigine | Tukey's multiple comparisons test | P=0.0149 |
| | 60:9-AC Vehicle vs. 120:Vehicle Vehicle | Tukey's multiple comparisons test | P=0.0004 |
| | 60:9-AC Vehicle vs. 120:Vehicle Suzetrigine | Tukey's multiple comparisons test | P=0.0013 |
| | 60:9-AC Vehicle vs. 120:9-AC Vehicle | Tukey's multiple comparisons test | P>0.9999 |
| | 60:9-AC Vehicle vs. 120:9-AC Suzetrigine | Tukey's multiple comparisons test | P=0.0156 |
| | 60:9-AC Vehicle vs. 180:Vehicle Vehicle | Tukey's multiple comparisons test | P=0.0043 |
| | 60:9-AC Vehicle vs. 180:Vehicle Suzetrigine | Tukey's multiple comparisons test | P=0.0005 |
| | 60:9-AC Vehicle vs. 180:9-AC Vehicle | Tukey's multiple comparisons test | P=0.4787 |
| | 60:9-AC Vehicle vs. 180:9-AC Suzetrigine | Tukey's multiple comparisons test | P=0.9846 |
| | 60:9-AC Suzetrigine vs. 120:Vehicle Vehicle | Tukey's multiple comparisons test | P=0.7312 |
| | 60:9-AC Suzetrigine vs. 120:Vehicle Suzetrigine | Tukey's multiple comparisons test | P=0.9887 |
| | 60:9-AC Suzetrigine vs. 120:9-AC Vehicle | Tukey's multiple comparisons test | P=0.0468 |
| | 60:9-AC Suzetrigine vs. 120:9-AC Suzetrigine | Tukey's multiple comparisons test | P>0.9999 |
| | 60:9-AC Suzetrigine vs. 180:Vehicle Vehicle | Tukey's multiple comparisons test | P>0.9999 |
| | 60:9-AC Suzetrigine vs. 180:Vehicle Suzetrigine | Tukey's multiple comparisons test | P=0.7993 |
| | 60:9-AC Suzetrigine vs. 180:9-AC Vehicle | Tukey's multiple comparisons test | P=0.3212 |
| | 60:9-AC Suzetrigine vs. 180:9-AC Suzetrigine | Tukey's multiple comparisons test | P=0.3242 |
| | 120:Vehicle Vehicle vs. 120:Vehicle Suzetrigine | Tukey's multiple comparisons test | P=0.9959 |
| | 120:Vehicle Vehicle vs. 120:9-AC Vehicle | Tukey's multiple comparisons test | P=0.0031 |
| | 120:Vehicle Vehicle vs. 120:9-AC Suzetrigine | Tukey's multiple comparisons test | P=0.9572 |
| | 120:Vehicle Vehicle vs. 180:Vehicle Vehicle | Tukey's multiple comparisons test | P=0.9922 |
| | 120:Vehicle Vehicle vs. 180:Vehicle Suzetrigine | Tukey's multiple comparisons test | P>0.9999 |
| | 120:Vehicle Vehicle vs. 180:9-AC Vehicle | Tukey's multiple comparisons test | P=0.0225 |
| | 120:Vehicle Vehicle vs. 180:9-AC Suzetrigine | Tukey's multiple comparisons test | P=0.1276 |
| | 120:Vehicle Suzetrigine vs. 120:9-AC Vehicle | Tukey's multiple comparisons test | P=0.0087 |
| | 120:Vehicle Suzetrigine vs. 120:9-AC Suzetrigine | Tukey's multiple comparisons test | P>0.9999 |
| | 120:Vehicle Suzetrigine vs. 180:Vehicle Vehicle | Tukey's multiple comparisons test | P>0.9999 |
| | 120:Vehicle Suzetrigine vs. 180:Vehicle Suzetrigine | Tukey's multiple comparisons test | P=0.9982 |
| | 120:Vehicle Suzetrigine vs. 180:9-AC Vehicle | Tukey's multiple comparisons test | P=0.0594 |
| | 120:Vehicle Suzetrigine vs. 180:9-AC Suzetrigine | Tukey's multiple comparisons test | P=0.2603 |
| | 120:9-AC Vehicle vs. 120:9-AC Suzetrigine | Tukey's multiple comparisons test | P=0.0396 |
| | 120:9-AC Vehicle vs. 180:Vehicle Vehicle | Tukey's multiple comparisons test | P=0.0166 |
| | 120:9-AC Vehicle vs. 180:Vehicle Suzetrigine | Tukey's multiple comparisons test | P=0.0037 |
| | 120:9-AC Vehicle vs. 180:9-AC Vehicle | Tukey's multiple comparisons test | P=0.8322 |

| | | | |
|---|---|---|---|
| | 120:9-AC Vehicle vs. 180:9-AC Suzetrigine | Tukey's multiple comparisons test | P=0.9926 |
| | 120:9-AC Suzetrigine vs. 180:Vehicle Vehicle | Tukey's multiple comparisons test | P>0.9999 |
| | 120:9-AC Suzetrigine vs. 180:Vehicle Suzetrigine | Tukey's multiple comparisons test | P=0.9787 |
| | 120:9-AC Suzetrigine vs. 180:9-AC Vehicle | Tukey's multiple comparisons test | P=0.2662 |
| | 120:9-AC Suzetrigine vs. 180:9-AC Suzetrigine | Tukey's multiple comparisons test | P=0.1473 |
| | 180:Vehicle Vehicle vs. 180:Vehicle Suzetrigine | Tukey's multiple comparisons test | P=0.9972 |
| | 180:Vehicle Vehicle vs. 180:9-AC Vehicle | Tukey's multiple comparisons test | P=0.1285 |
| | 180:Vehicle Vehicle vs. 180:9-AC Suzetrigine | Tukey's multiple comparisons test | P=0.4108 |
| | 180:Vehicle Suzetrigine vs. 180:9-AC Vehicle | Tukey's multiple comparisons test | P=0.0263 |
| | 180:Vehicle Suzetrigine vs. 180:9-AC Suzetrigine | Tukey's multiple comparisons test | P=0.1456 |
| | 180:9-AC Vehicle vs. 180:9-AC Suzetrigine | Tukey's multiple comparisons test | P>0.9999 |
| **9E** | Myotonia x Drug Interaction | RM Mixed-effects model F (1, 29) = 58.93 | P<0.0001 |
| | Myotonia (Vehicle or 9-AC) | F (1, 29) = 8.086 | P=0.0081 |
| | Drug (Vehicle or Suzetrigine) | F (1, 29) = 7.014 | P=0.0129 |
| | Vehicle: Vehicle vs. Suzetrigine | Uncorrected Fisher's LSD | P=0.8896 |
| | 9-AC: Vehicle vs. Suzetrigine | Uncorrected Fisher's LSD | *** P=0.0006 |
| | Vehicle: Vehicle vs. 9-AC | Uncorrected Fisher's LSD | P<0.0001 |
| | Suzetrigine: Vehicle vs. 9-AC | Uncorrected Fisher's LSD | P=0.0011 |
| **9F** | Time post-Drug | Three-way RM ANOVA F (2.785, 80.77) = 13.67 | P<0.0001 |
| | Vehicle (9-AC) vs. 9-AC | F (1, 29) = 251.4 | P<0.0001 |
| | Vehicle (Suzetrigine) vs. Suzetrigine | F (1, 29) = 2.175 | P=0.1102 |
| | Time post-drug x Vehicle (9-AC) vs. 9-AC | F (3, 87) = 13.33 | P<0.0001 |
| | Time post-drug x Vehicle (Suzetrigine) vs. Suzetrigine | F (3, 87) = 20.62 | P<0.0001 |
| | Vehicle (9-AC) vs. 9-AC vs Suzetrigine | F (1, 29) = 3.519 | P=0.0708 |
| | Time post-drug x Vehicle (9-AC) vs. Suzetrigine | F (3, 87) = 18.57 | P<0.0001 |
| | 0:Vehicle Vehicle vs. 0:Vehicle Suzetrigine | Tukey's multiple comparisons test | P=0.9997 |
| | 0:Vehicle Vehicle vs. 0:9-AC Vehicle | Tukey's multiple comparisons test | P=0.0004 |
| | 0:Vehicle Vehicle vs. 0:9-AC Suzetrigine | Tukey's multiple comparisons test | P<0.0001 |
| | 0:Vehicle Vehicle vs. 60:Vehicle Vehicle | Tukey's multiple comparisons test | P>0.9999 |
| | 0:Vehicle Vehicle vs. 60:Vehicle Suzetrigine | Tukey's multiple comparisons test | P>0.9999 |
| | 0:Vehicle Vehicle vs. 60:9-AC Vehicle | Tukey's multiple comparisons test | P=0.0010 |
| | 0:Vehicle Vehicle vs. 60:9-AC Suzetrigine | Tukey's multiple comparisons test | P=0.2024 |
| | 0:Vehicle Vehicle vs. 120:Vehicle Vehicle | Tukey's multiple comparisons test | P=0.9998 |
| | 0:Vehicle Vehicle vs. 120:Vehicle Suzetrigine | Tukey's multiple comparisons test | P=0.9542 |
| | 0:Vehicle Vehicle vs. 120:9-AC Vehicle | Tukey's multiple comparisons test | P<0.0001 |
| | 0:Vehicle Vehicle vs. 120:9-AC Suzetrigine | Tukey's multiple comparisons test | P=0.0260 |
| | 0:Vehicle Vehicle vs. 180:Vehicle Vehicle | Tukey's multiple comparisons test | P=0.9999 |
| | 0:Vehicle Vehicle vs. 180:Vehicle Suzetrigine | Tukey's multiple comparisons test | P>0.9999 |
| | 0:Vehicle Vehicle vs. 180:9-AC Vehicle | Tukey's multiple comparisons test | P<0.0001 |
| | 0:Vehicle Vehicle vs. 180:9-AC Suzetrigine | Tukey's multiple comparisons test | P=0.0047 |
| | 0:Vehicle Suzetrigine vs. 0:9-AC Vehicle | Tukey's multiple comparisons test | P=0.0003 |
| | 0:Vehicle Suzetrigine vs. 0:9-AC Suzetrigine | Tukey's multiple comparisons test | P<0.0001 |
| | 0:Vehicle Suzetrigine vs. 60:Vehicle Vehicle | Tukey's multiple comparisons test | P=0.9985 |
| | 0:Vehicle Suzetrigine vs. 60:Vehicle Suzetrigine | Tukey's multiple comparisons test | P=0.9995 |
| | 0:Vehicle Suzetrigine vs. 60:9-AC Vehicle | Tukey's multiple comparisons test | P=0.0010 |
| | 0:Vehicle Suzetrigine vs. 60:9-AC Suzetrigine | Tukey's multiple comparisons test | P=0.2784 |
| | 0:Vehicle Suzetrigine vs. 120:Vehicle Vehicle | Tukey's multiple comparisons test | P>0.9999 |
| | 0:Vehicle Suzetrigine vs. 120:Vehicle Suzetrigine | Tukey's multiple comparisons test | P>0.9999 |
| | 0:Vehicle Suzetrigine vs. 120:9-AC Vehicle | Tukey's multiple comparisons test | P<0.0001 |

| | | | |
|---|---|---|---|
| | 0:Vehicle Suzetrigine vs. 120:9-AC Suzetrigine | Tukey’s multiple comparisons test | P=0.0439 |
| | 0:Vehicle Suzetrigine vs. 180:Vehicle Vehicle | Tukey’s multiple comparisons test | P>0.9999 |
| | 0:Vehicle Suzetrigine vs. 180:Vehicle Suzetrigine | Tukey’s multiple comparisons test | P=0.9991 |
| | 0:Vehicle Suzetrigine vs. 180:9-AC Vehicle | Tukey’s multiple comparisons test | P<0.0001 |
| | 0:Vehicle Suzetrigine vs. 180:9-AC Suzetrigine | Tukey’s multiple comparisons test | P=0.0054 |
| | 0:9-AC Vehicle vs. 0:9-AC Suzetrigine | Tukey’s multiple comparisons test | P=0.1435 |
| | 0:9-AC Vehicle vs. 60:Vehicle Vehicle | Tukey’s multiple comparisons test | P=0.0004 |
| | 0:9-AC Vehicle vs. 60:Vehicle Suzetrigine | Tukey’s multiple comparisons test | P=0.0005 |
| | 0:9-AC Vehicle vs. 60:9-AC Vehicle | Tukey’s multiple comparisons test | P=0.9560 |
| | 0:9-AC Vehicle vs. 60:9-AC Suzetrigine | Tukey’s multiple comparisons test | P=0.6372 |
| | 0:9-AC Vehicle vs. 120:Vehicle Vehicle | Tukey’s multiple comparisons test | P=0.0004 |
| | 0:9-AC Vehicle vs. 120:Vehicle Suzetrigine | Tukey’s multiple comparisons test | P=0.0007 |
| | 0:9-AC Vehicle vs. 120:9-AC Vehicle | Tukey’s multiple comparisons test | P=0.9498 |
| | 0:9-AC Vehicle vs. 120:9-AC Suzetrigine | Tukey’s multiple comparisons test | P=0.0866 |
| | 0:9-AC Vehicle vs. 180:Vehicle Vehicle | Tukey’s multiple comparisons test | P=0.0005 |
| | 0:9-AC Vehicle vs. 180:Vehicle Suzetrigine | Tukey’s multiple comparisons test | P=0.0005 |
| | 0:9-AC Vehicle vs. 180:9-AC Vehicle | Tukey’s multiple comparisons test | P=0.9473 |
| | 0:9-AC Vehicle vs. 180:9-AC Suzetrigine | Tukey’s multiple comparisons test | P>0.9999 |
| | 0:9-AC Suzetrigine vs. 60:Vehicle Vehicle | Tukey’s multiple comparisons test | P<0.0001 |
| | 0:9-AC Suzetrigine vs. 60:Vehicle Suzetrigine | Tukey’s multiple comparisons test | P<0.0001 |
| | 0:9-AC Suzetrigine vs. 60:9-AC Vehicle | Tukey’s multiple comparisons test | P=0.4864 |
| | 0:9-AC Suzetrigine vs. 60:9-AC Suzetrigine | Tukey’s multiple comparisons test | * P=0.0220 |
| | 0:9-AC Suzetrigine vs. 120:Vehicle Vehicle | Tukey’s multiple comparisons test | P<0.0001 |
| | 0:9-AC Suzetrigine vs. 120:Vehicle Suzetrigine | Tukey’s multiple comparisons test | P=0.0001 |
| | 0:9-AC Suzetrigine vs. 120:9-AC Vehicle | Tukey’s multiple comparisons test | P=0.3157 |
| | 0:9-AC Suzetrigine vs. 120:9-AC Suzetrigine | Tukey’s multiple comparisons test | ** P=0.0013 |
| | 0:9-AC Suzetrigine vs. 180:Vehicle Vehicle | Tukey’s multiple comparisons test | P<0.0001 |
| | 0:9-AC Suzetrigine vs. 180:Vehicle Suzetrigine | Tukey’s multiple comparisons test | P<0.0001 |
| | 0:9-AC Suzetrigine vs. 180:9-AC Vehicle | Tukey’s multiple comparisons test | P=0.6104 |
| | 0:9-AC Suzetrigine vs. 180:9-AC Suzetrigine | Tukey’s multiple comparisons test | P=0.0998 |
| | 60:Vehicle Vehicle vs. 60:Vehicle Suzetrigine | Tukey’s multiple comparisons test | P>0.9999 |
| | 60:Vehicle Vehicle vs. 60:9-AC Vehicle | Tukey’s multiple comparisons test | P=0.0012 |
| | 60:Vehicle Vehicle vs. 60:9-AC Suzetrigine | Tukey’s multiple comparisons test | P=0.1972 |
| | 60:Vehicle Vehicle vs. 120:Vehicle Vehicle | Tukey’s multiple comparisons test | P=0.9998 |
| | 60:Vehicle Vehicle vs. 120:Vehicle Suzetrigine | Tukey’s multiple comparisons test | P=0.8018 |
| | 60:Vehicle Vehicle vs. 120:9-AC Vehicle | Tukey’s multiple comparisons test | P<0.0001 |
| | 60:Vehicle Vehicle vs. 120:9-AC Suzetrigine | Tukey’s multiple comparisons test | P=0.0260 |
| | 60:Vehicle Vehicle vs. 180:Vehicle Vehicle | Tukey’s multiple comparisons test | P=0.9980 |
| | 60:Vehicle Vehicle vs. 180:Vehicle Suzetrigine | Tukey’s multiple comparisons test | P>0.9999 |
| | 60:Vehicle Vehicle vs. 180:9-AC Vehicle | Tukey’s multiple comparisons test | P<0.0001 |
| | 60:Vehicle Vehicle vs. 180:9-AC Suzetrigine | Tukey’s multiple comparisons test | P=0.0049 |
| | 60:Vehicle Suzetrigine vs. 60:9-AC Vehicle | Tukey’s multiple comparisons test | P=0.0013 |
| | 60:Vehicle Suzetrigine vs. 60:9-AC Suzetrigine | Tukey’s multiple comparisons test | P=0.2039 |
| | 60:Vehicle Suzetrigine vs. 120:Vehicle Vehicle | Tukey’s multiple comparisons test | P>0.9999 |
| | 60:Vehicle Suzetrigine vs. 120:Vehicle Suzetrigine | Tukey’s multiple comparisons test | P=0.6311 |
| | 60:Vehicle Suzetrigine vs. 120:9-AC Vehicle | Tukey’s multiple comparisons test | P<0.0001 |
| | 60:Vehicle Suzetrigine vs. 120:9-AC Suzetrigine | Tukey’s multiple comparisons test | P=0.0285 |
| | 60:Vehicle Suzetrigine vs. 180:Vehicle Vehicle | Tukey’s multiple comparisons test | P=0.9995 |
| | 60:Vehicle Suzetrigine vs. 180:Vehicle Suzetrigine | Tukey’s multiple comparisons test | P>0.9999 |
| | 60:Vehicle Suzetrigine vs. 180:9-AC Vehicle | Tukey’s multiple comparisons test | P<0.0001 |

| | | | |
|---|---|---|---|
| | 60:Vehicle Suzetrigine vs. 180:9-AC Suzetrigine | Tukey’s multiple comparisons test | P=0.0054 |
| | 60:9-AC Vehicle vs. 60:9-AC Suzetrigine | Tukey’s multiple comparisons test | P=0.4705 |
| | 60:9-AC Vehicle vs. 120:Vehicle Vehicle | Tukey’s multiple comparisons test | P=0.0011 |
| | 60:9-AC Vehicle vs. 120:Vehicle Suzetrigine | Tukey’s multiple comparisons test | P=0.0016 |
| | 60:9-AC Vehicle vs. 120:9-AC Vehicle | Tukey’s multiple comparisons test | P>0.9999 |
| | 60:9-AC Vehicle vs. 120:9-AC Suzetrigine | Tukey’s multiple comparisons test | P=0.0717 |
| | 60:9-AC Vehicle vs. 180:Vehicle Vehicle | Tukey’s multiple comparisons test | P=0.0014 |
| | 60:9-AC Vehicle vs. 180:Vehicle Suzetrigine | Tukey’s multiple comparisons test | P=0.0013 |
| | 60:9-AC Vehicle vs. 180:9-AC Vehicle | Tukey’s multiple comparisons test | P>0.9999 |
| | 60:9-AC Vehicle vs. 180:9-AC Suzetrigine | Tukey’s multiple comparisons test | P=0.9999 |
| | 60:9-AC Suzetrigine vs. 120:Vehicle Vehicle | Tukey’s multiple comparisons test | P=0.2354 |
| | 60:9-AC Suzetrigine vs. 120:Vehicle Suzetrigine | Tukey’s multiple comparisons test | P=0.2693 |
| | 60:9-AC Suzetrigine vs. 120:9-AC Vehicle | Tukey’s multiple comparisons test | P=0.2837 |
| | 60:9-AC Suzetrigine vs. 120:9-AC Suzetrigine | Tukey’s multiple comparisons test | P>0.9999 |
| | 60:9-AC Suzetrigine vs. 180:Vehicle Vehicle | Tukey’s multiple comparisons test | P=0.2360 |
| | 60:9-AC Suzetrigine vs. 180:Vehicle Suzetrigine | Tukey’s multiple comparisons test | P=0.2101 |
| | 60:9-AC Suzetrigine vs. 180:9-AC Vehicle | Tukey’s multiple comparisons test | P=0.1997 |
| | 60:9-AC Suzetrigine vs. 180:9-AC Suzetrigine | Tukey’s multiple comparisons test | P=0.1927 |
| | 120:Vehicle Vehicle vs. 120:Vehicle Suzetrigine | Tukey’s multiple comparisons test | P>0.9999 |
| | 120:Vehicle Vehicle vs. 120:9-AC Vehicle | Tukey’s multiple comparisons test | P<0.0001 |
| | 120:Vehicle Vehicle vs. 120:9-AC Suzetrigine | Tukey’s multiple comparisons test | P=0.0331 |
| | 120:Vehicle Vehicle vs. 180:Vehicle Vehicle | Tukey’s multiple comparisons test | P>0.9999 |
| | 120:Vehicle Vehicle vs. 180:Vehicle Suzetrigine | Tukey’s multiple comparisons test | P>0.9999 |
| | 120:Vehicle Vehicle vs. 180:9-AC Vehicle | Tukey’s multiple comparisons test | P<0.0001 |
| | 120:Vehicle Vehicle vs. 180:9-AC Suzetrigine | Tukey’s multiple comparisons test | P=0.0052 |
| | 120:Vehicle Suzetrigine vs. 120:9-AC Vehicle | Tukey’s multiple comparisons test | P<0.0001 |
| | 120:Vehicle Suzetrigine vs. 120:9-AC Suzetrigine | Tukey’s multiple comparisons test | P=0.0457 |
| | 120:Vehicle Suzetrigine vs. 180:Vehicle Vehicle | Tukey’s multiple comparisons test | P=0.9998 |
| | 120:Vehicle Suzetrigine vs. 180:Vehicle Suzetrigine | Tukey’s multiple comparisons test | P=0.7940 |
| | 120:Vehicle Suzetrigine vs. 180:9-AC Vehicle | Tukey’s multiple comparisons test | P=0.0001 |
| | 120:Vehicle Suzetrigine vs. 180:9-AC Suzetrigine | Tukey’s multiple comparisons test | P=0.0070 |
| | 120:9-AC Vehicle vs. 120:9-AC Suzetrigine | Tukey’s multiple comparisons test | P=0.0084 |
| | 120:9-AC Vehicle vs. 180:Vehicle Vehicle | Tukey’s multiple comparisons test | P<0.0001 |
| | 120:9-AC Vehicle vs. 180:Vehicle Suzetrigine | Tukey’s multiple comparisons test | P<0.0001 |
| | 120:9-AC Vehicle vs. 180:9-AC Vehicle | Tukey’s multiple comparisons test | P>0.9999 |
| | 120:9-AC Vehicle vs. 180:9-AC Suzetrigine | Tukey’s multiple comparisons test | P=0.9959 |
| | 120:9-AC Suzetrigine vs. 180:Vehicle Vehicle | Tukey’s multiple comparisons test | P=0.0356 |
| | 120:9-AC Suzetrigine vs. 180:Vehicle Suzetrigine | Tukey’s multiple comparisons test | P=0.0297 |
| | 120:9-AC Suzetrigine vs. 180:9-AC Vehicle | Tukey’s multiple comparisons test | P=0.0060 |
| | 120:9-AC Suzetrigine vs. 180:9-AC Suzetrigine | Tukey’s multiple comparisons test | P=0.1241 |
| | 180:Vehicle Vehicle vs. 180:Vehicle Suzetrigine | Tukey’s multiple comparisons test | P>0.9999 |
| | 180:Vehicle Vehicle vs. 180:9-AC Vehicle | Tukey’s multiple comparisons test | P<0.0001 |
| | 180:Vehicle Vehicle vs. 180:9-AC Suzetrigine | Tukey’s multiple comparisons test | P=0.0060 |
| | 180:Vehicle Suzetrigine vs. 180:9-AC Vehicle | Tukey’s multiple comparisons test | P<0.0001 |
| | 180:Vehicle Suzetrigine vs. 180:9-AC Suzetrigine | Tukey’s multiple comparisons test | P=0.0055 |
| | 180:9-AC Vehicle vs. 180:9-AC Suzetrigine | Tukey’s multiple comparisons test | P=0.9638 |
| **9G** | Myotonia x Drug | RM Mixed-effects model | P<0.0001 |

| | | | |
|---|---|---|---|
| | Interaction | F (1, 15) = 176.3 | |
| | Myotonia (Vehicle or 9-AC) | F (1, 14) = 7.597 | P=0.0155 |
| | Drug (Vehicle or Suzetrigine) | F (1, 14) = 8.991 | P=0.0096 |
| | Vehicle: Vehicle vs. Suzetrigine | Uncorrected Fisher's LSD | P=0.8649 |
| | 9-AC: Vehicle vs. Suzetrigine | Uncorrected Fisher's LSD | ** P=0.0013 |
| | Vehicle: Vehicle vs. 9-AC | Uncorrected Fisher's LSD | P<0.0001 |
| | Suzetrigine: Vehicle vs. 9-AC | Uncorrected Fisher's LSD | P<0.0001 |
| **9H** | Time post-Drug | Three-way RM ANOVA<br>F (2.632, 76.33) = 1.036 | P=0.3749 |
| | Vehicle (9-AC) vs. 9-AC | F (1, 29) = 34.20 | P<0.0001 |
| | Vehicle (Suzetrigine) vs. Suzetrigine | F (1, 29) = 6.827 | P=0.0141 |
| | Time post-drug x Vehicle (9-AC) vs. 9-AC | F (3, 87) = 2.083 | P=0.1083 |
| | Time post-drug x Vehicle (Suzetrigine) vs. Suzetrigine | F (3, 87) = 1.073 | P=0.3647 |
| | Vehicle (9-AC) vs. 9-AC vs Suzetrigine | F (1, 29) = 0.005706 | P=0.9403 |
| | Time post-drug x Vehicle (9-AC) vs. Suzetrigine | F (3, 87) = 3.396 | P=0.0214 |
| | 0:Vehicle Vehicle vs. 0:Vehicle Suzetrigine | Tukey's multiple comparisons test | P=0.3209 |
| | 0:Vehicle Vehicle vs. 0:9-AC Vehicle | Tukey's multiple comparisons test | P=0.0157 |
| | 0:Vehicle Vehicle vs. 0:9-AC Suzetrigine | Tukey's multiple comparisons test | P=0.3914 |
| | 0:Vehicle Vehicle vs. 60:Vehicle Vehicle | Tukey's multiple comparisons test | P=0.9856 |
| | 0:Vehicle Vehicle vs. 60:Vehicle Suzetrigine | Tukey's multiple comparisons test | P>0.9999 |
| | 0:Vehicle Vehicle vs. 60:9-AC Vehicle | Tukey's multiple comparisons test | P=0.3871 |
| | 0:Vehicle Vehicle vs. 60:9-AC Suzetrigine | Tukey's multiple comparisons test | P>0.9999 |
| | 0:Vehicle Vehicle vs. 120:Vehicle Vehicle | Tukey's multiple comparisons test | P>0.9999 |
| | 0:Vehicle Vehicle vs. 120:Vehicle Suzetrigine | Tukey's multiple comparisons test | P=0.9199 |
| | 0:Vehicle Vehicle vs. 120:9-AC Vehicle | Tukey's multiple comparisons test | P=0.2808 |
| | 0:Vehicle Vehicle vs. 120:9-AC Suzetrigine | Tukey's multiple comparisons test | P>0.9999 |
| | 0:Vehicle Vehicle vs. 180:Vehicle Vehicle | Tukey's multiple comparisons test | P>0.9999 |
| | 0:Vehicle Vehicle vs. 180:Vehicle Suzetrigine | Tukey's multiple comparisons test | P=0.6678 |
| | 0:Vehicle Vehicle vs. 180:9-AC Vehicle | Tukey's multiple comparisons test | P=0.0039 |
| | 0:Vehicle Vehicle vs. 180:9-AC Suzetrigine | Tukey's multiple comparisons test | P=0.2503 |
| | 0:Vehicle Suzetrigine vs. 0:9-AC Vehicle | Tukey's multiple comparisons test | P=0.0007 |
| | 0:Vehicle Suzetrigine vs. 0:9-AC Suzetrigine | Tukey's multiple comparisons test | P=0.0479 |
| | 0:Vehicle Suzetrigine vs. 60:Vehicle Vehicle | Tukey's multiple comparisons test | P=0.9968 |
| | 0:Vehicle Suzetrigine vs. 60:Vehicle Suzetrigine | Tukey's multiple comparisons test | P=0.4561 |
| | 0:Vehicle Suzetrigine vs. 60:9-AC Vehicle | Tukey's multiple comparisons test | P=0.0235 |
| | 0:Vehicle Suzetrigine vs. 60:9-AC Suzetrigine | Tukey's multiple comparisons test | P=0.8984 |
| | 0:Vehicle Suzetrigine vs. 120:Vehicle Vehicle | Tukey's multiple comparisons test | P=0.3461 |
| | 0:Vehicle Suzetrigine vs. 120:Vehicle Suzetrigine | Tukey's multiple comparisons test | P>0.9999 |
| | 0:Vehicle Suzetrigine vs. 120:9-AC Vehicle | Tukey's multiple comparisons test | P=0.0179 |
| | 0:Vehicle Suzetrigine vs. 120:9-AC Suzetrigine | Tukey's multiple comparisons test | P=0.8435 |
| | 0:Vehicle Suzetrigine vs. 180:Vehicle Vehicle | Tukey's multiple comparisons test | P=0.5848 |
| | 0:Vehicle Suzetrigine vs. 180:Vehicle Suzetrigine | Tukey's multiple comparisons test | P>0.9999 |
| | 0:Vehicle Suzetrigine vs. 180:9-AC Vehicle | Tukey's multiple comparisons test | P=0.0003 |
| | 0:Vehicle Suzetrigine vs. 180:9-AC Suzetrigine | Tukey's multiple comparisons test | P=0.0106 |
| | 0:9-AC Vehicle vs. 0:9-AC Suzetrigine | Tukey's multiple comparisons test | P>0.9999 |
| | 0:9-AC Vehicle vs. 60:Vehicle Vehicle | Tukey's multiple comparisons test | P=0.0076 |
| | 0:9-AC Vehicle vs. 60:Vehicle Suzetrigine | Tukey's multiple comparisons test | P=0.3788 |
| | 0:9-AC Vehicle vs. 60:9-AC Vehicle | Tukey's multiple comparisons test | P>0.9999 |
| | 0:9-AC Vehicle vs. 60:9-AC Suzetrigine | Tukey's multiple comparisons test | P=0.8446 |
| | 0:9-AC Vehicle vs. 120:Vehicle Vehicle | Tukey's multiple comparisons test | P=0.0403 |
| | 0:9-AC Vehicle vs. 120:Vehicle Suzetrigine | Tukey's multiple comparisons test | P=0.0265 |
| | 0:9-AC Vehicle vs. 120:9-AC Vehicle | Tukey's multiple comparisons test | P>0.9999 |
| | 0:9-AC Vehicle vs. 120:9-AC Suzetrigine | Tukey's multiple comparisons test | P=0.4327 |

| | | | |
|---|---|---|---|
| | 0:9-AC Vehicle vs. 180:Vehicle Vehicle | Tukey's multiple comparisons test | P=0.5943 |
| | 0:9-AC Vehicle vs. 180:Vehicle Suzetrigine | Tukey's multiple comparisons test | P=0.0180 |
| | 0:9-AC Vehicle vs. 180:9-AC Vehicle | Tukey's multiple comparisons test | P>0.9999 |
| | 0:9-AC Vehicle vs. 180:9-AC Suzetrigine | Tukey's multiple comparisons test | P>0.9999 |
| | 0:9-AC Suzetrigine vs. 60:Vehicle Vehicle | Tukey's multiple comparisons test | P=0.1484 |
| | 0:9-AC Suzetrigine vs. 60:Vehicle Suzetrigine | Tukey's multiple comparisons test | P=0.5984 |
| | 0:9-AC Suzetrigine vs. 60:9-AC Vehicle | Tukey's multiple comparisons test | P>0.9999 |
| | 0:9-AC Suzetrigine vs. 60:9-AC Suzetrigine | Tukey's multiple comparisons test | P=0.4060 |
| | 0:9-AC Suzetrigine vs. 120:Vehicle Vehicle | Tukey's multiple comparisons test | P=0.4503 |
| | 0:9-AC Suzetrigine vs. 120:Vehicle Suzetrigine | Tukey's multiple comparisons test | P=0.1336 |
| | 0:9-AC Suzetrigine vs. 120:9-AC Vehicle | Tukey's multiple comparisons test | P>0.9999 |
| | 0:9-AC Suzetrigine vs. 120:9-AC Suzetrigine | Tukey's multiple comparisons test | P=0.4238 |
| | 0:9-AC Suzetrigine vs. 180:Vehicle Vehicle | Tukey's multiple comparisons test | P=0.8086 |
| | 0:9-AC Suzetrigine vs. 180:Vehicle Suzetrigine | Tukey's multiple comparisons test | P=0.0721 |
| | 0:9-AC Suzetrigine vs. 180:9-AC Vehicle | Tukey's multiple comparisons test | P>0.9999 |
| | 0:9-AC Suzetrigine vs. 180:9-AC Suzetrigine | Tukey's multiple comparisons test | P=0.9997 |
| | 60:Vehicle Vehicle vs. 60:Vehicle Suzetrigine | Tukey's multiple comparisons test | P>0.9999 |
| | 60:Vehicle Vehicle vs. 60:9-AC Vehicle | Tukey's multiple comparisons test | P=0.1152 |
| | 60:Vehicle Vehicle vs. 60:9-AC Suzetrigine | Tukey's multiple comparisons test | P=0.9981 |
| | 60:Vehicle Vehicle vs. 120:Vehicle Vehicle | Tukey's multiple comparisons test | P=0.9299 |
| | 60:Vehicle Vehicle vs. 120:Vehicle Suzetrigine | Tukey's multiple comparisons test | P>0.9999 |
| | 60:Vehicle Vehicle vs. 120:9-AC Vehicle | Tukey's multiple comparisons test | P=0.0818 |
| | 60:Vehicle Vehicle vs. 120:9-AC Suzetrigine | Tukey's multiple comparisons test | P=0.9987 |
| | 60:Vehicle Vehicle vs. 180:Vehicle Vehicle | Tukey's multiple comparisons test | P=0.6291 |
| | 60:Vehicle Vehicle vs. 180:Vehicle Suzetrigine | Tukey's multiple comparisons test | P=0.9987 |
| | 60:Vehicle Vehicle vs. 180:9-AC Vehicle | Tukey's multiple comparisons test | P=0.0031 |
| | 60:Vehicle Vehicle vs. 180:9-AC Suzetrigine | Tukey's multiple comparisons test | P=0.0647 |
| | 60:Vehicle Suzetrigine vs. 60:9-AC Vehicle | Tukey's multiple comparisons test | P=0.7239 |
| | 60:Vehicle Suzetrigine vs. 60:9-AC Suzetrigine | Tukey's multiple comparisons test | P>0.9999 |
| | 60:Vehicle Suzetrigine vs. 120:Vehicle Vehicle | Tukey's multiple comparisons test | P>0.9999 |
| | 60:Vehicle Suzetrigine vs. 120:Vehicle Suzetrigine | Tukey's multiple comparisons test | P=0.9996 |
| | 60:Vehicle Suzetrigine vs. 120:9-AC Vehicle | Tukey's multiple comparisons test | P=0.5886 |
| | 60:Vehicle Suzetrigine vs. 120:9-AC Suzetrigine | Tukey's multiple comparisons test | P>0.9999 |
| | 60:Vehicle Suzetrigine vs. 180:Vehicle Vehicle | Tukey's multiple comparisons test | P>0.9999 |
| | 60:Vehicle Suzetrigine vs. 180:Vehicle Suzetrigine | Tukey's multiple comparisons test | P=0.9851 |
| | 60:Vehicle Suzetrigine vs. 180:9-AC Vehicle | Tukey's multiple comparisons test | P=0.2397 |
| | 60:Vehicle Suzetrigine vs. 180:9-AC Suzetrigine | Tukey's multiple comparisons test | P=0.6414 |
| | 60:9-AC Vehicle vs. 60:9-AC Suzetrigine | Tukey's multiple comparisons test | P=0.9712 |
| | 60:9-AC Vehicle vs. 120:Vehicle Vehicle | Tukey's multiple comparisons test | P=0.4799 |
| | 60:9-AC Vehicle vs. 120:Vehicle Suzetrigine | Tukey's multiple comparisons test | P=0.1277 |
| | 60:9-AC Vehicle vs. 120:9-AC Vehicle | Tukey's multiple comparisons test | P>0.9999 |
| | 60:9-AC Vehicle vs. 120:9-AC Suzetrigine | Tukey's multiple comparisons test | P=0.7925 |
| | 60:9-AC Vehicle vs. 180:Vehicle Vehicle | Tukey's multiple comparisons test | P=0.9189 |
| | 60:9-AC Vehicle vs. 180:Vehicle Suzetrigine | Tukey's multiple comparisons test | P=0.0670 |
| | 60:9-AC Vehicle vs. 180:9-AC Vehicle | Tukey's multiple comparisons test | P=0.9997 |
| | 60:9-AC Vehicle vs. 180:9-AC Suzetrigine | Tukey's multiple comparisons test | P>0.9999 |
| | 60:9-AC Suzetrigine vs. 120:Vehicle Vehicle | Tukey's multiple comparisons test | P>0.9999 |
| | 60:9-AC Suzetrigine vs. 120:Vehicle Suzetrigine | Tukey's multiple comparisons test | P=0.9917 |
| | 60:9-AC Suzetrigine vs. 120:9-AC Vehicle | Tukey's multiple comparisons test | P=0.9241 |
| | 60:9-AC Suzetrigine vs. 120:9-AC Suzetrigine | Tukey's multiple comparisons test | P>0.9999 |
| | 60:9-AC Suzetrigine vs. 180:Vehicle Vehicle | Tukey's multiple comparisons test | P>0.9999 |

| | | | |
|---|---|---|---|
| | 60:9-AC Suzetrigine vs. 180:Vehicle Suzetrigine | Tukey's multiple comparisons test | P=0.9286 |
| | 60:9-AC Suzetrigine vs. 180:9-AC Vehicle | Tukey's multiple comparisons test | P=0.7062 |
| | 60:9-AC Suzetrigine vs. 180:9-AC Suzetrigine | Tukey's multiple comparisons test | P=0.6264 |
| | 120:Vehicle Vehicle vs. 120:Vehicle Suzetrigine | Tukey's multiple comparisons test | P=0.9151 |
| | 120:Vehicle Vehicle vs. 120:9-AC Vehicle | Tukey's multiple comparisons test | P=0.3503 |
| | 120:Vehicle Vehicle vs. 120:9-AC Suzetrigine | Tukey's multiple comparisons test | P>0.9999 |
| | 120:Vehicle Vehicle vs. 180:Vehicle Vehicle | Tukey's multiple comparisons test | P>0.9999 |
| | 120:Vehicle Vehicle vs. 180:Vehicle Suzetrigine | Tukey's multiple comparisons test | P=0.6630 |
| | 120:Vehicle Vehicle vs. 180:9-AC Vehicle | Tukey's multiple comparisons test | P=0.0124 |
| | 120:Vehicle Vehicle vs. 180:9-AC Suzetrigine | Tukey's multiple comparisons test | P=0.3356 |
| | 120:Vehicle Suzetrigine vs. 120:9-AC Vehicle | Tukey's multiple comparisons test | P=0.0886 |
| | 120:Vehicle Suzetrigine vs. 120:9-AC Suzetrigine | Tukey's multiple comparisons test | P=0.9924 |
| | 120:Vehicle Suzetrigine vs. 180:Vehicle Vehicle | Tukey's multiple comparisons test | P=0.9302 |
| | 120:Vehicle Suzetrigine vs. 180:Vehicle Suzetrigine | Tukey's multiple comparisons test | P>0.9999 |
| | 120:Vehicle Suzetrigine vs. 180:9-AC Vehicle | Tukey's multiple comparisons test | P=0.0139 |
| | 120:Vehicle Suzetrigine vs. 180:9-AC Suzetrigine | Tukey's multiple comparisons test | P=0.0855 |
| | 120:9-AC Vehicle vs. 120:9-AC Suzetrigine | Tukey's multiple comparisons test | P=0.6599 |
| | 120:9-AC Vehicle vs. 180:Vehicle Vehicle | Tukey's multiple comparisons test | P=0.8172 |
| | 120:9-AC Vehicle vs. 180:Vehicle Suzetrigine | Tukey's multiple comparisons test | P=0.0466 |
| | 120:9-AC Vehicle vs. 180:9-AC Vehicle | Tukey's multiple comparisons test | P>0.9999 |
| | 120:9-AC Vehicle vs. 180:9-AC Suzetrigine | Tukey's multiple comparisons test | P>0.9999 |
| | 120:9-AC Suzetrigine vs. 180:Vehicle Vehicle | Tukey's multiple comparisons test | P>0.9999 |
| | 120:9-AC Suzetrigine vs. 180:Vehicle Suzetrigine | Tukey's multiple comparisons test | P=0.9101 |
| | 120:9-AC Suzetrigine vs. 180:9-AC Vehicle | Tukey's multiple comparisons test | P=0.2763 |
| | 120:9-AC Suzetrigine vs. 180:9-AC Suzetrigine | Tukey's multiple comparisons test | P=0.1121 |
| | 180:Vehicle Vehicle vs. 180:Vehicle Suzetrigine | Tukey's multiple comparisons test | P=0.7294 |
| | 180:Vehicle Vehicle vs. 180:9-AC Vehicle | Tukey's multiple comparisons test | P=0.3914 |
| | 180:Vehicle Vehicle vs. 180:9-AC Suzetrigine | Tukey's multiple comparisons test | P=0.8672 |
| | 180:Vehicle Suzetrigine vs. 180:9-AC Vehicle | Tukey's multiple comparisons test | P=0.0103 |
| | 180:Vehicle Suzetrigine vs. 180:9-AC Suzetrigine | Tukey's multiple comparisons test | P=0.0459 |
| | 180:9-AC Vehicle vs. 180:9-AC Suzetrigine | Tukey's multiple comparisons test | P=>0.9999 |
| **9I** | Myotonia x Drug Interaction | RM Mixed-effects model F (1, 15) = 22.49 | P=0.0003 |
| | Myotonia (Vehicle or 9-AC) | F (1, 14) = 7.286 | P=0.0173 |
| | Drug (Vehicle or Suzetrigine) | F (1, 14) = 0.3848 | P=0.5450 |
| | Vehicle: Vehicle vs. Suzetrigine | Uncorrected Fisher's LSD | P=0.1587 |
| | 9-AC: Vehicle vs. Suzetrigine | Uncorrected Fisher's LSD | * P=0.0361 |
| | Vehicle: Vehicle vs. 9-AC | Uncorrected Fisher's LSD | P=0.0005 |
| | Suzetrigine: Vehicle vs. 9-AC | Uncorrected Fisher's LSD | P=0.0037 |
| **10B** | *Clcn1*$^{+/+}$ vs. *Clcn1*$^{adr/+}$ vs. *Clcn1*$^{adr/adr}$ | One-way ANOVA F (2, 26) = 15.80 | P<0.0001 |
| | *Clcn1*$^{+/+}$ vs. *Clcn1*$^{adr/+}$ | Tukey's multiple comparisons test | P=0.9953 |
| | *Clcn1*$^{+/+}$ vs. *Clcn1*$^{adr/adr}$ | Tukey's multiple comparisons test | *** P=0.0001 |
| | *Clcn1*$^{adr/+}$ vs. *Clcn1*$^{adr/adr}$ | Tukey's multiple comparisons test | *** P=0.0002 |
| **10C** | *Clcn1*$^{+/+}$ vs. *Clcn1*$^{adr/+}$ vs. *Clcn1*$^{adr/adr}$ | One-way ANOVA F (2, 26) = 15.45 | P<0.0001 |
| | *Clcn1*$^{+/+}$ vs. *Clcn1*$^{adr/+}$ | Tukey's multiple comparisons test | P=0.9778 |
| | *Clcn1*$^{+/+}$ vs. *Clcn1*$^{adr/adr}$ | Tukey's multiple comparisons test | *** P=0.0001 |
| | *Clcn1*$^{adr/+}$ vs. *Clcn1*$^{adr/adr}$ | Tukey's multiple comparisons test | *** P=0.0004 |

| | | | |
|---|---|---|---|
| **10D** | *Clcn1*$^{+/+}$ vs. *Clcn1*$^{adr/+}$ vs. *Clcn1*$^{adr/adr}$ | One-way ANOVA<br>F (2, 26) = 68.90 | P<0.0001 |
| | *Clcn1*$^{+/+}$ vs. *Clcn1*$^{adr/+}$ | Tukey's multiple comparisons test | P=0.9826 |
| | *Clcn1*$^{+/+}$ vs. *Clcn1*$^{adr/adr}$ | Tukey's multiple comparisons test | $^{****}$ P<0.0001 |
| | *Clcn1*$^{adr/+}$ vs. *Clcn1*$^{adr/adr}$ | Tukey's multiple comparisons test | $^{****}$ P<0.0001 |
| **10E** | *Clcn1*$^{+/+}$ vs. *Clcn1*$^{adr/+}$ vs. *Clcn1*$^{adr/adr}$ | One-way ANOVA<br>F (2, 26) = 13.77 | P<0.0001 |
| | *Clcn1*$^{+/+}$ vs. *Clcn1*$^{adr/+}$ | Tukey's multiple comparisons test | P=0.4861 |
| | *Clcn1*$^{+/+}$ vs. *Clcn1*$^{adr/adr}$ | Tukey's multiple comparisons test | $^{****}$ P<0.0001 |
| | *Clcn1*$^{adr/+}$ vs. *Clcn1*$^{adr/adr}$ | Tukey's multiple comparisons test | $^{**}$ P=0.0041 |
| **10F** | *Clcn1*$^{+/+}$ vs. *Clcn1*$^{adr/+}$ vs. *Clcn1*$^{adr/adr}$ | One-way ANOVA<br>F (2, 26) = 34.79 | P<0.0001 |
| | *Clcn1*$^{+/+}$ vs. *Clcn1*$^{adr/+}$ | Tukey's multiple comparisons test | P=0.9569 |
| | *Clcn1*$^{+/+}$ vs. *Clcn1*$^{adr/adr}$ | Tukey's multiple comparisons test | $^{****}$ P<0.0001 |
| | *Clcn1*$^{adr/+}$ vs. *Clcn1*$^{adr/adr}$ | Tukey's multiple comparisons test | $^{****}$ P<0.0001 |
| **11C** | Genotype x Drug<br>Interaction | Two-way ANOVA<br>F (1, 65) = 0.01628 | P=0.8989 |
| | Genotype (*Clcn1*$^{+/+}$ or *Clcn1*$^{adr/adr}$) | F (1, 65) = 10.72 | P=0.0017 |
| | Drug (Vehicle or Suzetrigine) | F (1, 65) = 7.194 | P=0.0093 |
| **11D** | Genotype x Drug<br>Interaction | Two-way ANOVA<br>F (1, 65) = 1.353 | P=0.2490 |
| | Genotype (*Clcn1*$^{+/+}$ or *Clcn1*$^{adr/adr}$) | F (1, 65) = 0.5310 | P=0.4688 |
| | Drug (Vehicle or Suzetrigine) | F (1, 65) = 11.98 | P=0.0010 |
| **11E** | Genotype x Drug<br>Interaction | Two-way ANOVA<br>F (1, 65) = 0.2411 | P=0.6251 |
| | Genotype (*Clcn1*$^{+/+}$ or *Clcn1*$^{adr/adr}$) | F (1, 65) = 2.101 | P=0.1520 |
| | Drug (Vehicle or Suzetrigine) | F (1, 65) = 20.68 | P<0.0001 |
| **11G** | Genotype x Drug<br>Interaction | Two-way ANOVA<br>F (1, 65) = 0.06418 | P=0.8008 |
| | Genotype (*Clcn1*$^{+/+}$ or *Clcn1*$^{adr/adr}$) | F (1, 65) = 0.07793 | P=0.7810 |
| | Drug (Vehicle or Suzetrigine) | F (1, 65) = 59.35 | P<0.0001 |
| **11H** | Genotype x Drug<br>Interaction | Two-way ANOVA<br>F (1, 65) = 0.3984 | P=0.5301 |
| | Genotype (*Clcn1*$^{+/+}$ or *Clcn1*$^{adr/adr}$) | F (1, 65) = 0.9902 | P=0.3234 |
| | Drug (Vehicle or Suzetrigine) | F (1, 65) = 28.62 | P<0.0001 |
| **11I** | Genotype x Drug<br>Interaction | Two-way ANOVA<br>F (1, 65) = 0.2062 | P=0.6513 |
| | Genotype (*Clcn1*$^{+/+}$ or *Clcn1*$^{adr/adr}$) | F (1, 65) = 4.231 | P=0.0437 |
| | Drug (Vehicle or Suzetrigine) | F (1, 65) = 0.6202 | P=0.4338 |
| **11J** | Genotype x Drug<br>Interaction | Two-way ANOVA<br>F (1, 65) = 0.6007 | P=0.4411 |
| | Genotype (*Clcn1*$^{+/+}$ or *Clcn1*$^{adr/adr}$) | F (1, 65) = 0.2647 | P=0.6087 |
| | Drug (Vehicle or Suzetrigine) | F (1, 65) = 44.91 | P<0.0001 |
| **11K** | Genotype x Drug<br>Interaction | Two-way ANOVA<br>F (1, 65) = 1.085 | P=0.3015 |
| | Genotype (*Clcn1*$^{+/+}$ or *Clcn1*$^{adr/adr}$) | F (1, 65) = 2.190 | P=0.1438 |
| | Drug (Vehicle or Suzetrigine) | F (1, 65) = 1.430 | P=0.2360 |
| **11L** | Genotype x Drug<br>Interaction | Two-way ANOVA<br>F (1, 65) = 0.4791 | P=0.4913 |
| | Genotype (*Clcn1*$^{+/+}$ or *Clcn1*$^{adr/adr}$) | F (1, 65) = 0.03003 | P=0.8630 |
| | Drug (Vehicle or Suzetrigine) | F (1, 65) = 15.81 | P=0.0002 |
| **11M** | Genotype x Drug<br>Interaction | Two-way ANOVA<br>F (1, 65) = 1.300 | P=0.2585 |
| | Genotype (*Clcn1*$^{+/+}$ or *Clcn1*$^{adr/adr}$) | F (1, 65) = 1.266 | P=0.2647 |
| | Drug (Vehicle or Suzetrigine) | F (1, 65) = 11.63 | P=0.0011 |
| **11N** | Genotype x Drug<br>Interaction | Two-way ANOVA<br>F (1, 65) = 0.2482 | P=0.6200 |
| | Genotype (*Clcn1*$^{+/+}$ or *Clcn1*$^{adr/adr}$) | F (1, 65) = 4.705 | P=0.0337 |
| | Drug (Vehicle or Suzetrigine) | F (1, 65) = 5.535 | P=0.0217 |
| **12B** | Genotype x Drug | Two-way RM ANOVA | P=0.0018 |

| | | | |
|---|---|---|---|
| | Interaction | F (1, 19) = 13.22 | |
| | Genotype (*Clcn1*$^{+/+}$ or *Clcn1*$^{adr/adr}$) | F (1, 19) = 14.84 | P=0.0011 |
| | Drug (Vehicle or Suzetrigine) | F (1, 19) = 25.66 | P<0.0001 |
| | *Clcn1*$^{+/+}$: Vehicle vs. Suzetrigine | Holm-Šidák's multiple comparisons test | P=0.3358 |
| | *Clcn1*$^{adr/adr}$: Vehicle vs. Suzetrigine | Holm-Šidák's multiple comparisons test | $^{****}$ P<0.0001 |
| **12C** | Genotype x Drug Interaction | Two-way RM ANOVA F (1, 19) = 84.95 | P<0.0001 |
| | Genotype (*Clcn1*$^{+/+}$ or *Clcn1*$^{adr/adr}$) | F (1, 19) = 20.81 | P=0.0002 |
| | Drug (Vehicle or Suzetrigine) | F (1, 19) = 84.95 | P<0.0001 |
| | *Clcn1*$^{+/+}$: Vehicle vs. Suzetrigine | Holm-Šidák's multiple comparisons test | P>0.999 |
| | *Clcn1*$^{adr/adr}$: Vehicle vs. Suzetrigine | Holm-Šidák's multiple comparisons test | $^{****}$ P<0.0001 |
| **12D** | Genotype x Drug Interaction | Two-way RM ANOVA F (1, 19) = 49.78 | P<0.0001 |
| | Genotype (*Clcn1*$^{+/+}$ or *Clcn1*$^{adr/adr}$) | F (1, 19) = 198.9 | P<0.0001 |
| | Drug (Vehicle or Suzetrigine) | F (1, 19) = 54.09 | P<0.0001 |
| | *Clcn1*$^{+/+}$: Vehicle vs. Suzetrigine | Holm-Šidák's multiple comparisons test | P=0.8385 |
| | *Clcn1*$^{adr/adr}$: Vehicle vs. Suzetrigine | Holm-Šidák's multiple comparisons test | $^{****}$ P<0.0001 |
| **12E** | Genotype x Drug Interaction | Two-way RM ANOVA F (1, 19) = 31.15 | P<0.0001 |
| | Genotype (*Clcn1*$^{+/+}$ or *Clcn1*$^{adr/adr}$) | F (1, 19) = 6.366 | P=0.0207 |
| | Drug (Vehicle or Suzetrigine) | F (1, 19) = 2.475 | P=0.1322 |
| | *Clcn1*$^{+/+}$: Vehicle vs. Suzetrigine | Holm-Šidák's multiple comparisons test | $^{*}$ P=0.0122 |
| | *Clcn1*$^{adr/adr}$: Vehicle vs. Suzetrigine | Holm-Šidák's multiple comparisons test | $^{***}$ P=0.0001 |
| **12F** | Genotype x Drug Interaction | Two-way RM ANOVA F (1, 14) = 13.46 | P=0.0025 |
| | Genotype (*Clcn1*$^{+/+}$ or *Clcn1*$^{adr/adr}$) | F (1, 14) = 34.29 | P<0.0001 |
| | Drug (Vehicle or Suzetrigine) | F (1, 14) = 26.38 | P=0.0002 |
| | *Clcn1*$^{+/+}$: Vehicle vs. Suzetrigine | Holm-Šidák's multiple comparisons test | P=0.3170 |
| | *Clcn1*$^{adr/adr}$: Vehicle vs. Suzetrigine | Holm-Šidák's multiple comparisons test | $^{****}$ P<0.0001 |
| **S1C** | Vehicle vs. 9-AC | Unpaired Two-tailed t test with Welch's correction | P=0.7687 |
| **S1D** | Vehicle vs. 9-AC | Unpaired Two-tailed t test with Welch's correction | P=0.5552 |
| **S1E** | Vehicle vs. 9-AC | Unpaired Two-tailed t test with Welch's correction | P=0.2729 |
| **S2C** | Vehicle vs. 9-AC | Unpaired Two-tailed t test with Welch's correction | P=0.6062 |
| **S2D** | Vehicle vs. 9-AC | Unpaired Two-tailed t test with Welch's correction | P=0.1397 |
| **S3B** | Vehicle vs. 9-AC | Unpaired Two-tailed t test with Welch's correction | NS P=0.0502 |
| **S3C** | Vehicle vs. 9-AC | Unpaired Two-tailed t test with Welch's correction | P=0.7355 |
| **S3D** | Vehicle vs. 9-AC | Unpaired Two-tailed t test with Welch's correction | P=0.8971 |
| **S4C** | Current Step x Drug Interaction | Two-way RM ANOVA F (5, 155) = 0.7170 | P=0.6116 |
| | Current Step | F (1.925, 59.67) = 17.41 | P<0.0001 |
| | Drug (Vehicle vs. 9-AC) | F (1, 31) = 2.543 | P=0.1209 |
| **S4D** | Vehicle vs. 9-AC | Contingency: Fisher's exact test | P=0.2073 |
| **S4E** | Vehicle vs. 9-AC | Unpaired Two-tailed t test with Welch's correction | P=0.5961 |
| **S4F** | Vehicle vs. 9-AC | Unpaired Two-tailed t test with Welch's correction | P=0.2791 |
| **S4G** | Vehicle vs. 9-AC | Unpaired Two-tailed t test with Welch's correction | P=0.2216 |
| **S4H** | Vehicle vs. 9-AC | Unpaired Two-tailed t test with Welch's correction | NS P=0.0649 |
| **S4I** | Vehicle vs. 9-AC | Unpaired Two-tailed t test with Welch's correction | NS P=0.0979 |

| | | | |
|---|---|---|---|
| **S5B** | Current Step x Drug Interaction | Two-way RM ANOVA<br>F (5, 60) = 0.8498 | P=0.5201 |
| | Current Step | F (1.555, 18.66) = 14.10 | P=0.0004 |
| | Drug (Vehicle vs. 9-AC) | F (1, 12) = 4.761 | P=0.0497 |
| **S5C** | Vehicle vs. 9-AC | Unpaired Two-tailed t test with Welch's correction | P=0.2300 |
| **S5D** | Vehicle vs. 9-AC | Unpaired Two-tailed t test with Welch's correction | $^{*}$ P=0.0427 |
| **S5E** | Vehicle vs. 9-AC | Unpaired Two-tailed t test with Welch's correction | $^{**}$ P=0.0013 |
| **S5F** | Vehicle vs. 9-AC | Unpaired Two-tailed t test with Welch's correction | NS P=0.0658 |
| **S5G** | Vehicle vs. 9-AC | Unpaired Two-tailed t test with Welch's correction | $^{*}$ P=0.0166 |
| **S5I** | Current Step x Drug Interaction | Two-way RM ANOVA<br>F (3, 15) = 0.9719 | P=0.4319 |
| | Current Step | F (1.446, 7.231) = 44.99 | P=0.0001 |
| | Drug (Vehicle vs. 9-AC) | F (1, 5) = 1.966 | P=0.2198 |
| **S5J** | Vehicle vs. 9-AC | Unpaired Two-tailed t test with Welch's correction | P=0.7997 |
| **S5K** | Vehicle vs. 9-AC | Unpaired Two-tailed t test with Welch's correction | P=0.6428 |
| **S5L** | Vehicle vs. 9-AC | Unpaired Two-tailed t test with Welch's correction | P=0.4908 |
| **S5M** | Vehicle vs. 9-AC | Unpaired Two-tailed t test with Welch's correction | P=0.7022 |
| **S5N** | Vehicle vs. 9-AC | Unpaired Two-tailed t test with Welch's correction | P=0.7483 |
| **S5P** | Current Step x Drug Interaction | Two-way RM ANOVA<br>F (5, 25) = 0.5273 | P=0.7534 |
| | Current Step | F (1.208, 6.042) = 1.799 | P=0.2338 |
| | Drug (Vehicle vs. 9-AC) | F (1, 5) = 0.03673 | P=0.8556 |
| **S5Q** | Vehicle vs. 9-AC | Unpaired Two-tailed t test with Welch's correction | P=0.5394 |
| **S5R** | Vehicle vs. 9-AC | Unpaired Two-tailed t test with Welch's correction | P=0.9688 |
| **S5S** | Vehicle vs. 9-AC | Unpaired Two-tailed t test with Welch's correction | P=0.4070 |
| **S5T** | Vehicle vs. 9-AC | Unpaired Two-tailed t test with Welch's correction | P=0.7281 |
| **S5U** | Vehicle vs. 9-AC | Unpaired Two-tailed t test with Welch's correction | P=0.6955 |
| **S6A** | *Clcn1*$^{+/+}$ vs. *Clcn1*$^{adr/+}$ vs. *Clcn1*$^{adr/adr}$ | One-way ANOVA<br>F (2, 23) = 60.93 | P<0.0001 |
| | *Clcn1*$^{+/+}$ vs. *Clcn1*$^{adr/+}$ | Tukey's multiple comparisons test | P=0.5438 |
| | *Clcn1*$^{+/+}$ vs. *Clcn1*$^{adr/adr}$ | Tukey's multiple comparisons test | $^{****}$ P<0.0001 |
| | *Clcn1*$^{adr/+}$ vs. *Clcn1*$^{adr/adr}$ | Tukey's multiple comparisons test | $^{****}$ P<0.0001 |
| **S6B** | *Clcn1*$^{+/+}$ vs. *Clcn1*$^{adr/+}$ vs. *Clcn1*$^{adr/adr}$ | One-way ANOVA<br>F (2, 23) = 129.1 | P<0.0001 |
| | *Clcn1*$^{+/+}$ vs. *Clcn1*$^{adr/+}$ | Tukey's multiple comparisons test | P=0.5392 |
| | *Clcn1*$^{+/+}$ vs. *Clcn1*$^{adr/adr}$ | Tukey's multiple comparisons test | $^{****}$ P<0.0001 |
| | *Clcn1*$^{adr/+}$ vs. *Clcn1*$^{adr/adr}$ | Tukey's multiple comparisons test | $^{****}$ P<0.0001 |
| **S6C** | *Clcn1*$^{+/+}$ vs. *Clcn1*$^{adr/+}$ vs. *Clcn1*$^{adr/adr}$ | Kruskal-Wallis Test<br>F (2, 23) = 129.1 | P<0.0001 |
| | *Clcn1*$^{+/+}$ vs. *Clcn1*$^{adr/+}$ | Dunn's multiple comparisons test | P>0.9999 |
| | *Clcn1*$^{+/+}$ vs. *Clcn1*$^{adr/adr}$ | Dunn's multiple comparisons test | $^{****}$ P<0.0001 |
| | *Clcn1*$^{adr/+}$ vs. *Clcn1*$^{adr/adr}$ | Dunn's multiple comparisons test | $^{***}$ P=0.0001 |

**Supplemental Statistics Table**: $^{*}$P<0.05, $^{**}$P<0.01, $^{***}$P<0.001, $^{****}$P<0.0001

| | *Clcn1*$^{+/+}$ | *Clcn1*$^{adr/adr}$ | P Value |
|---|---|---|---|
| RMP (mV) | -43.24 ± 0.511 | -41.50 ± 0.3807 | * P=0.012 |
| Rheobase (pA) | 46.61 ± 7.466 | 43.41 ± 5.913 | P=0.9271 |
| # of Action Potentials | 16.47 ±1.627 | 17.83 ± 1.237 | P=0.4253 |
| Amplitude (mV) | 63.36 ± 3.619 | 63.27 ± 2.559 | P=0.7886 |
| Half Width (ms) | 2.576 ± 0.3038 | 3.225 ± 0.2475 | * P=0.0325 |
| Threshold (mV) | -10.51 ± 1.363 | -8.255 ± 1.107 | P=0.2611 |
| Rise Time (ms) | 1.547 ± 0.1641 | 1.642 ± 0.1108 | P=0.5329 |
| Decay Time (ms) | 3.447 ± 0.4339 | 4.521 ± 0.3839 | * P=0.0293 |
| mAHP (mV) | -50.11 ± 1.883 | -48.24 ± 1.877 | P=0.4125 |
| Spike Time (ms) | 184.8 ± 27.90 | 184.4 ± 25.88 | P=0.9218 |
| Cell Size (pF) | 13.17 ± 0.8138 | 11.94 ± 0.4282 | P=0.2725 |
| Input Resistance | 464.2 ± 29.08 | 423.5 ± 30.92 | P=0.3434 |

**Supplemental Table 3: Comparison of excitability metrics between vehicle-treated neurons from *Clcn1*$^{+/+}$ and *Clcn1*$^{adr/adr}$ mice from Figure 11.** Data were collected from n=17 *Clcn1*$^{+/+}$ neurons and n=24 *Clcn1*$^{adr/adr}$ neurons. All comparisons were made with two-tailed Mann-Whitney tests. Values represent means ± SEM. *$p < 0.05$.
